# Single-cell resolved differentiation of pre-Kranz anatomy in maize leaf primordia

**DOI:** 10.1101/2024.07.10.602848

**Authors:** Juan Yi, Hong Su, Shilong Tian, Olga Sedelnikova, Yonghe Chen, Caiyao Zhao, Jianzhao Yang, Yijing Zhang, Xin-Guang Zhu, Jane A. Langdale, Jia-Wei Wang, Peng Wang

## Abstract

Typical C_4_ plants such as maize possess highly optimized Kranz-type leaf anatomy, whereby concentric wreaths of mesophyll and bundle sheath cells surround closely spaced veins. The veins and the cells that surround them are derived from the middle ground meristem (mGM) through processes that are as yet undefined. Here we distinguished the active zone of vascular development within early leaf primordia, and used comparative transcriptomics of sub-sectioned maize and rice primordia to identify cohorts of genes likely involved in early Kranz development. Leveraging single-nucleus RNA sequencing (snRNA-seq) we then explored the cell heterogeneity and developmental trajectories within single maize leaf primordia. Assisted by *in situ* hybridization, cell clusters of mGM and procambium were identified, with candidate marker genes showing different yet inter-related expression patterns. Localization of the vascular marker *ZmSHR1* was preceded by that of *ZmEREB161* and *ZmEREB114* in terms of procambium initiation. Potential subclusters of bundle sheath cells and different layer of mesophyll cells were depicted from developing cells toward the tip of sub-sectioned maize primordia. Collectively our results identify potential mGM derived or procambium localized Kranz regulators and provide resources for investigating leaf vein development in maize and rice, at sub-primordium and single-cell resolution.

## Introduction

Photosynthesis is an important part of the carbon cycle in nature, providing organic matter and oxygen to the entire biosphere. Plant photosynthesis occurs via C_3_, C_4_, or CAM pathways. In C_3_ photosynthesis, CO_2_ enters mesophyll (M) cells and is assimilated into carbohydrates by Ribulose-1,5-bisphosphate carboxylase/oxygenase (Rubisco) (Bassham, 2003). In C_4_ photosynthesis, a CO_2_ concentrating mechanism transfers C_4_ acids from M cells into bundle sheath (BS) cells where Rubisco is specifically enriched, such that CO_2_ assimilation operates more efficiently, with greatly improved radiation, water and nitrogen use efficiencies (Hatch and Slack, 1998; Ghannoum et al., 2010). Although C_4_ plants account for only 3% of land plant species, they account for 25% of terrestrial primary productivity (Edwards and Still, 2008). This high-efficiency photosynthetic mechanism is closely related to the unique Kranz anatomy of C_4_ leaves (Furbank, 2017; von Caemmerer et al., 2017).

The typical Kranz anatomy of C_4_ maize is exquisitely organized, with veins (V) surrounded by a ring of chloroplast-rich BS cells which are themselves surrounded by a layer of M cells, with many plasmodesmata connecting the two types of cells (Brown et al., 1975). Only two M cells separate adjacent veins, leading to a continuous V-BS-M-M-BS-V distribution across the medio-lateral leaf axis which maximizes the BS:M ratio to support the CO_2_ concentration mechanism (Fouracre et al., 2014). With a view to understanding how Kranz anatomy develops, comparative studies have been carried out in a number of different contexts, which generally included three aspects of Kranz development: (i) initiation of procambium (Wang et al., 2013; Liu et al., 2013; Liu et al., 2022; Robil and McSteen, 2023); (ii) BS and M-cell specification (Li et al., 2010; Chang et al., 2012; Aubry et al., 2014; John et al., 2014; Tausta et al., 2014; Hendron et al., 2020; Singh et al., 2020; Bezrutczyk et al., 2021); and (iii) integration of the C_4_ cycle (Brautigam et al., 2011; Furumoto et al., 2011; Gowik et al., 2011; Christin et al., 2013; Brautigam et al., 2014; Kulahoglu et al., 2014; Mallmann et al., 2014; Wang et al., 2014; Ding et al., 2015; Covshoff et al., 2016; Arrivault et al., 2019). Procambium initiation and BS/M-cell specification occurs early in maize leaf development (Wang et al., 2013), and little is known about the regulators of these early processes. Our previous study comparing transcriptional profiles of maize foliar and husk (with and without Kranz anatomy) leaf primordia proposed potential regulators including the SHOOT ROOT (SHR) / SCARECROW (SCR) regulatory module (Wang et al., 2013; Fouracre et al., 2014). Cell type-specific transcriptomes of early precursors of maize BS+V and M cells using laser capture microdissection (LCM) further identified genes encoding auxin transporters (PIN1a, PIN1d, and LAX2) and transcription factors (TFs) that are expressed during early Kranz development (Liu et al., 2022). Recent work indicated that a SHR-INDETERMINATE DOMAIN (IDD) regulatory circuit mediates auxin transport by negatively regulating PIN-FORMED (PIN) expression to modulate minor vein patterning in leaves of both C_3_ and C_4_ grasses (Liu et al., 2023); while the combined action of SCR and NAKED ENDOSPERM (NKD) IDD controls the number of mesophyll cells specified between veins in the leaves of C_4_ but not C_3_ grasses (Hughes et al., 2023). Fluorescent protein reporters mapping auxin, cytokinin, and gibberellic acid response patterns in maize leaf primordia further defined the roles of these hormones in medial-lateral growth and vein formation (Robil and McSteen, 2023). Despite these advances critical events in the development of Kranz anatomy remain to be uncovered.

Given that Kranz patterning begins with the specification of individual procambial initials from within the middle ground meristem layer (mGM) (Fouracre et al., 2014; Liu et al., 2022), tissue-level approaches are likely to miss important steps in the developmental trajectory. To address this deficit, we have investigated the early stages of Kranz development at the single cell level. We first identified and separated the active and inactive domains of vein formation within maize leaf primordia, and then performed comparative transcriptomic analysis with corresponding rice tissues. We then used single-nucleus RNA sequencing (snRNA-seq) to investigate the critical developmental period within C_4_ leaf primordia. These approaches together revealed cell clusters specific to the initiating procambium and identified gene expression profiles and developmental trajectories of essential cell types during Kranz development.

## Results

### Identification of regions within single maize leaf primordia where intermediate veins are being actively initiated

Previous studies on vein development in maize suggested that the midvein and lateral veins develop/extend from the base toward the tip of leaf primordium (acropetally), while the secondary/intermediate veins subsequently start and extend from the tip toward the base (basipetally) (Sharman,1942). To identify regions where veins are being actively initiated within the developing leaf, maize *pZmPIN1a::ZmPIN1a:YFP* reporter lines were examined. In plastochron 3 (P3) and P4 primordium, intermediate veins had been initiated but many had not extended into the base region **(Figure 1A; Supplemental Figure 1)**. Flattened P4 primordium of ∼1.5 mm exhibited a wide conical shape, and veins were distributed from the midvein towards both margins, forming a distinguished spindle-shaped region. As the P4 primordium elongated longitudinally to ∼2.5 mm, both the number of veins and the leaf width increased, with new veins initiated between the lateral veins. Between ∼2.5 and ∼5 mm, the leaf tip lengthened considerably, vein density increased significantly, and the spindle area elongated (schematically illustrated in **Figure 1B** and representative cross sections shown in **Figure 1C**). At P5 the primordium was much larger in size and the development of intermediate veins was more advanced. In P5 primordia of 10 mm, the spindle shaped region was elongated, veins appeared to be near parallel, and the leaf shape was close to that of mature leaves **(Supplemental Figure 1)**.

**Figure 1.**
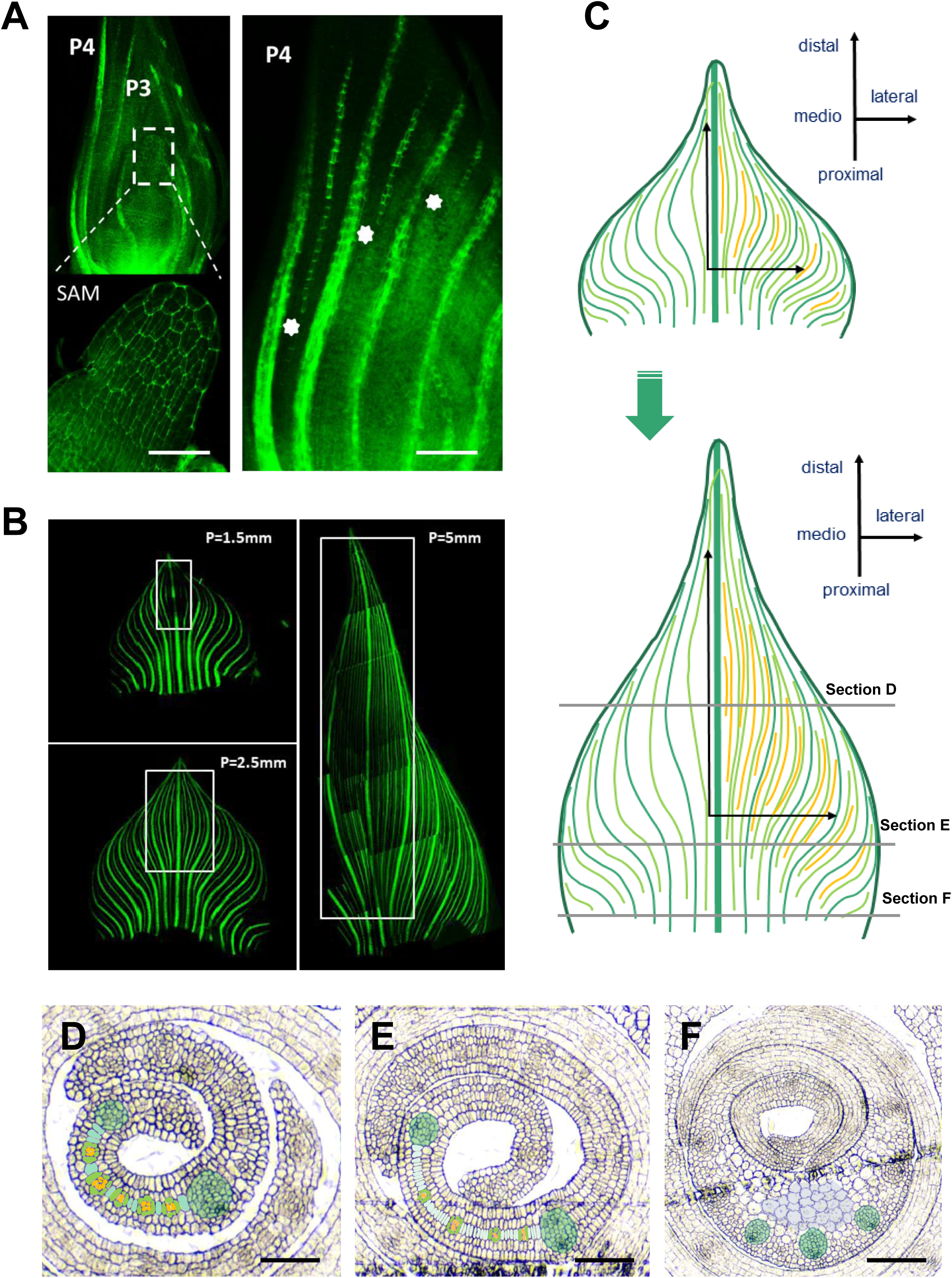
Analysis of vein development patterns in maize P4 leaf primordium with proximal–distal (PD), medial–lateral (ML), and cross-section dimensions. **A.** Confocal image showing p*Zm*PIN1a::*Zm*PIN1a:YFP fluorescence marked developing veins in maize leaf primodium. Left: SAM plus four most recently initiated leaf primordium (P1-P4). Right: greater magnification of P4, asterisks indicate the leading ends of elongating veins. P, plastochron; SAM, shoot apical meristem. Scale bar: 50 μm. **B.** The primodia of 1.5mm, 2.5mm, and 5mm in size are detached, unrolled, and flattented with the adaxial side facing up (P=1.5mm means the length of primordium is 1.5 mm). Developing veins and procambial strands marked by PIN1a-YFP fluorescence are shown to be most proliferative in the lance-shaped regions (indicated by white rectangles) closer to the mid-rib, along with PD elongation and ML expansion. **C.** Schematic depiction (according to unrolled and flattened samples in B) of vein formation and organization along with PD elongation and ML expansion of maize primordia, highlighting the recently and densely proliferated intermediate veins (marked in orange colour, and enriched in lance-shaped regions closer to mid-rib and middle-upper part of the primordium). Mid-rib and lateral veins are coloured dark green; older intermediate veins are coloured light green. **D-F.** Transverse sections of the upper middle (D), lower middle (E), and base (F) of P4 leaf primordia highlighting part of the middle layer of ground meristem cells, which give rise to the procambial strands and intermediate veins (orange) and eventually lead to the differentiation of Kranz-type bundle sheath (light green) and mesophyll cells (water green). The Kranz anatomy becomes evident in (D) with more developed bundle sheath and reduced number of mesophyll cells between veins, while in (E) multiple middle ground meristem cells or mesophyll precursors (water white) exist between veins, representing an actively developing stage of pre-Kranz anatomy. At the base section, parenchymal cells (light purple) next to the mid-rib (dark green) are indicated. Scale bar: 60 μm.

Having established the time window for intermediate vein initiation, we then serially sectioned P3, P4, and P5 leaf primordia from 2-week-old maize seedlings. Transverse sections showed different distributions of smaller intermediate veins between lateral veins across upper middle, lower middle, and the base of P4 primordia **(Figures 1D-1F)**. There was a similar but less obvious gradient in P5 primordia, with more vein extension visible. In P4 and P5 primordia, leaf sheaths are often not yet differentiated, while in the newly differentiated sheath of P6, only the midrib and lateral veins are visible, with no intermediate veins between them, similar to the base section of the blade. As a comparison, serial sections of rice primordia were also examined. Notably, the distribution of small intermediate relative to large lateral veins was similar between upper middle and lower middle sections, unlike in maize primordia of a similar size where a clear gradient was found **(Supplemental Figure 2)**. The vein pattern thus appears to establish more rapidly in the rice primordium albeit with fewer intermediate veins forming in between lateral veins.

Statistical analysis of vein numbers from tip to base confirmed a larger number of lateral and intermediate veins in the middle region of maize leaf primordia **(Figure 2A)**. By counting in both primordia and expanded blades of the same leaves, the number of veins across the broadest part was found to increase rapidly at the early primordium stage (P3-P5), and then only slowly increase until maturity **(Figure 2A)**. In order to screen for regulators of vein formation in C_4_ leaves, we conducted a transcriptomic analysis of different leaf primordium regions. We first micro-dissected and divided 5 mm long P4 leaf primordia into upper (M3tip), middle (M3middle) and lower (M3base) segments. Notably, the middle part of the spindle-rich region was extending both longitudinally and transversely with more veins initiating actively. We also dissected 3 mm long P4 leaf primordia in which the tip was not significantly elongated and the spindle-shaped area was concentrated in the middle and upper part of the primordium. These primordia were divided into two segments: M2top and M2base. Transcriptome sequencing was performed for each of the above 5 segments plus early differentiated sheath from P6 leaf primordia (Msheath) **(Figure 2B)**. At the same time, in order to exclude common regulators of vein formation in grass leaf primordia, we obtained the upper (Rtip), middle (Rmiddle) and lower (Rbase) segments of 5 mm rice leaf primordia, as well as the 2 mm primary leaf sheath (Rsheath) for comparison.

**Figure 2.**
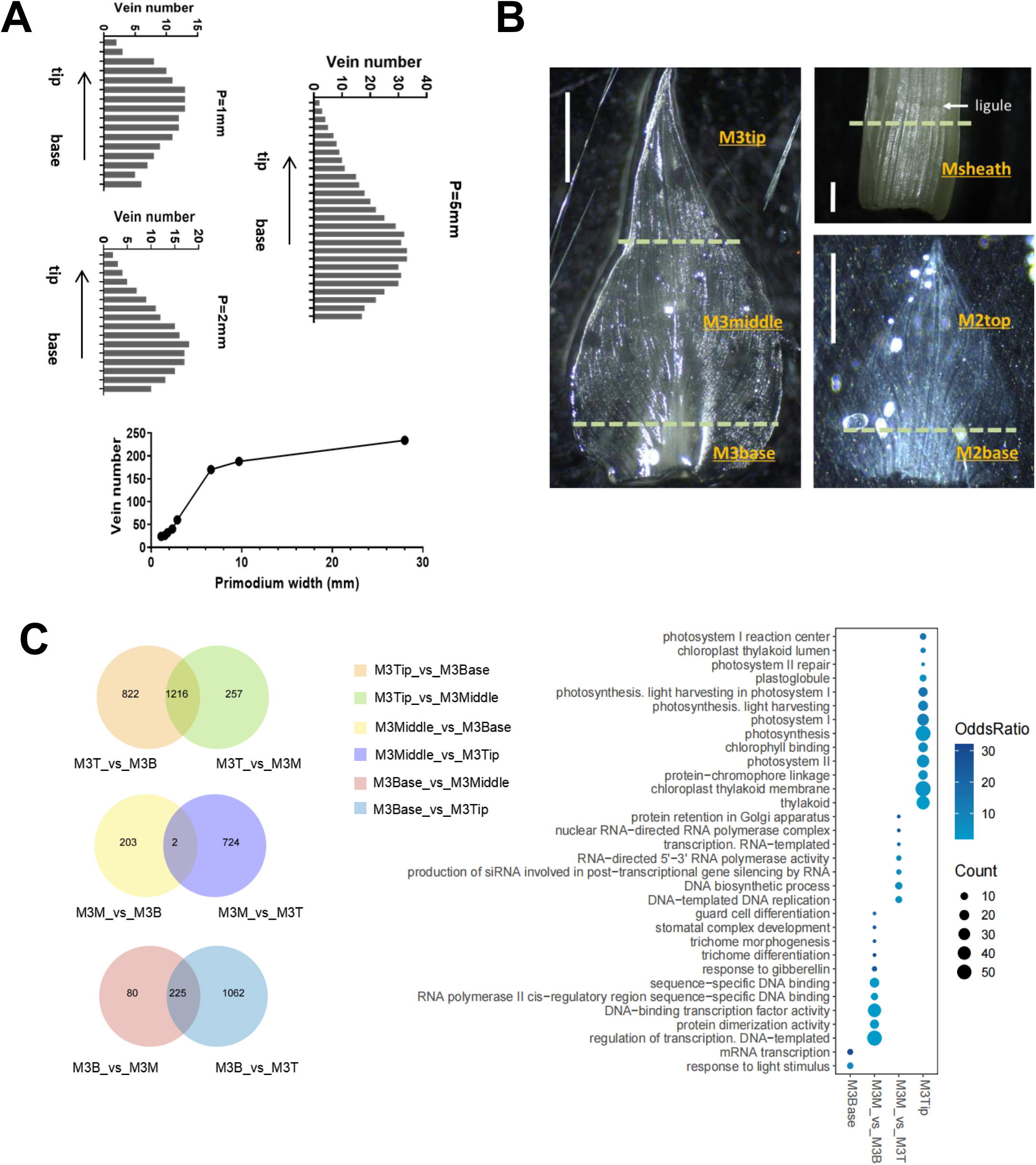
Bulk RNA-seq of leaf primordia according to proximal–distal developmental gradients. **A.** Vein number distribution in maize leaf primordia from tip to base. Veins were counted from serial paraffin sections and half of the number from each section were presented in the column charts. The longitudinal length of leaf primordia were 1 mm, 2 mm, and 5mm, respectively. In addition, a vein number growing curve from primordia or leaves of different developmental stages is presented in the line chart. Vein numbers across the maximal primordium or leaf width were counted, from samples ranging from early primordia to fully expanded leaves. **B.** Images indicating the 6 types of maize tissues sampled for bulk RNA sequencing. Left: the P4 leaf primordium of about 5 mm were partitioned under dissection microscope into three parts: M3tip≈1.5 mm, M3middle≈2.5 mm, and M3base≈1 mm. Bottom right: the P4 leaf primordium of about 3 mm were partitioned into two parts: M2top≈2 mm, and M2base≈1 mm. Top right: Msheath represents the leaf sheath of 1-2 mm, cut below the dashed line from P6. Scale bar: 1000 μm. Note: Dashed lines indicate the position where it was partitioned; Sections D and E from Figure 1 represent the Kranz developmental gradients within M3middle or M2top. **C.** Left: Venn diagram showing overlap of DEGs grouped on the basis of M3tip, M3middle, and M3base respectively. Right: Graph summarizing representative enriched GO terms describing the significantly upregulated genes in each type of tissue (from the overlap of M3tip group, the overlap of M3base group, and all of the M3middle group). Point size represents the gene counts.

### Comparative transcriptomic analysis of segmented leaf primordium from maize and rice to screen genes regulating Kranz-type vein development

To determine the extent to which M3middle and M2top were different from other samples in terms of gene expression, the numbers of differentially expressed genes (DEGs) were obtained by pairwise comparisons. As expected, M3middle exhibited large differences from M3tip (2,199 DEGs), Msheath (2,965 DEGs), and M2base (2,878 DEGs), but smaller differences from M3base (510 DEGs) and M2top (537 DEGs). The low numbers of DEGs for M3middle_vs_M2top, M3middle_vs_M3base, and M2top_vs_M3base compared to other groups **(Supplemental Figure 3A)** indicate the similarities not only between M3middle and M2top, but also the similarities of both to M3base. This is supported to certain extent by the correlation matrix shown in **Supplemental Figure 3B**. Overlapping and GO enrichment analysis were performed on 4 of the DEG lists: M3base, M3middle_vs_M3base, M3middle_vs_M3tip, and M3tip. The biological functions enriched by M3base mainly include transcriptional regulation. The biological functions enriched in M3middle were more diverse with sequence-specific DNA binding, transcription regulation, DNA biosynthesis and replication, protein interaction, and other developmental or regulatory processes. M3tip enriched biological functions were mostly related to photosynthetic processes **(Figure 2C)**.

The gene enrichment results supported an active state of transcriptional regulation and cell division in M3middle, although it is not clear how and to what extent cell division occurs for vein formation and/or for mesophyll cell development in this region. To explore the regulators of C_4_ vein initiation and development, screening of differentially expressed genes was carried out following the filtration steps and principles shown in **Figure 3A**. After considering the enrichment of active vein formation elements and excluding those shared between maize and rice, we finally obtained 224 genes with high expression in M3middle, and 142 genes with high expression in both M3middle and M2top **(Supplemental Figure 4)**. Among the 224 genes, there were 24 transcription factors, 7 transcription regulation related genes, 14 kinases, 3 auxin related genes, and 5 cell wall related genes. The functions and characteristics of the remaining genes were mostly unknown.

**Figure 3.**
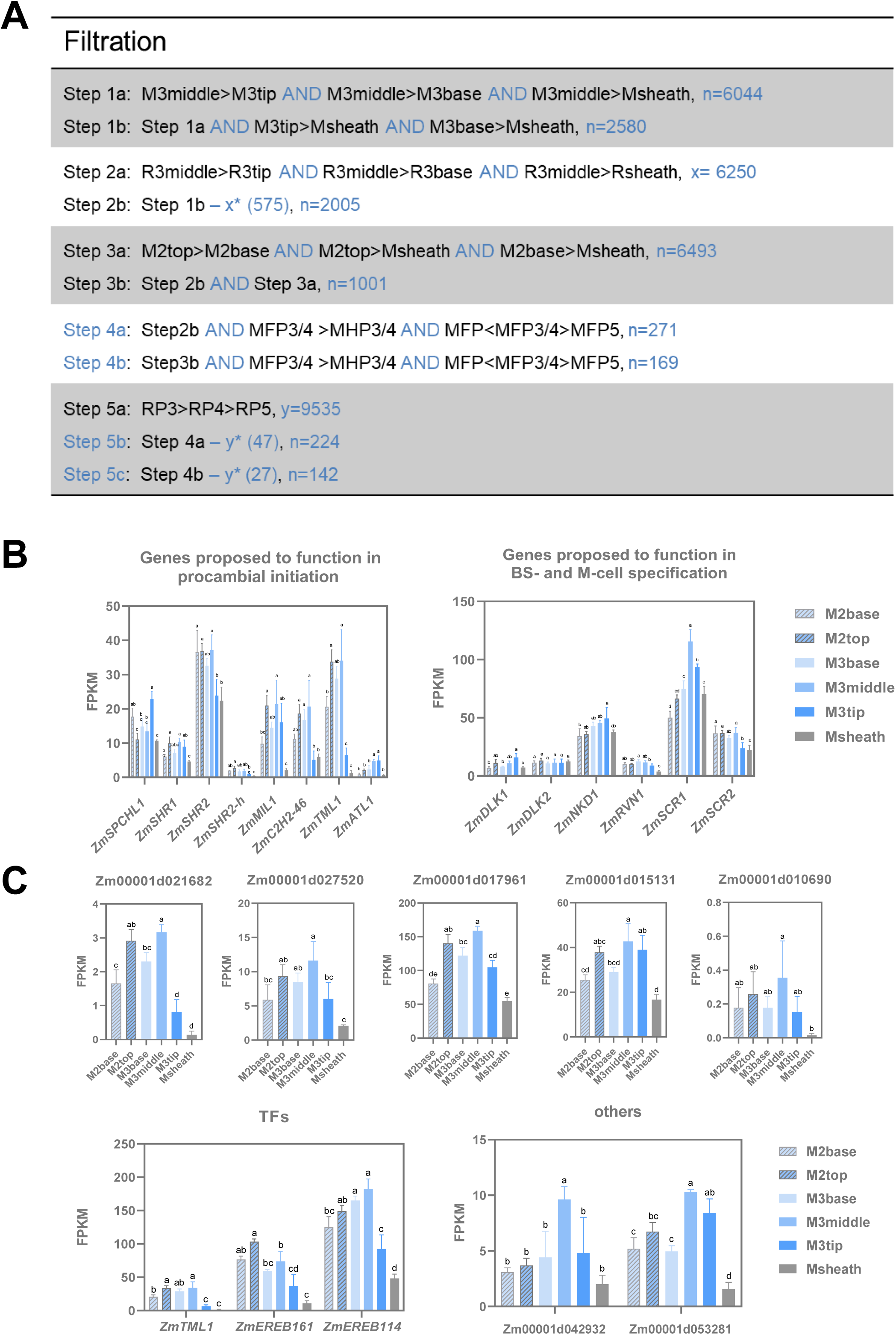
Candidate regulators of Kranz anatomy development. **A.** Filtration steps involving the maize and rice samples to identify putative regulators of Kranz anatomy. M: maize; R: rice; FP: foliar primordium; HP: husk primordium; RP: rice primordium. The transcriptome data from Wang et al., 2013 (SRS394616–SRS394626) and van Campen et al., 2016 (Supplemental Data S1) were used for comparison. The filtration was based on the differential transcript abundance of the given gene among the samples for comparison. Step 1: By comparing FPKM values, 2580 genes were obtained with higher expression in 5 mm maize leaf primordia than in leaf sheath and highest expression in M3middle. Step 2: In the 5 mm rice leaf primordium, 6250 genes were obtained with the highest expression in R3middle; By comparing and analysing the genes highly expressed in the middle of leaf primordia of maize and rice, and excluding the background of developmental gradient of grass leaves, 2005 genes specifically and highly expressed in M3middle were obtained. Step 3: 6493 genes were obtained in 3 mm maize leaf primordium, with the highest expression in M2top; Considering the similarity of Kranz developmental status in M3middle and M2top, 1001 genes highly expressed in both samples after removing the grass leaf background were obtained. Step 4: Previous work obtained 2935 genes highly expressed in P3/4 leaf primordia by comparing with P4, P5 leaf primordia, and husk primordia (Wang et al., 2013); After comparing with 2005 genes with high expression in M3middle (from step 2), 271 common highly expressed genes were obtained; Compared with 1001 genes highly expressed in both M3middle and M2top (from step 3), 169 common highly expressed genes were obtained. Step 5: Another previous work reported 9535 genes with higher expression in P3 than in P4 and P5 of rice leaf primordia (van Campen et al., 2016); Using the above information to further exclude developmental background of grass leaves, we finally obtained 224 genes with high expression in M3middle, and 142 genes with high expression in both M3middle and M2top. Steps 2b, 5b, and 5c were performed to exclude the common background of maize and rice leaf development. In step 2b, x* (575) is the number of maize homolog genes available out of the 6250 rice genes from step 2a, that were also found in the 2580 genes from step 1b. Similarly, y* (47) is the number of maize homolog genes available out of the 9535 rice genes from step 5a, that were also found in the 271 genes from step 4a. *y (27) is the number of maize homolog genes available out of the 9535 rice genes from step 5a, that were found in the 169 genes from step 4b. **B.** Bar graph illustrating the expression patterns of previously proposed genes (Wang et al., 2013; Fouracre et al., 2014) across 6 maize tissue types. **C.** Bar graph illustrating the expression profiles of putative regulators of Kranz anatomy identified in this study. The gene list of B and C is given in Supplemental dataset 7. Error bars represent mean±SD (n=3). Statistical analysis in B and C was performed using one-way ANOVA with Tukey’s HSD test; *P* < 0.05, different letters on the bar graphs indicate statistically significant difference.

In the screening process we found that some previously proposed regulators of Kranz development (Wang et al., 2013; Fouracre et al., 2014) were highly expressed in M3middle and M2top, including *ZmSHR1*, *ZmSHR2*, *ZmSHR2-h*, *ZmSCR1*, *ZmSCR2*, *ZmMIL1*, *ZmC2H2-46*, and *ZmTML1* (previously named Zm*DOT1*) **(Figure 3B)**. Some of these genes also had similar expression patterns in rice leaf primordia, including *ZmSHR1*, *ZmSCR1*, and *ZmMIL1* **(Supplemental Figure 5A)**, suggesting these genes may have relatively conserved functions in grass leaf development. To identify potential C_4_ specific regulators we therefore further analysed the 224 genes that were expressed highly in M3middle samples. Presented in **Figure 3C** were ten genes highly expressed in M3middle and M2top samples (passed significance test for all the pairwise comparisons, *P*<0.05) whereas their homologous genes were not obviously expressed at higher levels in Rmiddle samples of rice **(Supplemental Figure 5B)**. Three of the 10 candidate genes encoded transcription factors, including *EREB161* (Zm00001d048004) and *EREB114* (Zm00001d018731) of the AP2 family, which were speculated to influence auxin signalling during vein development (Kitomi et al., 2011; Liu et al., 2022), and *ZmTML1* (Zm00001d020037) of the C2H2 family, which was recently shown to specify vein rank (Vlad et al., 2024).

### Construction of a single-nucleus transcriptome atlas of maize early leaf primordia

To further explore the early development of maize leaves at a single-cell level, we carried out single-nucleus RNA sequencing (snRNA-seq) using 3-4 mm leaf primordia of P4 in which veins were actively developing. We obtained 7,473 effective cells with an average of 2,293 genes expressed in each nucleus. In total, we identified 25,009 genes, which represented 78.2% of the 32,000 predicted protein coding genes in maize. Dimensionality reduction and cluster analysis (t-SNE and UMAP) were performed on the snRNA-seq data, and the 7,473 high quality nuclei were divided into 14 different clusters **(Figure 4A)**. By analyzing DEGs between the clusters, we identified a series of cluster-enriched or specific genes. Dot plots of the top 10 marker genes from each of the different cell clusters were generated **(Supplemental Figure 6)** and a subset is shown in **Figure 4B**. The cellular anatomy of leaf primordia sampled for single cell analysis is such that five cell layers are identifiable across the adaxial-abaxial leaf axis: the upper and lower epidermis and three internal cell layers. The veins, bundle sheath cells and mesophyll cells between veins all differentiate from the middle layer ground meristem (mGM) **(Figures 1D-1F)**. Using marker genes reported in existing references, we were able to assign 14 clusters to the different cell-types observed.

**Figure 4.**
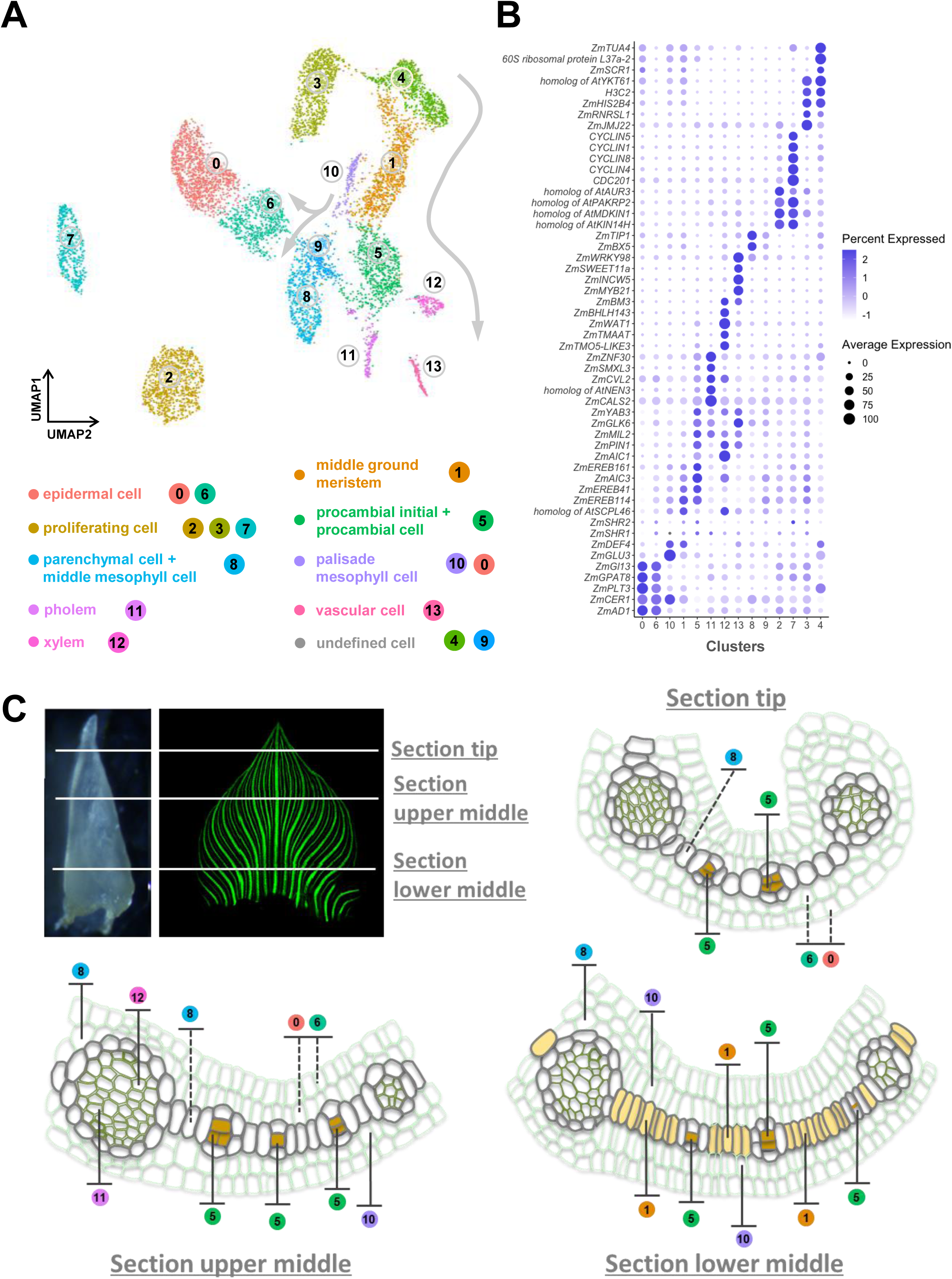
Cell heterogeneity in the maize P4 leaf primordium. **A.** Visualization of 14 cell clusters using UMAP. Dots, individual cells; n = 7473 cells; cell clusters are coloured differently and labelled with circled numbers. The grey line and arrow points to the putative developmental direction from cluster 1 toward clusters 5, 11, 12 and 13, as well as from cluster 10 toward clusters 6 and 9. **B.** Expression patterns of representative cluster-specific marker genes. Dot colour, proportion of cluster cells expressing a given gene; Dot size, the average expression level. **C.** Schematics of the anatomy and cell types representing the tip, upper middle, and lower middle sections of P4 leaf primordium. Bundle sheath cells and the middle layer mesophyll cells were profiled in dark grey colour. In the bottom right schematic, the middle layer of ground meristem cells were filled with orange colour, to highlight their pre-differentiated status. Cell clusters identified in (A) were projected onto the related cell types with coloured cycles and numbers. Dotted lines indicate annotations cross-referenced from Liu et al. (2022).

First of all, *ZmPIN1* as a marker gene of procambium and vascular tissue, was found in the cells of cluster 5 and cluster 12 in the UMAP **(Figure 5A)**. Cross-referencing to our candidate genes from comparative transcriptomic analysis of segmented leaf primordia (**Figure 3C**, TFs) and to results from Vlad et al. (2024), we presented here that *ZmTML1* was clearly located in cluster 5, while *ZmEREB161* and *ZmEREB114* were enriched in both clusters 1 and 5, with the former more distributed in cluster 5 and latter more distributed in cluster 1 **(Figures 4B, 5A and 5B)**. *In situ* hybridization assays confirmed that *ZmEREB114*, as well as *ZmEREB41* (both are co-orthologs of arabidopsis *ANT1*), were expressed in both the undifferentiated mGM and the early procambial cells **(Figures 5B, 5C and 6)**, whereas a homolog of *AtSCPL46* which is evident in cluster 1, was shown to be mainly restricted in undifferentiated cells of the mGM **(Figure 5D)**. *ZmEREB161* was expressed in both the initiating procambial cells and the differentiated procambial cells, while the expression of *SHR1* was detected predominantly in the developed procambium although in lower level **(Figure 7; Supplemental Figure 7A)**. Futher, *ZmYAB3* and *ZmGLK6* transcripts were detected by *in situ* hybridization in developing procambial cells in a way similar to that of *ZmEREB161*, with their distribution more concentrated in cluster 5 in the UMAP **(Supplemental Figure 7C and 7D)**. Considering the continuity of developmental status, we thus annotated cluster 1 as “middle ground meristem” and cluster 5 as “procambial initial + procambial cell”.

**Figure 5.**
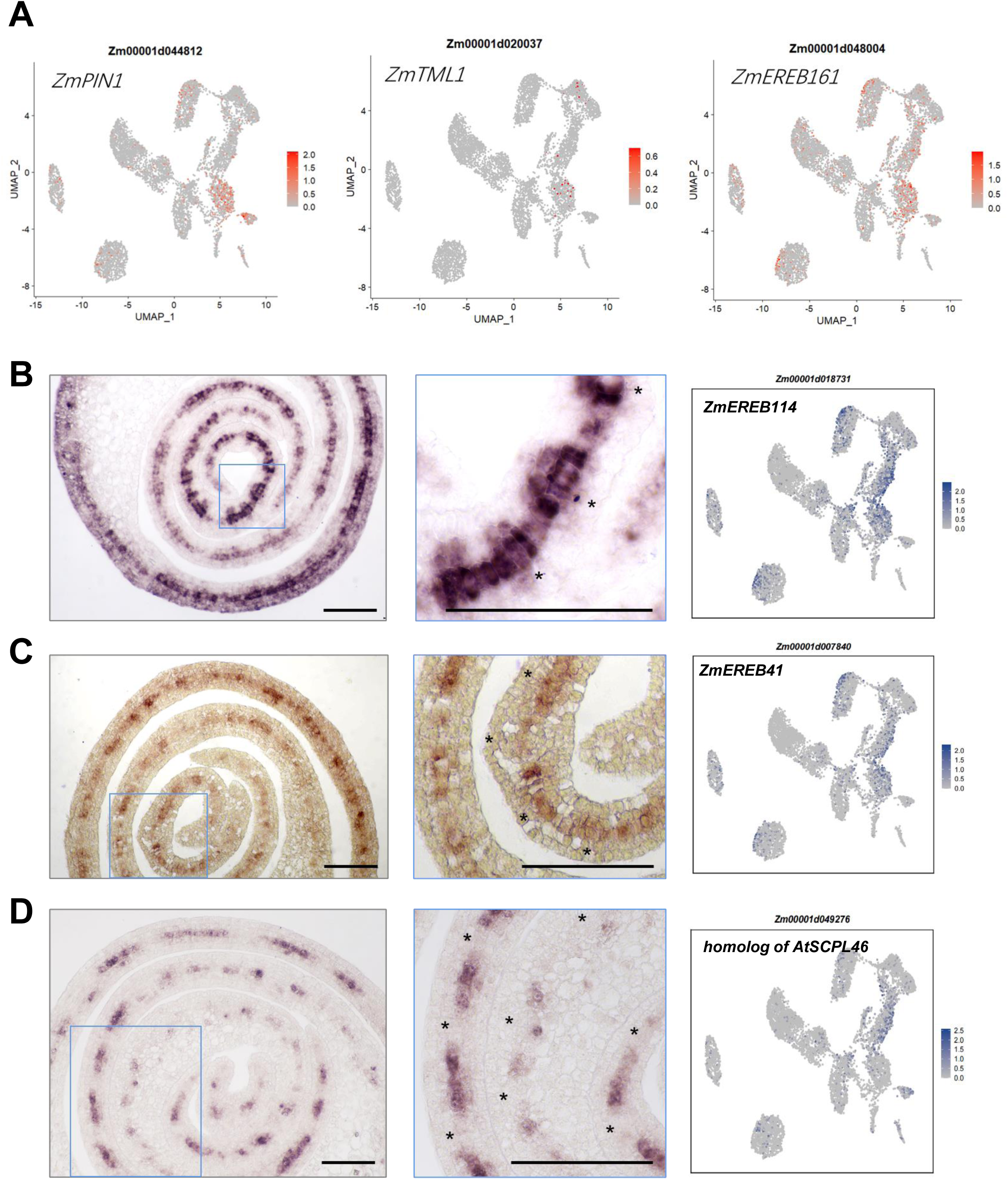
*In situ* expression profiles of representative marker genes from maize middle layer ground meristem cells. **A.** Expression pattern of *ZmPIN1*, *ZmTML1* and *ZmEREB161* by UMAP plots. **B-D.** Left column, *in situ* hybridization for the transcript localization of cluster 1 and 5-enriched genes on transverse sections of maize leaf primordia. Middle column, close-up images of the blue square framed regions from left column. Right column, expression pattern of selected genes by UMAP plots. Scale bar: 100 μm. Asterisks indicate procambia.

**Figure 6.**
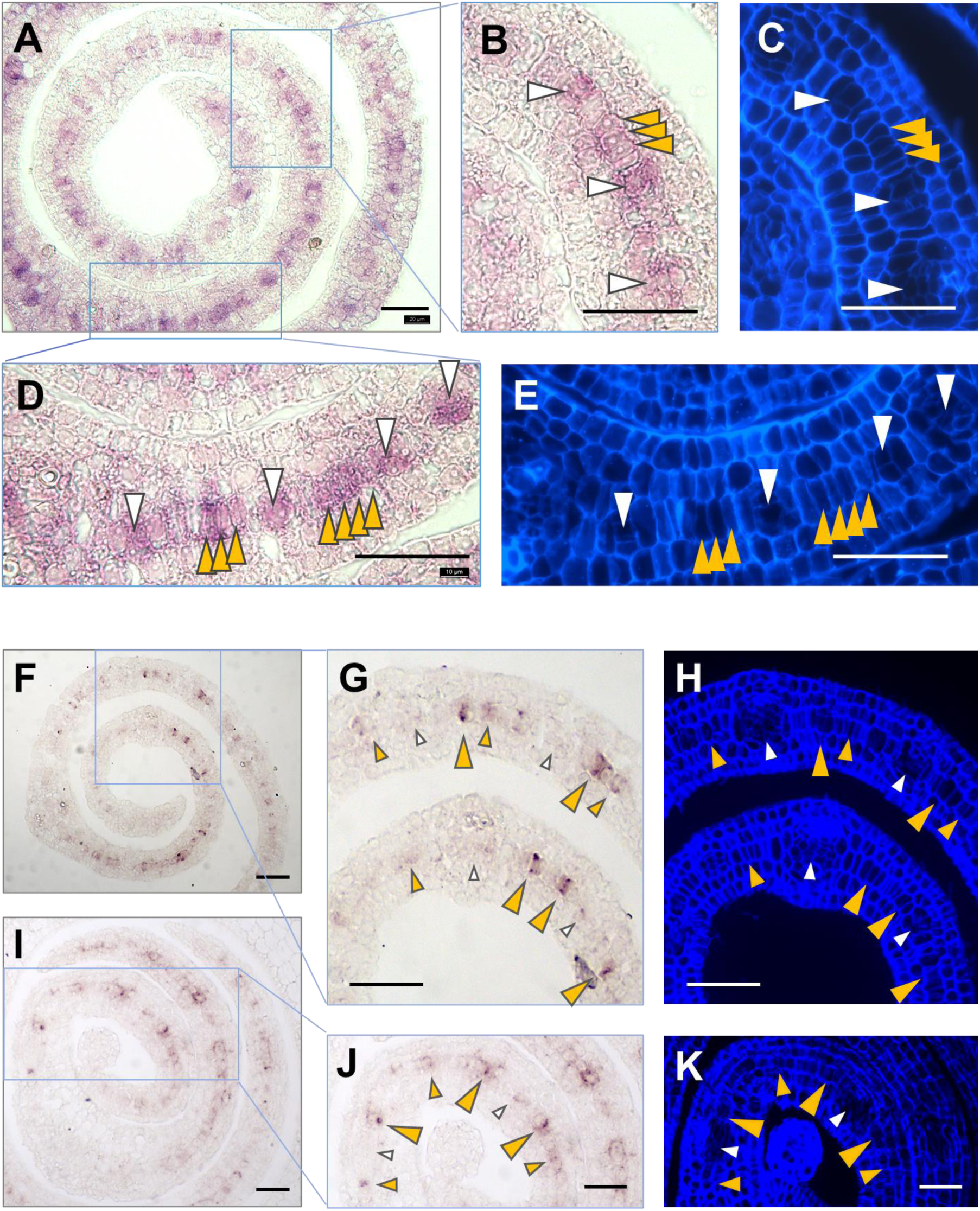
Localization of *ZmEREB114* and *ZmEREB41* transcripts in the middle cell layer of maize leaf primordium. **A.** *In situ* hybridization showing the distribution of *ZmEREB114* transcript in the middle section of P4 leaf primordia. Scale bar: 40 μm. **B and D.** Magnified images of the blue square framed regions from (A), showing *ZmEREB114* signals at different cell types of early Kranz anatomy; **C and E.** UV activated fluorescent images of the corresponding regions from (B) and (D). Scale bar: 40 μm. **F and I.** *In situ* hybridization showing the distribution of *ZmEREB41* transcript in the middle section of P4 leaf primordia. Scale bar: 40 μm. **G and J.** Magnified images of the blue square framed regions from (F) and (I), showing *ZmEREB41* signals at different cell types of early Kranz anatomy; **H and K.** UV activated fluorescent images of the corresponding regions from (G) and (J). Scale bar: 40 μm. White arrows indicate the differentiating or differentiated procambium (including early differentiating ones with the primary adaxial and abaxial precursors of BS cells, generated by the 1^st^ and 2^nd^ periclinal divisions of a single procambial initial cell); orange arrows indicate the potential procambial initial cells (indicated by *in situ* hybridization signals) prior to periclinal division. Smaller sized arrows indicate less strong *in situ* hybridization signals.

**Figure 7.**
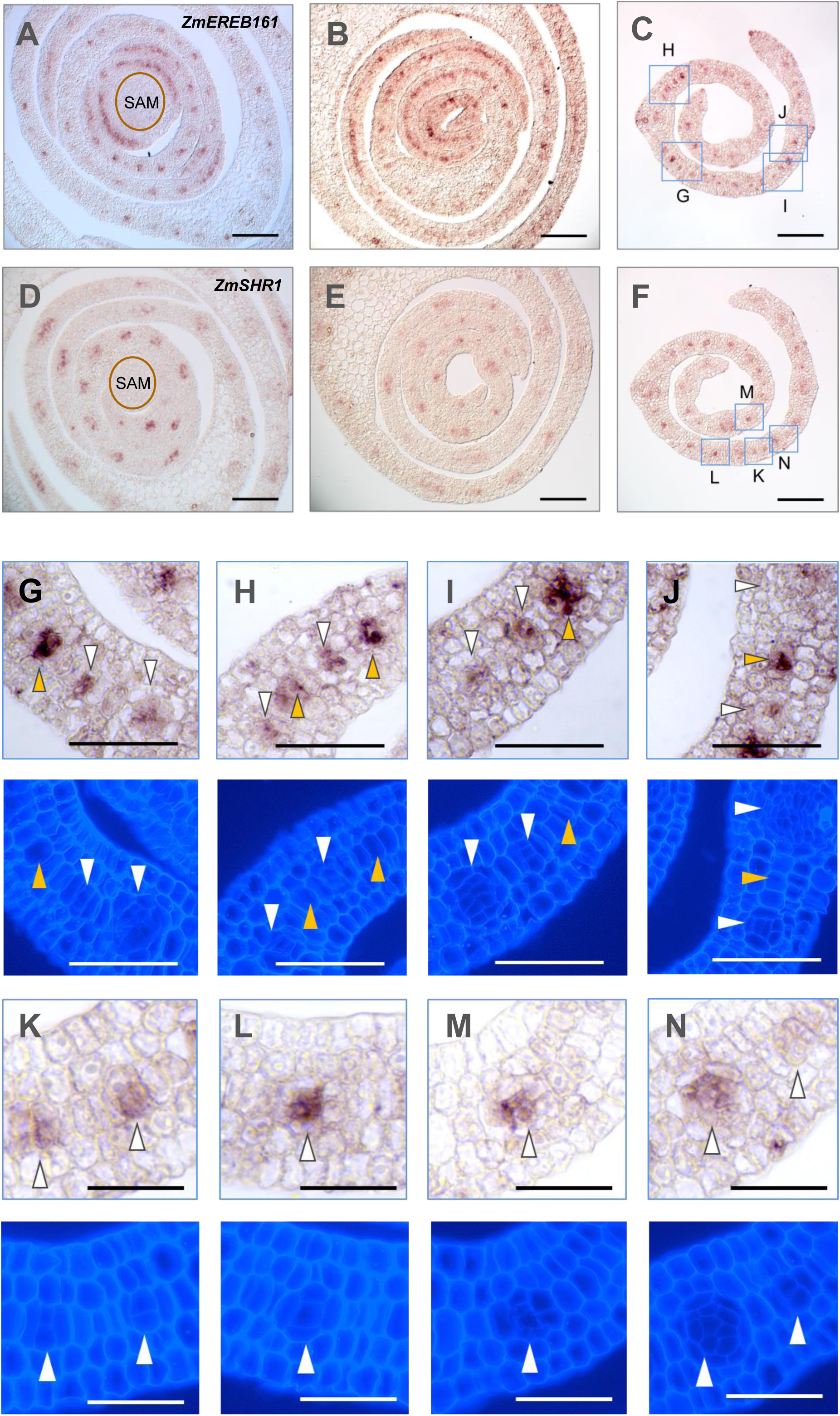
Localization of *ZmEREB161* and *ZmSHR1* transcripts in the middle cell layer of maize leaf primordium. **A-C.** *In situ* hybridization showing *ZmEREB161* transcript densely distributed in younger leaf primordia (A), toward the margin of larger leaf primordia sectioned in the base (B), and in the middle section of P4 leaf primordia (C). Scale bar: 120 μm. **D-F.** *ZmSHR1* transcript distribution in the vascular bundles and larger procambium strands, found from similar tissue types of primordia as in (A), (B), and (C). Scale bar: 120 μm. **G-J.** Upper panel: Magnified images of the blue square framed regions from (C), showing *ZmEREB161* signals at different developmental stages of early Kranz anatomy; lower panel: UV activated fluorescent images of the corresponding regions from upper panel. Scale bar: 50 μm. **K-N.** Upper panel: Magnified images of the blue square framed regions from (F), showing *ZmSHR1* signals at different developmental stages of early Kranz anatomy; lower panel: UV activated fluorescent images of the corresponding regions from upper panel. Scale bar: 50 μm. White arrows indicate the differentiating procambium (including those with the primary adaxial and abaxial precursors of BS cells, generated by the 1^st^ and 2^nd^ periclinal divisions of a single procambial initial cell); orange arrows indicate the potential single procambial initial cell (indicated by *in situ* hybridization signals) prior to periclinal division.

Clusters 0 and 6 were identified to be related with epidermal cells, cluster 11 as primary phloem, cluster 12 as primary xylem, and cluster 13 as undefined vascular cells (maybe related with phloem, xylem, and other vascular cells). Cluster 10 was identified as palisade mesophyll cells, and cluster 8 was identified to be related with middle layer mesophyll and/or parenchyma cells. These annotations were verified by selecting the enriched marker genes for *in situ* hybridization **(Figure 4; Supplemental Figure 8)**. *GLU3* which is enriched in cluster 10 and encodes a β-glucosidase that participates in the metabolism of cellulose and modifies cell wall polysaccharides during the elongation growth of cells (Bosch et al., 2011; Nazipova et al., 2022), was specifically expressed in mesophyll cells adjacent to the abaxial epidermis, or in mesophyll cells both adaxially and abaxially close to the lateral veins **(Figure 8A)**. *ZmWIP1* (encoding a Bowman-Birk type wound-induced proteinase inhibitor) (Rohrmeier and Lehle, 1993) which is enriched in cluster 8 (also present in cluster 0) was distributed in the swollen parenchyma cells (also detected in epidermal cells) close to the midrib and in the adaxial epidermis **(Figure 8B)**. *In situ* hybridization further showed that cluster 8 was composed of not only the enlarged parenchyma cells close to the midrib, but also those surrounding other larger vascular strands (including mesophyll cells in the middle layer), in which a cytochrome P450-dependent monooxygenase gene *BX5* (required for 2,4-dihydroxy-1, 4-benzoxazin-3-one biosynthesis, involved in chemical plant defense mechanism) was preferentially expressed **(Figure 8C)** (Frey et al., 1997).

**Figure 8.**
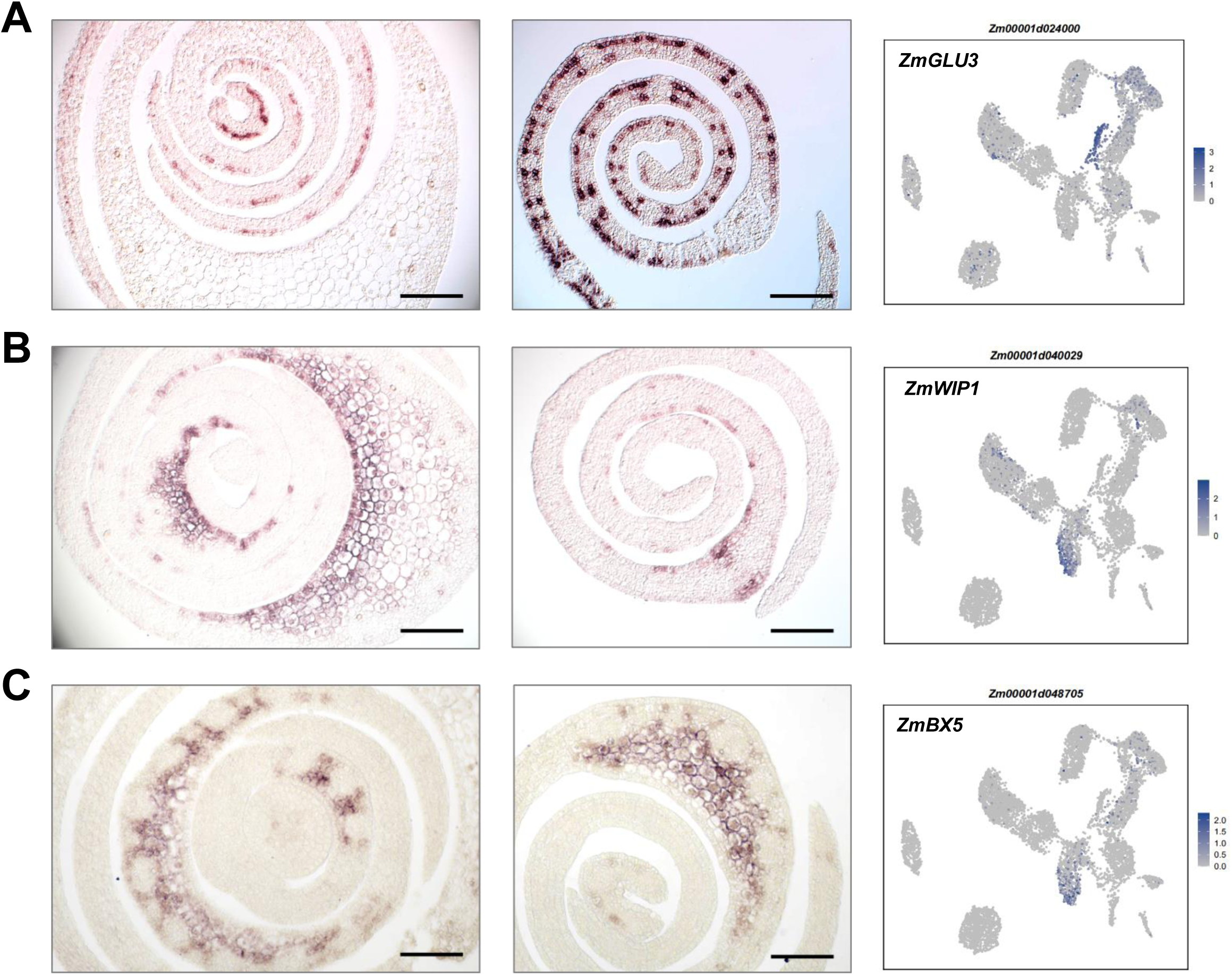
*In situ* expression patterns of representative marker genes from mesophyll and parenchymal cell related clusters. Left column, *in situ* hybridization for the expression patterns of cluster 10 (A) and 8 (B and C)-enriched genes on transverse sections of maize leaf primordia. Middle column, *in situ* hybridization with extra transverse sections toward the middle-upper position of primordia. Right column, expression pattern of selected genes by UMAP plots. Scale bar: 120 μm.

Cluster 9, which remains to be further characterized, may contain a group of cells at the developmental junction between mGM and mesophyll/parenchymal cells. Cluster 4 is also unknown, with enrichment of a large number of ribosomal protein-related genes. *In situ* hybridization of selected genes presenting in cluster 4 showed expression in veins, in cells around veins, and in the epidermis **(Supplemental Figures 9A and 9B)**. For the other remaining clusters, the presence of cell division markers suggests that cluster 3 is likely to represent proliferative cells in S phase, and clusters 2 and 7 proliferating cells in G2/M phase **(Supplemental Figures 9C and 9D)**.

**Figure 9.**
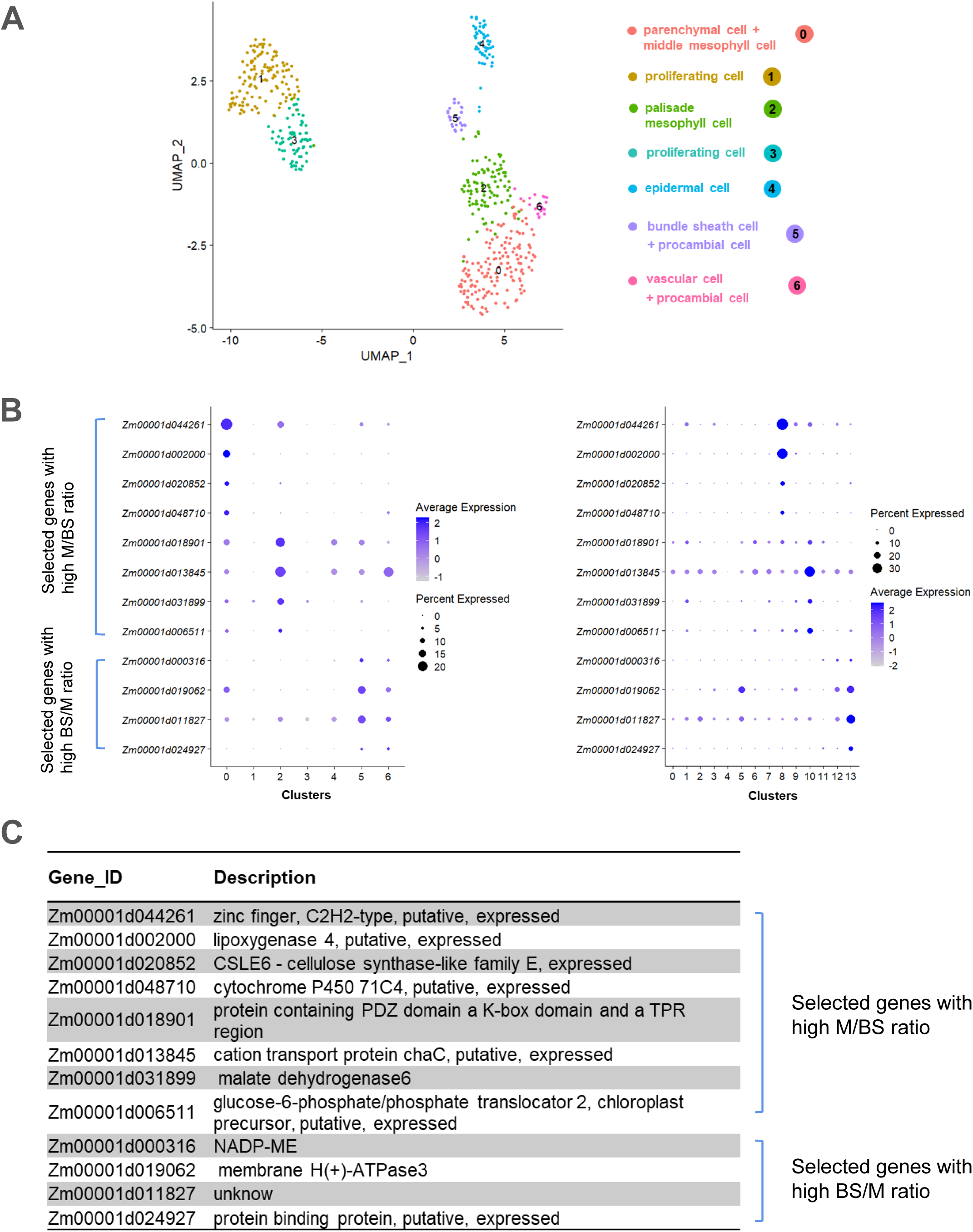
Cross-reference of photosynthetic genes between total primordium and M3tip UMAP clusters. **A.** Visualization of 7 cell clusters using UMAP. Dots, individual cells; n = 535 cells; cell clusters are coloured differently and labelled with numbers. **B.** Distribution of selected genes with high BS/M or M/BS ratio of expression in the 7 clusters from (A) (left), and in the 14 clusters from Figure 4A (right). **C.** Annotation and description of genes listed in (A).

We have identified seven major cell types: epidermal cells, mesophyll cells, parenchyma cells, proliferating cells, middle layer ground meristem, procambium, and vascular tissue cells (including xylem and phloem). Specifically, cluster 1 represents a larger group of cells containing the pre-procambial cells and other middle layer ground meristem cells, whereas cluster 5 contains cells derived from cluster 1 that are on a developmental trajectory towards procambium and vein formation. Importantly, because the 3-4 mm leaf primordia showed strong cellular heterogeneity, new marker genes for each cell type were suggested **(Supplemental Figure 6)** and early stages of cell differentiation in the maize leaf have been revealed.

### Comparing the cell-specific expression of *ZmEREBs* and *ZmSHR1* in the middle cell layer of leaf primordium

To further investigate the early transition of mGM cells toward Kranz-type leaf anatomy, we compared the expression patterns of *ZmEREB114*, *ZmEREB41*, and *ZmEREB161* (they are co-orthologs of arabidopsis *ANT1*) at cellular resolution by combination of *in situ* hybridization and fluorescent images. *ZmEREB114* expression was not only detected in vascular procambia (white triangle indicated), but also observed in the three or four-contiguous cells (orange triangle indicated) between existing procambial centres **(Figure 6A-6E)**. For *ZmEREB41*, with less developed *in situ* hybridization signals than **Figure 5C**, its expression appeared with stronger signals in single procambial initial cells (orange triangle indicated) flanking procambial centres, while less strong signals (smaller orange triangle indicated) were found further away from the procambial centres **(Figure 6F-6K)**.

More clearly, *ZmEREB161* was strongly expressed in the single procambial initials of intermediate veins, in addition to the procambia of mid and lateral veins **(Figures 7A-7C)**. Taking *ZmSHR1* for comparison, we examined the expression of *ZmEREB161* in the context of cell division and arrangement patterns **(Figure 7)**. *In situ* hybridization showed that the distribution of *ZmEREB161* expression appeared not only in different developmental stages of procambium (white triangle indicated), but also evidently in mGM cells that had not yet undergone peripheral division (orange triangle indicated, may be equivalent to one of the three-contiguous cells, at 3C stage, in Liu et al., 2022) several cells away from the procambium **(Figures 7G-7J)**. The peripherally divided cell in the middle of the three adjacent mGM cells produced two cells, with one of them expressing the *ZmSHR1* gene. With the second peripheral division, *ZmSHR1* gene expression was detected in the middle of the three progeny cells. During subsequent cell division and differentiation, *ZmSHR1* was confined to procambial cells within vascular bundles **(Figures 7D-7F, 7K-7N)**. Therefore, the localization of *ZmSHR1* was preceded by that of *ZmEREB161* and *ZmEREB114* during procambium initiation and development.

### Indication of bundle sheath related cell clusters by integrating M3tip section and snRNA-seq data

Most of the genes specifically expressed in M cells and BS cells are photosynthesis related genes identified from mature leaves (Bezrutczyk et al., 2021). Due to the low differentiation degree of the two types of cells in the leaf primordium, these genes are generally not expressed. However, since M3tip section seems enriched with photosynthesis-related genes, the photosynthetic gene expressing cells could be clustered to see whether they display better distinguished BS cell types, which may also serve as a good cross-linking of the bulk and single-cell RNA-seq. As chloroplasts were not yet developed in the proplastid state in the primordia **(Supplemental Figure 10A)**, photosynthetic genes (although enriched by GO term in M3tip) were under-expressed in our snRNA-seq data, with no significant enrichment in the total cell clusters. Dot plots show the expression distribution pattern of these photosynthesis genes **(Supplemental Figures 10B and 10C)**, with relatively higher expression level and percentage of cells expressed in clusters 1, 4, 8,9,10, 13, but not 5, 11 and 12. This makes sense because cells in clusters 5, 11 and 12 are developing toward vascular cells while those in clusters 8, 9, 10 are developing toward photosynthetic cells. The interesting distribution of photosynthesis genes in clusters 4 and 13 may provide special clues for cluster annotation.

**Figure 10.**
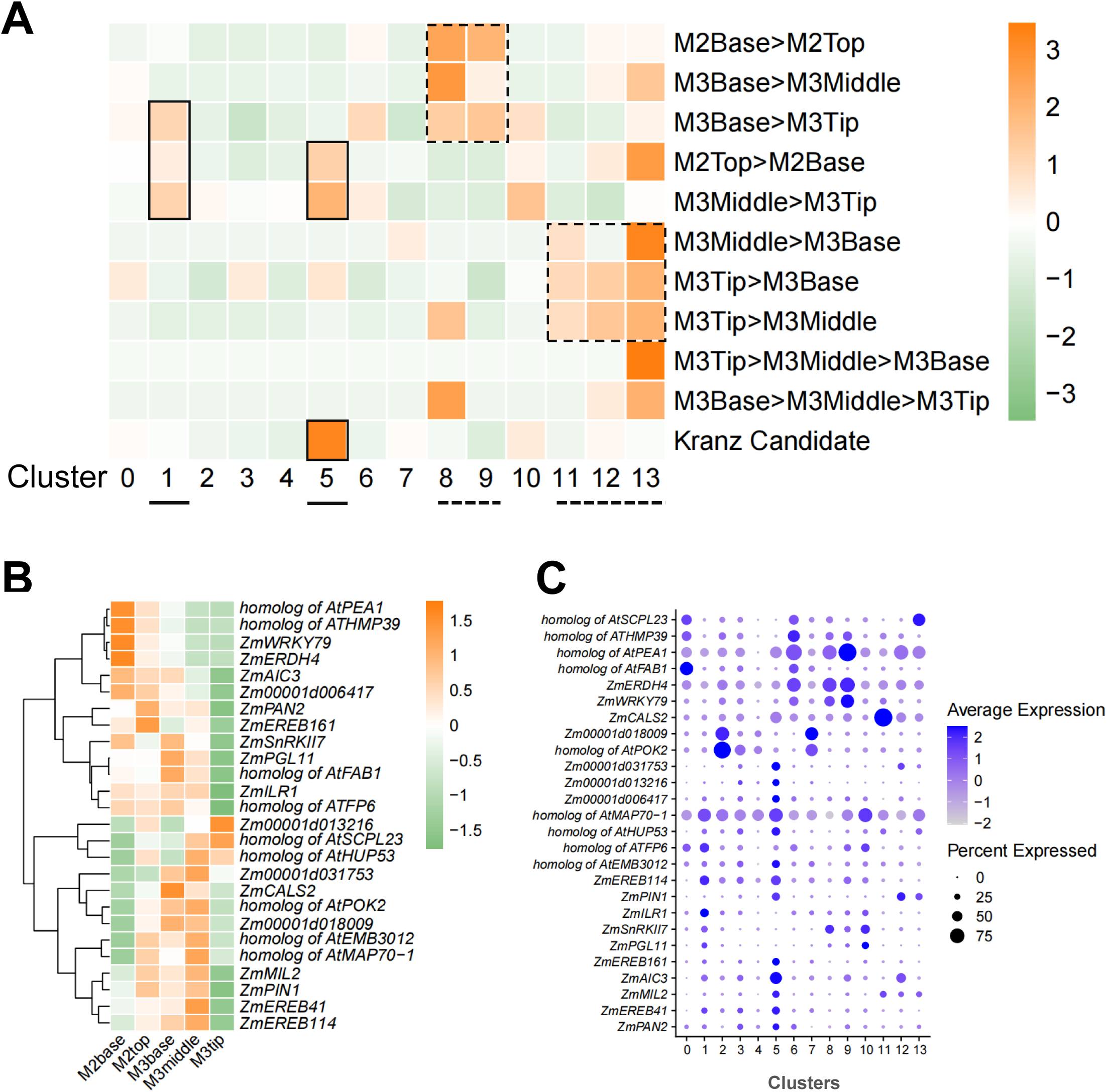
SnRNA-seq versus gradient bulk RNA-seq of leaf primordium: finding spatial transcriptional signatures. **A.** The distribution frequency of DEGs (number of overlapped divided by the number from corresponding subsection comparison on the right) between primordium subsections and each cluster of single cells. The analysis was partly drawn from Figure 3A, but instead of differential transcript abundance, here it was based on the number of differentially expressed genes. “Kranz candidate” represents the 224 genes obtained in Figure 3A. Kranz candidate genes and part of the DEGs enriched in M2top and M3middle were projected to clusters 1 and 5. The solid or dotted boxes help to indicate the different groups of projections. **B.** Heat map grouping of the expression patterns of the boxed genes among leaf subsections. **C.** Dot diagram showing the expression patterns of the boxed genes among different cell clusters.

We have extracted cell groups (3681 cells) expressing the M3tip enriched photosynthetic genes (91 genes) to perform clustering analysis. We then took the top 20% of these genes, and new UMAP clusters (based on 535 cells represented) were obtained as shown in **Figure 9A**. Importantly, we were able to infer that bundle sheath (BS) cells (or BS precursors) were potentially included in the new cluster 5, based on the enrichment of genes with high BS/M ratio of expression. In contrast, mesophyll (M) cells were identified to be included in clusters 0 and 2 of the new UMAP based on the enrichment of genes with high M/BS ratio of expression **(Figures 9B and 9C; Supplemental Figure 11**, BS/M data sourced from Li et al., 2010). Further, when looking back in the total cell clusters, cluster 13 from the original UMAP **(Figure 4A)** potentially also carried BS cell (or its precursor) identity. This was because selected new cluster 5-enriched genes (*NADP-ME* for example) could be re-plotted to the original cluster 13 **(Figures 9B and 9C)**, and some BS-enriched genes (*NDHU* for example) were also found accumulated in cluster 13 **(Supplemental Figures 10B and 10C)**. However, our *in situ* hybridizations haven’t detected distinctive BS specific expression of genes in the primordia, although the BS-enrichment of transcripts was detected in expended leaves **(Supplemental Figure 11E-11H)**. It could be attributed to low transcript levels, insufficient cell differentiation, or the lack of light-induction for BS/M specific distribution of C_4_ enzymes (Langdale et al., 1988), the regulatory factors of which could be further investigated on the basis of our cell clustering analysis described above.

**Figure 11.**
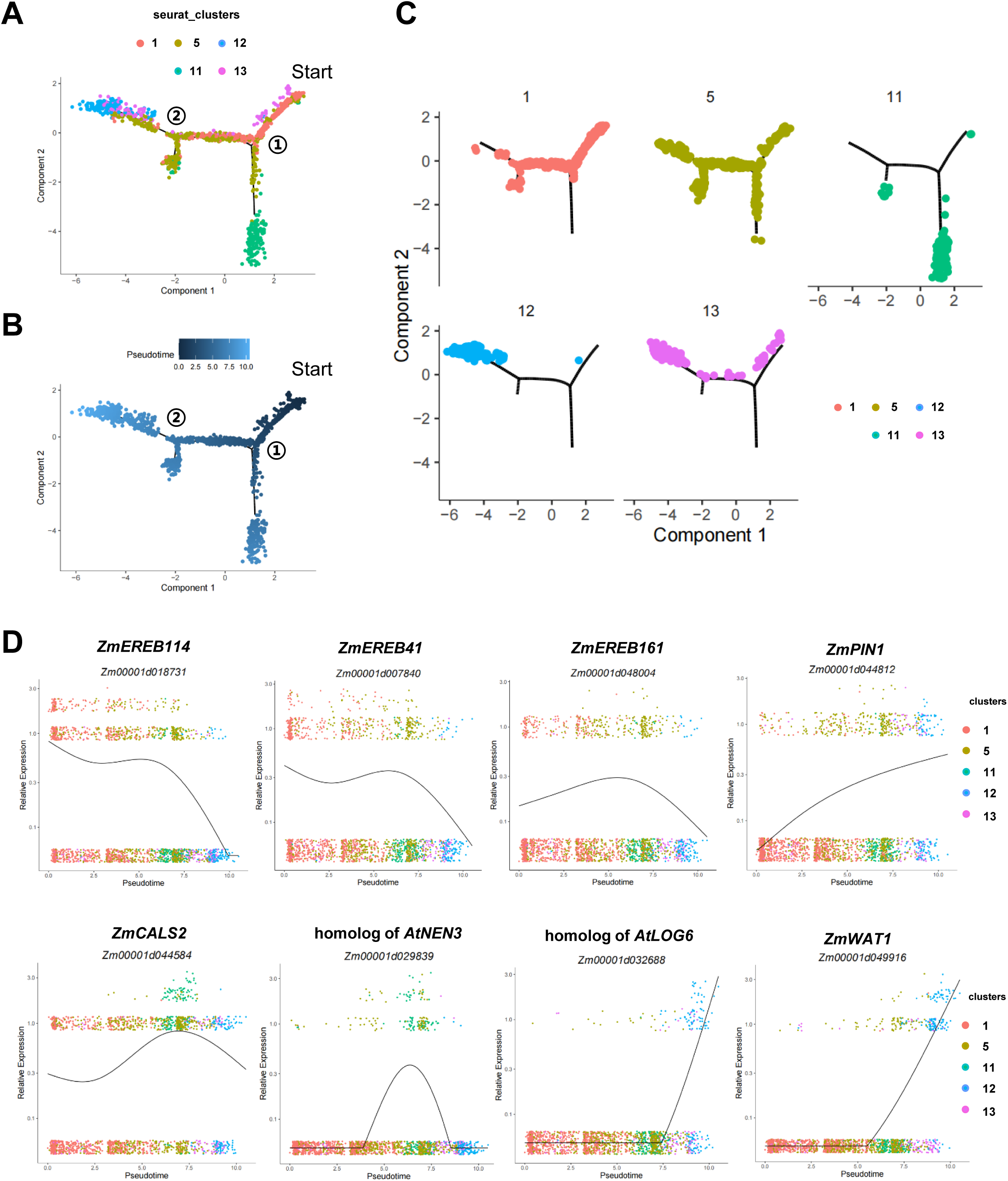
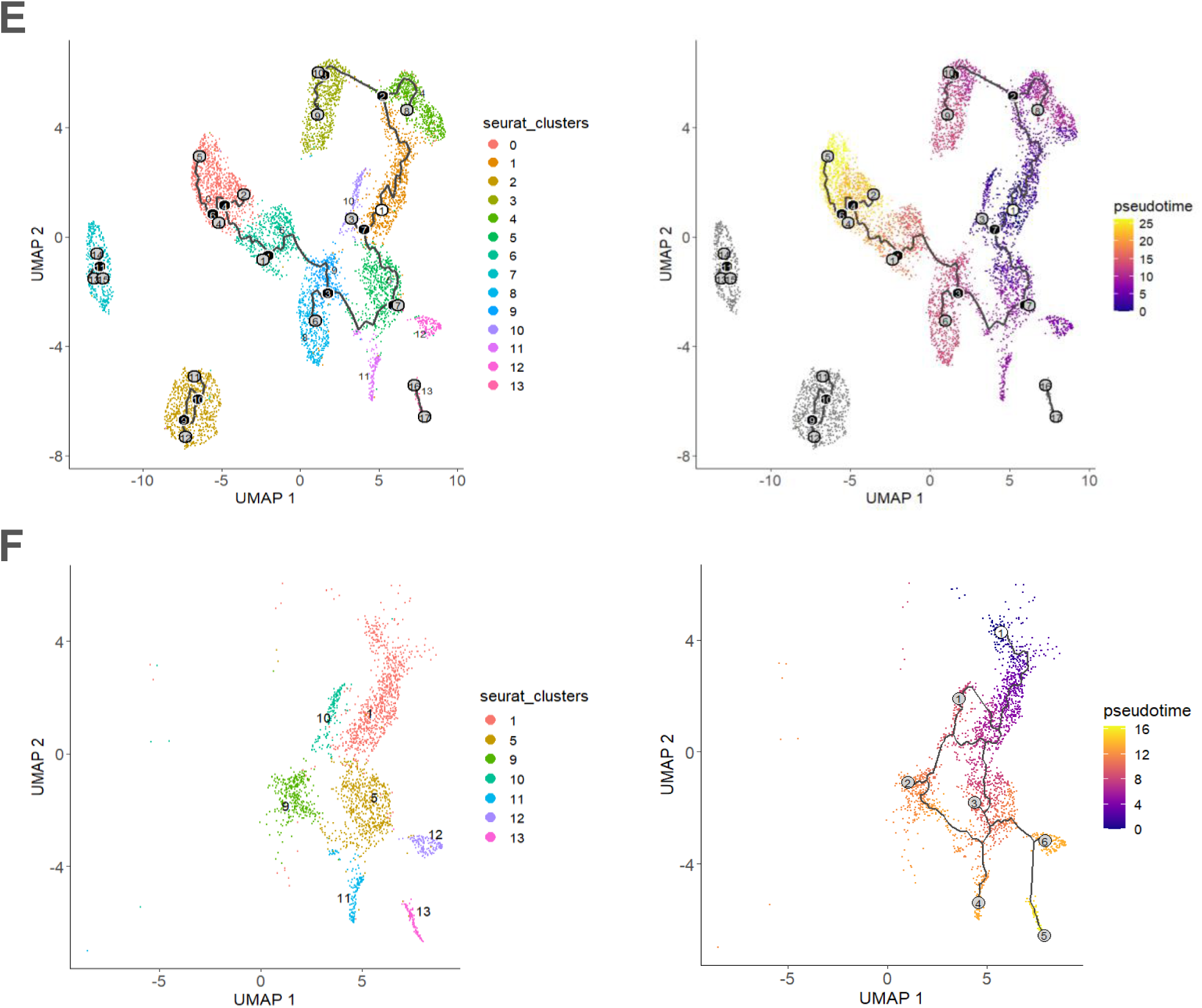
Differentiation trajectory of Kranz anatomy in a P4 maize primodium. **A and B.** Simulation of the successive differentiation trajectory of Kranz cell over pseudo-time (generated by “Monocle 2”), with “Start” indicating the beginning of the pseudo-time axis. Pseudo-time increases along the differentiation trajectory. The horizontal and vertical coordinates are two principal components, and different dots represent different cells, with different colors representing the cluster identity (A) or the pseudotime (B). Color from darkest to lightest blue in (B) represents pseudotime from beginning to end. The circled numbers represent different branch points. **C.** The split pseudo-temporal distribution of different cell clusters showing that the middle ground meristem (1) and kranz procambium cells (5) are at the early stage of differentiation and gradually differentiate into phloem (11) and xylem (12). **D.** Kinetics plot showing relative expression of representative genes from different clusters across developmental pseudotime. The abscissa represents the quasi-chronological order, and the ordinate represents the relative expression value of genes. The black line denotes the smoothed average expression. The colored dots on the top shows the major cluster of contributing cells to the expression, and the underneath stripes of cells contains all the clusters with different levels of expression. **E and F.** The successive differentiation trajectory of the whole (E) and a subset (F) of the 14 cell clusters over pseudo-time generated by “Monocle 3”. Dots, individual cells; cell clusters are coloured differently and labelled with numbers (refer to the circled numbers in Figure 4A). The white colour circled number “1” on cluster 1 indicates the beginning of the pseudo-time. Colour gradient on the right represents increasing pseudotime from beginning to end. The grey colour circled numbers represent different branches along the pseudotime trajectory, and the black colour circled numbers indicated the points where cell fate differentiates.

### Projection of single-nucleus transcriptome to segmented primordium RNA-seq — clusters 1 and 5 form the key group of cells for Kranz initiation

Due to the developmental gradient that exists between the tip, middle, and base of the primordium, bundle sheath cells, mesophyll cells and mGM display progressive stages of cellular differentiation from base to tip. For example, many intermediate veins had been initiated in the middle of the primordium whereas more undifferentiated mGM cells were located in the base of leaf primordium **(Figures 1D-1F)**. As part of the data verification, we performed a comparative analysis between the bulk RNA-seq data from segmented primordia and the snRNA-seq data. On the one hand, different tissue segments should be associated with cell clusters of different differentiation states in single-cell data. On the other hand, candidate Kranz regulators screened based on comparison of segmented primordium may be more enriched in the mGM and procambium cell clusters. The above hypothesis was confirmed by the pair-wise comparison between differentially expressed genes in segmented primordia and enriched genes in each of the 14 clusters from the single-nucleus transcriptome **(Figure 10)**.

Genes with higher expression in M2base than M2top, M3base than M3tip, and M3base than M3middle were more concentrated in clusters 8/9 (parenchyma cells), which was consistent with the phenomenon that parenchyma cells were mainly distributed around the midrib in the base of the primordium. The genes with higher expression in M3tip than M3base, M3tip than M3middle, and M3middle than M3base were enriched in clusters 11/12/13 (vascular tissue cells), corresponding to the higher degree of vein development toward the tip. The distribution frequency of these DEGs are marked by the dotted boxes in **Figure 10A**. Importantly, the genes with higher expression in M2top than M2base, and M3middle than M3tip were more concentrated in clusters 1 and 5 (mGM and procambial cells), and the 224 candidate genes screened from the segmented primordium RNA-seq were more enriched in cluster 5 (procambial cells). This is consistent with the hypothesis that the expression of the Kranz anatomy regulatory genes should be up-regulated in the mGM and procambial cells in the middle or upper middle part of the leaf primordium. The distribution frequency of these DEGs are marked by the solid boxes in **Figure 10A**. **Figure 10B** shows heat map grouping of the expression patterns of the boxed genes within the segmented primordia. Dot diagram in **Figure 10C** shows the expression patterns of the boxed genes among different cell clusters.

### Pseudo-time trajectory analysis and prediction of gene regulatory networks during Kranz development

To further explore co-regulators along Kranz development in maize leaves, we took the mGM cells as starting cells and carried out a pseudo-time analysis of the vein related clusters 1/5/11/12/13 by Monocle2 **(Figure 11)**. The reconstructed trajectory showed that the whole cell population branched at junction 1 and differentiated toward phloem (cluster 11) and xylem (cluster 12). At junction 2, the trajectory exhibited another branching event, resulting in two smaller branches. One of these eventually led to cluster 12, while the other remained closely associated with cluster 11. It should be noted that cluster 5 cells were more closely associated with phloem and xylem clusters than cluster 1, while cluster 1 cells were mainly aggregated at the early stage of pseudo-time **(Figures 11A-11C)**. This observation is in accordance with their respective temporal properties, with cluster 1 representing “mGM” and cluster 5 representing “procambial initial + procambial cell”. Cluster 13 which represents other vascular cells covers part of the early trajectory. We also tried Monocle3 with the successive differentiation trajectory overviewed for the whole 14 cell clusters along pseudo-time. A subset of clusters 1/5/11/12/13 plus clusters 10 and 9 showed that the differentiation trajectory of cell clusters is in principle similar to Monocle2 results, while their pseudo-time developmental relationships were better resolved compared to the whole set **(Figures 11E and 11F)**.

Dynamic gene expression patterns were inferred from the pseudo-time trajectories **(Figure 11D)**. *ZmEREB114* and *ZmEREB41* displayed high expression at the beginning (mostly contributed by cluster 1 cells), then slightly decreased before peaking (expression contribution from cluster 5 cells) at the middle stage and then continuously declining to lower or no-expression. By contrast *ZmEREB161* was initially expressed at a lower level but then also peaked after the middle stage (mostly contributed by cluster 5 cells), followed by continuous decline. *ZmPIN1,* however, showed very low expression at the beginning, then gradually increased throughout the pseudo-time trajectory. The expression of phloem system marker gene *ZmCALS2* (Zhong et al., 2023) was also lower at the early stage but increased and peaked after middle stage, while another marker gene homolog of *AtNEN3* displayed a prominent peak erupted from very low base level (contribution of both mostly from cluster 11 cells) **(Figure 11D; Supplemental Figures 8A and 8B)**. Xylem related genes including homolog of *AtLOG6* and *ZmWAT1* (Xu et al., 2021) were not expressed at the early stage, but expression levels increased steeply at later stages of pseudo-time (the contribution of expression was clearly from cluster 12) **(Figure 11D; Supplemental Figures 8C and 8D)**.

Given that cluster 5 cells appeared along most of the pseudo-time developmental track, we took these cells for gene co-expression analysis to understand how different gene cohorts were represented at this transitional stage. To this end, we selected regulatory genes from clusters 1/5/11/12/13, and after co-expression analysis with cluster 5 cells, a network of 33 genes were obtained. The network included ARF, AP2 family transcription factor and other auxin related genes, with many of them also showing protein interaction relationships **(Figures 12A and 12B)**. The expression patterns of 33 genes, plus *ZmSHR1*, *ZmSCR1*, *ZmSCR2*, *ZmTML1* and *ZmEREB184* along pseudotime are shown in the heatmap of **Figure 12C**. It can be seen that to a certain extent hierarchical regulation corresponds to the pseudo-time expression pattern: *ZmARF25*, *ZmEREB114*, and *ZmEREB41* are expressed first; *ZmSHR1/2* and *ZmEREB161* are next; followed by another 14 genes (including *ZmTUB4*, *ZmPIN4*, *ZmEREB200*, *ZmMYB121*, *ZmATL1*, *ZmSBP16*, *ZmRLD1*, *ZmNAC131*, *ZmIAA45*, *ZmbHLH63*, *ZmBIF1*, *ZmGATA4*, *ZmIAA4*, and *ZmLBD26*). A schematic model combining *in situ* arrangement of cell clusters and distribution of potential regulators of Kranz organization, is presented along a pseudo-time developmental axis in **Figure 12D**.

**Figure 12.**
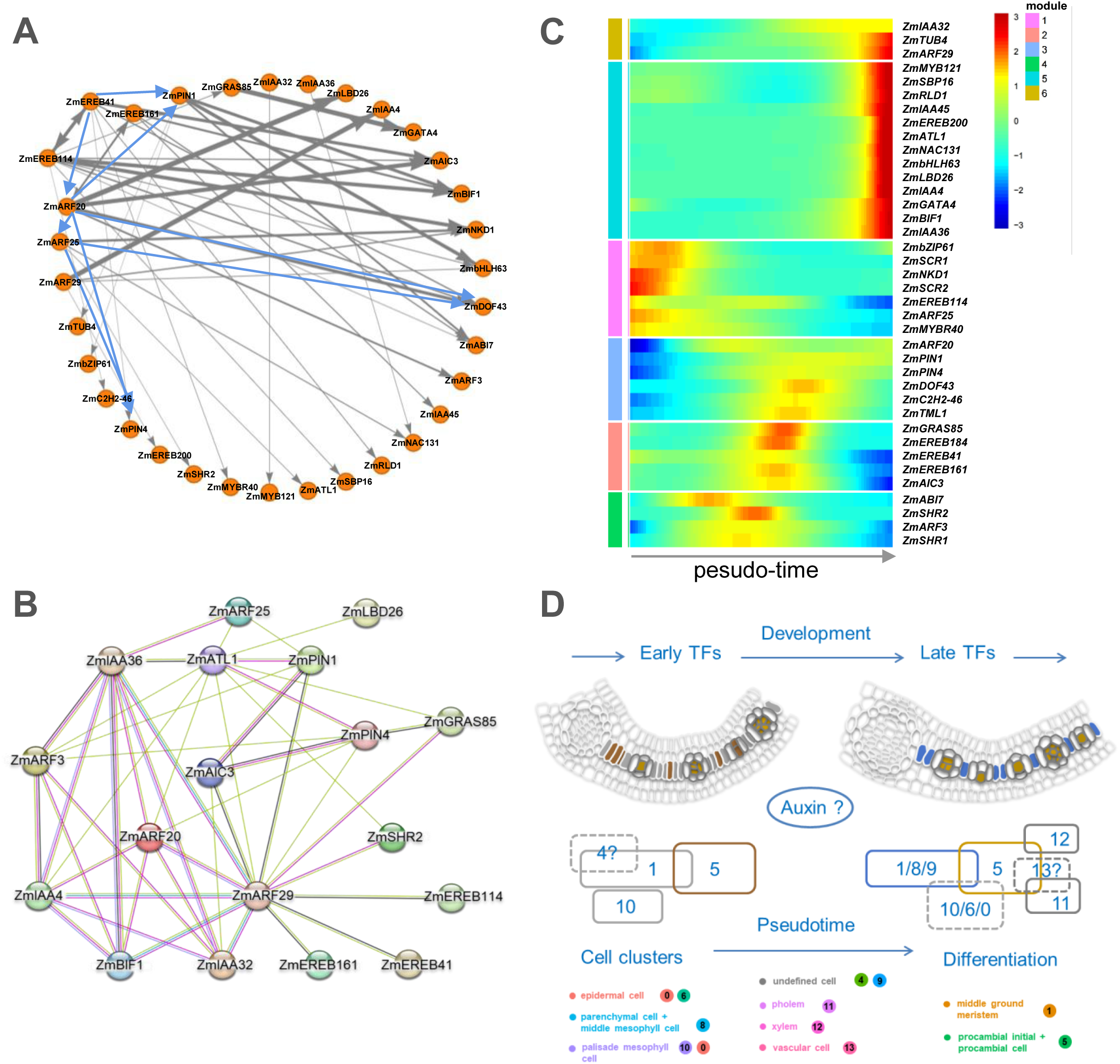
Predicted gene regulation network associated with the process of Kranz cell differentiation. **A.** The regulatory network of 33 TF and auxin related genes from the co-expression network of clusters 1,5,11,12, and 13. Gray lines denote connections of co-expression inferred by WGCNA. The thickness of the gray line represents the level of correlation. The blue lines denote the TF binding validated by EMSA in Liu et al. (2022). **B.** The protein-protein interaction network of 18 TF and auxin related proteins (hand selected with highlight on auxin regulation) based on String database. Strings: blue, interaction relationship from curated databases; megenta, experimentally determined; green, gene neighborhood; red, gene fusions; dark blue, gene co-occurrence; yellow green, textmining; black, co-expression; light purple, protein homology. **C.** Heatmap showing the six gene modules (1–6) of 33 genes, plus *ZmSHR1*, *ZmSCR1*, *ZmSCR2*, *ZmEREB184*, and *ZmTML1* along pseudotime. The list of 33 genes is given in Supplemental dataset 7. The list and heatmap of 475 top significant DEGs along pseudotime are given in Supplemental dataset 6. **D.** Schematic model combining cell clusters, developmental trajectory, and potential regulators of Kranz anatomy. Clusters 4, 13 and 6/0 are drawn with dashed lines, meaning their locations or identities are under further investigation.

## Discussion

Compared with the C_3_ plant rice, the increased number of intermediate veins leads to high vein density in maize leaves, which is an important contributing factor for the formation of efficient C_4_ leaves. As for the development of typical Kranz anatomy in maize leaves, the key scientific questions pending to be answered include: (1) How do the intermediate veins in maize leaves initiate? (2) What regulatory factors are required, and in what temporal-spatial manner, to regulate the extension and assembly of the intermediate veins?

### Primordium sub-section facilitates identifying Kranz-propagation zone and multi-level comparative transcriptomics

After realizing the limitations of working at a tissue-level, we have moved our focus to the active initiation/proliferation regions inside single C_4_ leaf primordia. Combining fluorescent labelling and sequential sectioning, we were able to analyse vein development patterns in maize P4 leaf primordia in proximal-distal (PD), medial-lateral (ML), and adaxial-abaxial contexts **(Figure 1)**. This allowed us to capture the active zone within the leaf where the initiation, and extension of intermediate veins was expected to be concentrated. It laid a solid foundation for us to draw systematic cell arrangement maps at the tissue and cell levels, and to separate active Kranz development locations for comparative analysis of micro-tissue and single-cell level transcriptomes.

We obtained gene expression signatures for sub-sectioned leaf primordia of maize and rice, contributing to the early leaf primordium development database at a sub-tissue level, which will facilitate further investigation of monocotyledonous leaf development. For example, photosynthetic competence was previously reported to be established in rice at the P3/P4 transition stage, and chlorophyll fluorescence signal was restricted to the distal tip of the P4 stage leaves (van Campen et al., 2016). Correspondingly, in maize primordia, M3tip enriched biological functions were mostly related to photosynthetic processes **(Figure 2C)**. These sub-tissue transcriptomic profiles should help to shed light on early photosynthetic development of grass leaves. We were also able to perform comparative analysis of early Kranz development with species comparisons between C_4_ vs C_3_ (maize vs rice), and tissue comparisons between Kranz vs non-Kranz (M3middle vs Msheath), which was a significant extension to the relevant database (Wang et al., 2013; Wang et al., 2014).

### SnRNA-seq identifies cell heterogeneity and clusters responsible for intermediate vein initiation in maize leaf primordia

Single-cell RNA sequencing (scRNA-seq) technology has revolutionized molecular and cellular biology by improving the spatiotemporal resolution of transcriptomic analysis to the level of individual cells, which is rapidly expanding our ability to elucidate cell types, states, origins and differentiation (Seyfferth et al., 2021). However, scRNA-seq analysis in maize leaf primordia faces the challenge of obtaining high quality protoplasts. We therefore tested the use of isolated nuclei for transcriptome sequencing, as single-nucleus transcriptomes of various animal and plant systems has provided biologically meaningful information (Bakken et al., 2018; Farmer et al., 2021). Although some cytoplasmic message may be missed, but the extraction of nucleus helps to better capture RNAs from cell types suffering from limited protoplast integrity (vascular cells for example). Our single-cell resolution transcription map revealed high cell heterogeneity in early maize leaf primordia, showing continuity of the cell developmental trajectory during initial Kranz-type vein formation. In the future, single-nucleus ATAC-seq (SnATAC-seq) or spatial transcriptomic technology (Farmer et al., 2021; Xia et al., 2022; Perico et al., 2024) may assist to gain deeper or wider understanding of the regulatory mechanisms controlling Kranz anatomy formation.

Cell clustering and annotation are other challenges for the single-cell analysis of P4 leaf primordia of interest in this study. First, there are few known cell type-specific marker genes in maize primordia. Second, the sample is restricted to a specific leaf primordium which lacks single-cell data for reference. For the single-cell data of maize SAM+P1+P2 (Satterlee et al., 2020), the clustering of cell types spans information among different tissues, while our samples need to be subdivided into cell types within the P4 leaf primordium. Therefore, *in situ* hybridization was heavily relied on for subsequent annotation and validation. We retrieved 7 broad populations of cells with 14 transcriptionally distinct clusters, representing strong cell heterogeneity in maize leaf primordia during this critical period of C_4_ Kranz anatomy development. Importantly, we identified the cell groups (clusters 1/5) involved in giving rise to the intermediate veins **(Figures 4-7)**. The projection of segmented primordium transcriptome to single-nucleus transcriptome supports that cluster 1/5 cells initiate the core components of Kranz anatomy **(Figure 10)**. The pseudo-time trajectory analysis again showed that cluster 1/5 cells include transitional cells, which effectively linked multiple developmental stages and branches such as the mGM that had not yet begun differentiation, the procambial initial cells, and the early differentiating vascular tissues **(Figures 11)**. However, cell groups representing the bundle sheath cells remains to be further confirmed. Due to limitations imposed by sequencing depth and gene coverage, lowly expressed genes which could be specifically expressed in bundle sheath cells, may not be effectively represented in the current snRNA-seq datasets, nor be effectively picked out by *in situ* hybridizations.

### Insights in the Kranz organization from middle ground meristem by networking and mutation analysis

A role for auxin in vascular tissue formation was identified many years ago. Understanding the spatial and temporal patterns of auxin activity within and between tissues or cells is required to elucidate the role of auxin in this process (Perico et al., 2022). According to our analysis, the enrichment of auxin-related genes during Kranz-type vein formation mainly included auxin transporter (including *ZmAIC3, ZmPIN1,* and *ZmPIN4*) and auxin signaling pathway (such as *ZmARF29*, *ZmARF25*, *ZmARF20, ZmARF3,* and several *ZmIAAs*) genes **(Figure 12A)**. *ZmPIN1* is expressed in pre-procambial, procambial initial, and procambial cells, across the development stages (Robil and McSteen, 2023), showing auxin as a major positional signal. Inspired by the recent advances in the role of auxin in vein patterning (Liu et al., 2023; Dong et al., 2023), the genes of Aux/IAA and ARF families we identified here are worth future investigation.

Robil and McSteen (2023) analyzed the expression of DR5, TCS, and GAR2 reporters to investigate the roles of auxin, cytokinin (CK), and gibberellic acid (GA) respectively in developing maize leaf primordia. They presented evidence that auxin efflux precedes the CK response in procambial strand development. We have collected about 83 genes related to the CK signalling pathway, and found that the ones expressed in our snRNA-seq data were relatively enriched in clusters 11, 12 and 13 **(Supplemental Figure 12A)**. This is consistent with the relatively late response of CK during vein formation. Among the genes enriched in clusters 11, 12 and 13, there is a group of 5 genes which exhibited increased expression from M3base toward M3tip, as well as from M2base toward M2top, but not in Msheath **(Supplemental Figure 12B)**. Due to the blade preference of these DEGs, they are pending to be tested for possible roles in intermediate vein development.

Notably as well, AP2 family transcription factor genes *ZmEREB114*, *ZmEREB41* and *ZmEREB161* were identified and their expression location were verified **(Figures 5-7)**. We compared the expression patterns of *ZmEREB161* and *ZmSHR1*, and found that *ZmSHR1* was expressed in dividing procambial centres, similar to the pattern observed in rice primordia (Liu et al., 2023). By contrast *ZmEREB161* was expressed not only in the dividing procambium, but also more strongly in single procambial initials recently specified from the mGM cells **(Figure 7)**. The expression of the rice homologue of *OsEREB161* was restricted to procambial centres, without clear signs of expression in cells of the middle layer in between existing veins **(Supplemental Figure 13)**. Therefore, *EREB161* might be more related than *SHR1* to pre-Kranz events in maize leaves, which is also supported by their relative enrichment in intermediate veins and lateral veins respectively in recent spatial transcriptomics of P4 primordium (Perico et al., 2024).

Arabidopsis has a single *AINTEGUMENTA1* (*ANT1*) gene, while maize has four (*ZmANT1-4*, corresponding to *ZmEREB184*, *114*, *41*, *161* respectively), and setaria and rice have three. Liu et al. (2020) observed mild and sporadically occurring defects in *Svant1* mutant (mutation of the homolog of *ZmEREB184*) of *S. viridis*, and suggested that the lack of perturbations might be attributed to the existence of two other *ANT* genes in the genome. To determine the function of *ZmEREB161*, we first obtained maize mutants (point mutation and premature stop codon) of *ZmEREB161*, and characterized the phenotype of lines encoding the truncated protein. Although some instances of directly adjacent veins and supernumerary bundle sheath cells were found, vein density was not statistically significant between mutant and wild type leaves **(Supplemental Figures 14A-14C)**. Similarly, vein density was not altered in rice when *ZmEREB161* was constitutively expressed (*ZmUBIpro::ZmEREB161)* but exhibited reduced plant height, leaf length and leaf width **(Supplemental Figures 14D-14H)**. However, by transforming *ZmEREB161pro::ZmEREB161* into Arabidopsis, increased vein density was observed, together with increased leaf area **(Supplemental Figure 15)**. It is possible that co-regulation of gene targets by different EREB (ANT) proteins may be important for venation patterning in grasses. The co-orthologs of Arabidopsis ANT1 in maize (ZmEREB 161, 41, 114, and 184) may work in combination during Kranz development. More in-depth and systematic characterization of mutants in maize or different C_4_ grasses is necessary for the mGM and procambium related candidate genes. Promisingly, *TOO MANY LATERALS* (*ZmTML1*), one of the 3 transcription factor encoding genes from the 10 candidates we highlighted in **Figure 3C**, was recently shown to specify vein rank in maize leaves (Vlad et al., 2024).

### Comparison between snRNA-seq and the transcriptome of laser capture microdissection (LCM) isolated cells of maize embryonic leaf

The work by Liu et al. (2022) used LCM-dissected maize embryonic leaf primordia to generate transcriptomes of different stages of pre-Bundle sheath cells and pre-Mesophyll cells. Regulators related to Kranz anatomy and/or vascular development were suggested from gene expression analysis of the middle ground meristem (mGM) cells. It is suggested that the differentiation of pre-Kranz cells occurs largely before the three-contiguous cell (3C) stage, whereas rapid differentiation of pre-Mesophyll cells continues until the four-prepalisade mesophyll cell (4PM) stage. Importantly, by comparing between Liu et al. (2022) and our current work, genes that are highly expressed in mGM and 3C from the LCM-transcriptomes (Fig. 7A of Liu et al., 2022) are also partly enriched in clusters 1 and 5, and enriched to less extent in clusters 11 and 12 in our snRNA-seq data **(Supplemental Figure 16A)**.

To further show that this overlap is more significant than random chance, we have performed further grouping analysis taking into account of both the LCM data and our snRNAseq clusters **(Supplemental Figure 16B)**. The sections framed in blue lines represent the selected LCM data modules overlapping with our snRNAseq clusters. Importantly, we think these two studies support each other not only in a way that they overlap to certain extent, but also in a way that they compensate or help to adjust each other’s uncertainties. For example, despite chances of unavoidable contamination between samples, the “PM” and “2M” samples of the LCM data are being helpful in assisting us to further determine different types of mesophyll cells for the single-cell clusters **(Supplemental Figure 17)**. Accordingly, our clusters 0 and 6 were suggested to contain palisade mesophyll cells apart from epidermal cells, and cluster 8 was identified to be related with middle layer mesophyll in addition to parenchyma cells. Looking further ahead, together with new data set such as spatial transcriptomics (Perico et al., 2024), researchers in the field will benefit from more comprehensive resources for dissecting C4 leaf Kranz anatomy.

In summary, using a combination of cellular, developmental and system biology approaches, this study provides both theoretical and practical platforms for the exploration and verification of Kranz anatomy regulators and their spatial-temporal combinations. We proposed a group of potential mGM derived or procambium localized regulators associated with Kranz initiation and development, and provided single-cell resolution resources for studying the early division and differentiation of Kranz cells in maize leaves. Further studies on the regulatory functions and the potential application on C_4_ leaf engineering need to be conducted using relevant genetic materials.

## Materials and methods

### Plant material and growth conditions

In Shanghai, *Zea mays* (L.) ssp. Mays cv. B73 and *pZmPIN1a::ZmPIN1a:YFP* transgenic line (Yang et al., 2015), as well as *Oryza sativa* (L.) ssp. Japonica cv. Nipponbare and *DR5::VENUS* line, were grown in a phytotron, 27 °C in day and 25 °C at night, with 600 µmol photons m^−2^ s^−1^ light intensity, 16 h light and 8 h dark photoperiod. Tissue was harvested from 2-, 3- or 4-week-old maize plants and 2-week-old rice plants.

*Zmereb161* mutant of maize by EMS (Ethyl methanesulfonate) chemical mutation was obtained from http://www.elabcaas.cn/memd/index.php. The M3-generation was used and genotyping was conducted by genomic PCR with primers listed in **Supplemental Dataset 8**. PCR products were sequenced to confirm the mutation of *zmereb161*, in which the peptide chain coding is terminated prematurely from CAG to TAG.

To generate *ZmUBIpro::ZmEREB161* transgenic plants in rice, coding sequence of *ZmEREB161* was PCR amplified from maize cDNA that had been generated using RNA isolated from P1-5 leaf primordia. The amplified sequence was subcloned into Gateway® donor vector, sequenced, and then cloned into destination vector pSC310, downstream of the maize ubiquitin promoter. The construct was transformed into japonica rice cultivar Nipponbare, and after T1 genotyping and expression analysis, two lines of *ZmUBIpro::ZmEREB161-15* and *ZmUBIpro::ZmEREB161-16* seedlings were used for phenotypic analysis. Primers used are listed in **Supplemental Dataset 8**

*Arabidopsis thaliana* cv. Columbia (Col-0) identified to be heterozygous for a T-DNA insertion in *AtSHR1* (mutant line *shr-6*, Yu et al., 2010) (NASC) was used for transformation of a *ZmEREB161pro::ZmEREB161* construct, but only the non-mutant *SHR/SHR* progeny were characterized. The *ZmEREB161pro* promoter consisted of a 2960 bp region upstream of the start codon and was used instead of a constitutive promoter to reduce the risk of sterile transgenic plants. After T2 genotyping and expression analysis to exclude lines with gene silencing, two lines (*ZmEREB161pro::ZmEREB161-3* and *ZmEREB161pro::ZmEREB161-4*) expressing *ZmEREB161* were used for phenotypic analysis. Plants were grown in a glasshouse at 21°C with 16 h of supplemental LED light.

Leaf 5 from 21 day old arabidopsis plants was cleared with chloral hydrate solution (100% w/v chloral hydrate, 1% v/v glycerol) for 48 h, mounted on glass slides using 50% glycerol and sealed with nail varnish. The leaves were imaged using darkfield photography with a Nikon D300 camera, and vein patterning analysed using LIMANI (Leaf Image Analysis Interface) software (Dhondt et al., 2012).

### ClearSee and confocal imaging

ClearSee assays were performed as previously described (Kurihara et al., 2015). In brief, leaf primordium were immersed in fixation buffer (4% w/v paraformaldehyde, Cat. No. AR-0211) under vacuum at 25 mbar for 1h, washed once with 1× PBS, and then immersed in ClearSee reagent (10% xylitol, 15% sodium deoxycholate, and 25% urea) for 3 days. The primordium were flattened with the adaxial side facing up in ClearSee reagent, and observed under a ZEISS LSM880 confocal microscope using a 514-nm laser excitation and 520-to 560-nm emission for detection of the YFP.

### Histology

Leaf primordium samples were fixed overnight in ethanol/acetic acid (3:1), and embedded in Paraplast Plus (Sigma, Cat. No. P3683-1KG) using a modular automated tissue processor (KD-TS3A and KD-BM/BL). Paraffin-embedded samples were sectioned (5 μm) with a Rotary microtome (Lecia, RM2125). Sections not dewaxed were viewed using an Olympus CX23 microscope.

### *In situ* hybridization and imaging

For probe synthesis, templates of RNA probes were amplified from cDNAs using gene-specific primers containing T7 promoter sequences at the 5’ end. *In vitro* transcription was performed with T7 RNA polymerase (Roche, Cat. No. 10881767001; Thermo, Cat. No. EP0111) and DIG RNA labeling mix (Roche, Cat. No. 11277073910). The primers used to generate the probes are listed in **Supplemental dataset 8**.

Tissue embedding and RNA *in situ* hybridization were performed as described with modifications (Langdale 1994; Zeng et al, 2021; Weigel et al., 2002). Briefly, leaf primordium were fixed with FAA (3.7% formaldehyde, 5% glacial acetic acid, and 50% ethanol), embedded in paraffin and sectioned (5 μm) as described above. The sections were dewaxed, digested with proteinase K (Sigma, Cat. No. P6556), dehydrated with gradient ethanol, and hybridized with RNA probes. After washing, the sections were incubated with anti-digoxigenin-AP Fab fragments (Roche, Cat. No. 11093274910). The signals were developed with the NBT/BCIP stock solution (Roche, Cat. No. 11681451001), and the sections were imaged using an Olympus CX23/BX3-CBH microscope. Where necessary, the sections were also stained with 10 mg/mL Calcofluor white for enhancing UV excited cell wall autofluorescence, and imaged with a fluorescence microscope (DM6000B, Leica).

### Transmission electron microscopy (TEM)

Maize leaf primordium was fixed in 2.5% (w/v) glutaraldehyde overnight at 4°C. Post fixation in osmium tetroxide, embedding in Spurr’s resin, and other steps were performed by the TEM platform in College of Life and Environmental Science, Shanghai Normal University, following standard procedures. The ultra-thin sections were imaged under a transmission electron microscope (Tecnai Spirit G2 BioTWIN, FEI) using a voltage of 120 kV.

### Sample collection for bulk RNA-seq

Maize leaf primordium of about 5 mm were partitioned under dissection microscope into three parts: M3tip≈1.5 mm, M3middle≈2.5 mm, and M3base≈1 mm. Maize leaf primordium of about 3 mm were partitioned into two parts: M2top≈2 mm and M2base≈1 mm. About 200 leaf primordium and 10 leaf sheath of 1-2 mm were obtained, partitioned, and pooled into different samples with three biological replicates. The rice leaf primordium of about 5 mm were also partitioned into R3tip≈1.5 mm, R3middle≈2.5 mm, and R3base≈1 mm. Similarly, about 400 leaf primordium and 20 leaf sheath of 1-2 mm were processed into different samples with three biological replicates. Total RNA was extracted using the GeneJET Plant RNA Purification Mini Kit (Thermo Scientific, Cat. No. K0801) according to the manufacturer’s instructions.

### RNA-seq

Six types of maize tissues and 4 types of rice tissues were sampled as described above for bulk RNA sequencing. RNA-seq was performed using the Illumina NovaSeq 6000 platform. Reads were mapped to the genome version of *Zea mays* V4. Differentially expressed genes (DEGs) were determined by DESeq2 (Love et al., 2014). Three biological replicates were used in the data analyses. Pearson correlation coefficients were calculated by function “cor” in the R package “stats” and visualized by function “pheatmap” in the R package “pheatmap”.

### Gene Ontology (GO) analysis

The Plant Regulomics database was used to analyze the GO functions of differentially expressed genes and FDR<0.05 was used as the selection condition (Ran et al., 2020). The ggplot R package was used to visualize the enrichment results.

### Sample collection and preparation of nucleus

According to our pilot experiments, ∼200 maize leaf primordium could yield enough high quality nuclei, using the methods previously described with modifications (Thibivilliers et al., 2020; Conde et al., 2021). To isolate nuclei, freshly collected leaf primordium of 3-4 mm (with group efforts to ensure enough amount within 1 h) were placed on Petri dishes and vigorously chopped with two razor blades for 5 minutes in ∼500 μL Nuclei Isolation Buffer [NIB, 10 mM MES-KOH pH 5.7, 10 mM NaCl, 10 mM KCl, 2.5 mM EDTA, 250 mM sucrose, 0.1% protease inhibitor cocktail (Roche), and 0.1% BSA (YESEN), finally adjust pH to 5.7]. Homogenized tissue was first filtered with 40 μm cell strainers (Falcon, Cat. No. 352340), then filtered with 30 μm cell strainers (pluriSelect, Cat. No. 43-50030-01). Nuclei were spun down in a swinging-bucket centrifuge (5 min, 500 rcf.) and re-suspended in 100 μL washing buffer (15 mM Tris-HCl pH 7.5, 160 mM KCl, 40 mM NaCl, finally adjust pH to 7.0). Nuclei quantification was done by staining with trypan blue (final concentration of ∼0.1%), and counting on a hemocytometer for a total of 200,000 nuclei. Nuclei suspensions were then spun down (5 min, 500 rcf.) and re-suspended in diluted nuclei buffer (10x Genomics) to a final concentration of 2,000∼2,500 nuclei per μL, and used as input for snRNA-seq library preparation (∼16,000 nuclei in total). Samples were kept on ice for all intermittent steps.

### SnRNA-seq library construction

Approximately 16,000 counted nuclei were loaded on Single Cell A Chip. The libraries were constructed using a Chromium Controller and Chromium Single Cell 30 Reagent Kits V3.1. Qualitative analysis of DNA library was performed by an Agilent 2100 Bioanalyzer. Libraries were sequenced by an Illumina NovaSeq 6000.

### SnRNA-seq data processing

The raw scRNA-seq dataset was first analyzed by Cell Ranger 7.0.0 (10x Genomics), to align reads and generate gene-cell matrices. The genome (V4) and GTF annotation files (Zm-B73-REFERENCE-GRAMENE-4.0) of Zea mays were downloaded from the Ensembl plant web-site.

The snRNA-seq data (10x Genomics Cell Ranger output) was corrected for ambient RNA expression using SoupX (v1.5.0) (Young et al., 2020). SoupX was run with clustering information derived from a generic processing workflow in Seurat. Seurat v.4.2.1 (https://satijalab.org/seurat/) was used for filtering the data (Germain et al., 2021), reducing dimensions, clustering cells and identifying DEGs, which were implemented in R (v. 4.1.0). The UMI expression matrix with genes expressed in at least 3 cells and cells expressing at least 200 genes was loaded into Seurat. Further cleaning steps were performed using log10GenesPerUMI, and set the log10 fold change of the proportion of number of genes per read counts more than 0.7 and mitochondria percentage below 5%. We also used scDblFinder (v1.2.0) software for detection and handling of doublets/multiplets in the snRNA-seq data (Hao et al., 2021). A total of 7,473 cells with average of 2,424 genes after filtering were used for downstream analysis. The SCTransform function with the default parameters was used for data normalization, scaling and transformation. The dimensions of the expression matrix were then reduced by the RunPCA function, and the top 30 dimensions were used for FindNeighbors and UMAP analysis. The cell clusters were identified by the FindClusters function with a resolution of 1. Calculations for the DEGs were conducted using the FindAllMarkers function with the default parameters.

Pseudo-time trajectory analysis of snRNA-seq data was conducted using Monocle 2 (Qiu et al., 2017) or Monocle 3 (Lee et al., 2021). Gene expression was plotted in Monocle 2 to track changes across pseudo-time. We also plotted TFs and marker genes along the inferred developmental pseudo-time.

### SnRNA-seq versus bulk RNA-seq of primordium

The overlap ratios of DEGs between different primordium partitions from bulk RNA-seq with marker genes in each cell cluster from snRNA-seq were obtained. The pheatmap R package was used to visualize the ratio matrix results.

### Construction of co-expression network

The regulation network was predicted by unsupervised learning method (GENIE3) based on the expression matrix of corresponding cell clusters, and was plotted by visNetwork according to the PlantTFDB database (Jin et al., 2017).

### Quantification and statistical analysis

One-way ANOVA with Tukey’s HSD test was used to determine the statistical significance among different samples. *P*-value < 0.05 is considered as statistical significance, and different letters on the bar graphs indicate statistically significant difference. All statistic results were generated by SPSS 22 and all graphs were generated by GraphPad Prism 8 (www.graphpad.com). The numbers of samples and types of statistical analyses are given in figure legends and the quantitative details can be found in Microsoft Excel spreadsheets of the **Supplemental Dataset 10**.

### Accession numbers

Accession numbers for the genes analyzed are listed in **Supplemental dataset 7 and 8**.

### Data availability

Single-cell and bulk RNA-seq data have been deposited at NCBI’s Sequence Read Archive (SRA) and are publicly available as of the date of publication. Microscopy data reported in this paper will be shared by the lead contact upon request.

## Author contributions

JY and PW conceived the project; JY, HS, YC and PW performed the experiments; OS and JAL worked on the *ZmEREB161* overexpression lines in rice and Arabidopsis; JY, ST, HS, OS, CZ, JY and PW analyzed the data and produced the figures; JY, YZ, XZ, JAL, JW and PW interpreted the results, wrote, and revised the paper.

## Acknowledgments

We thank Langdale lab for fruitful discussion. We thank Rui Zhang, Xiyu Zeng, Jiachun Wu, and Qiqi Zhang for the team work of dissecting and collection of maize and rice leaf primordium. We are grateful to Xiaotong Lv and Yutong Li for helping with figure generation and mutant analysis. We thank Prof. Fang Yang for providing the maize *pZmPIN1a::ZmPIN1a:YFP* transgenic line, and Prof. Dabing Zhang for providing the rice *DR5::VENUS* line. We are grateful to Dr. Hua Wang, Dr. Shuining Yin, and Dr. Naiying Yang for technical supports of *in situ* hybridization, ZEISS LSM880 confocal microscope, and transmission electron microscopy.

---

**Supplemental Figure 1.**
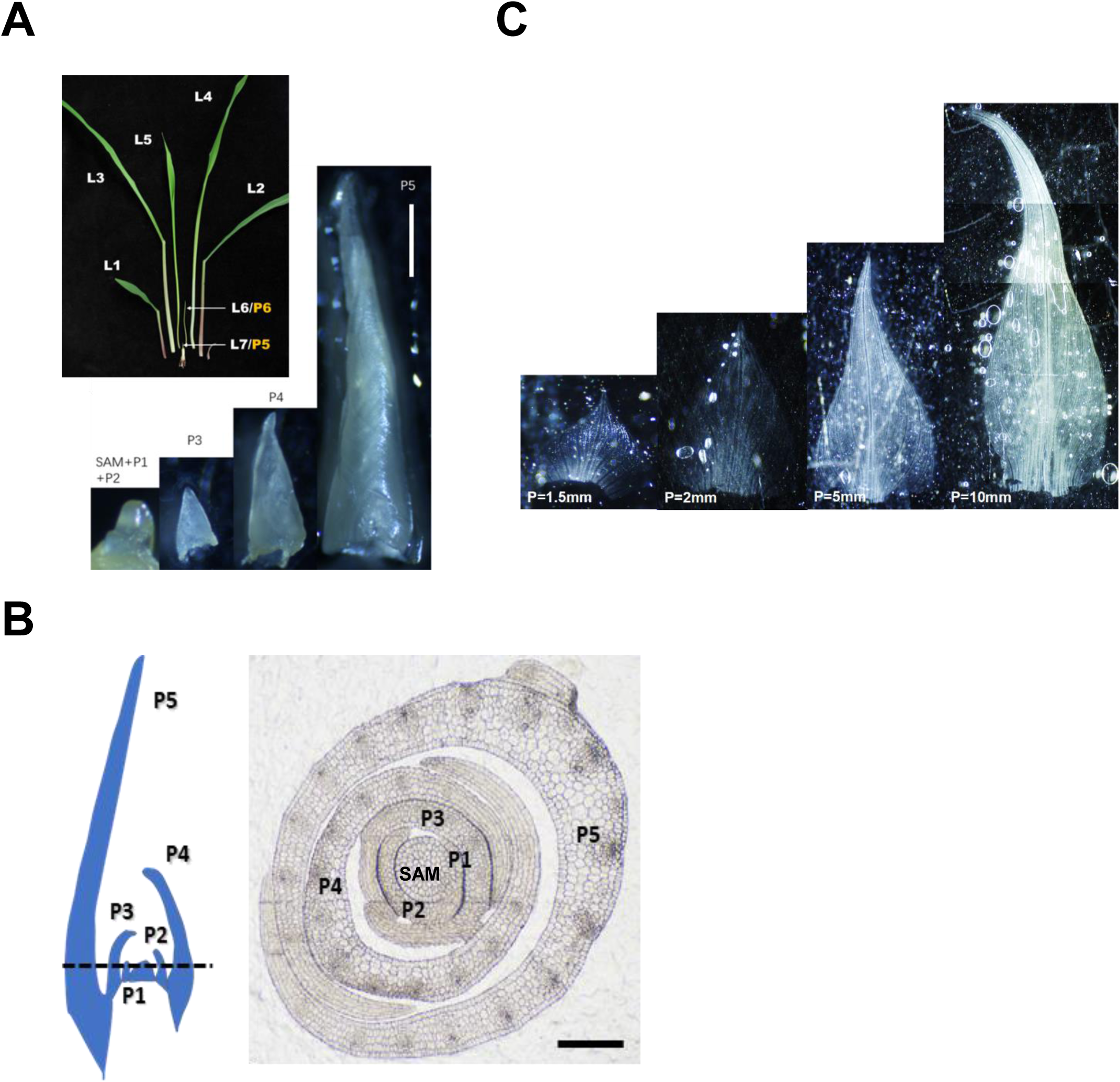
The growth pattern of maize leaf primordia. **A.** Leaf primordia are initiated at time intervals known as plastochrons (P), such that the youngest primordium (P1) is closest to the SAM, and older primordia (P2, P3, etc.) are consecutively further away. The first leaf to be produced after germination (and hence the oldest) is L1, and subsequent leaves are L2, L3, etc. Each leaf thus has a “P” number to denote relative developmental stage and an “L” number to denote age. Leaf blades are separated from leaf sheath tissue by the ligule, which is established around P5 (Want et al,. 2016). The image here shows the visible leaves: L1, L2, L3, L4, and L5. L6 is at P6 and L7 is at P5. Dissected leaf primordia are also shown: P1, P2, P3, P4, and P5. Scale bar: 1000 μm. **B.** Schematic on the left shows the longitudinal section of SAM plus five most recently initiated leaf primordia (P1-P5). Dashed line indicates the position of the transverse section shown on the right. Scale bar: 100 μm. **C.** The primodium of 1.5mm, 2.5mm, 5mm and 10mm in size are detached, unrolled, and flattented with the adaxial side facing up (P=1.5mm means the length of primordium is 1.5 mm).

**Supplemental Figure 2.**
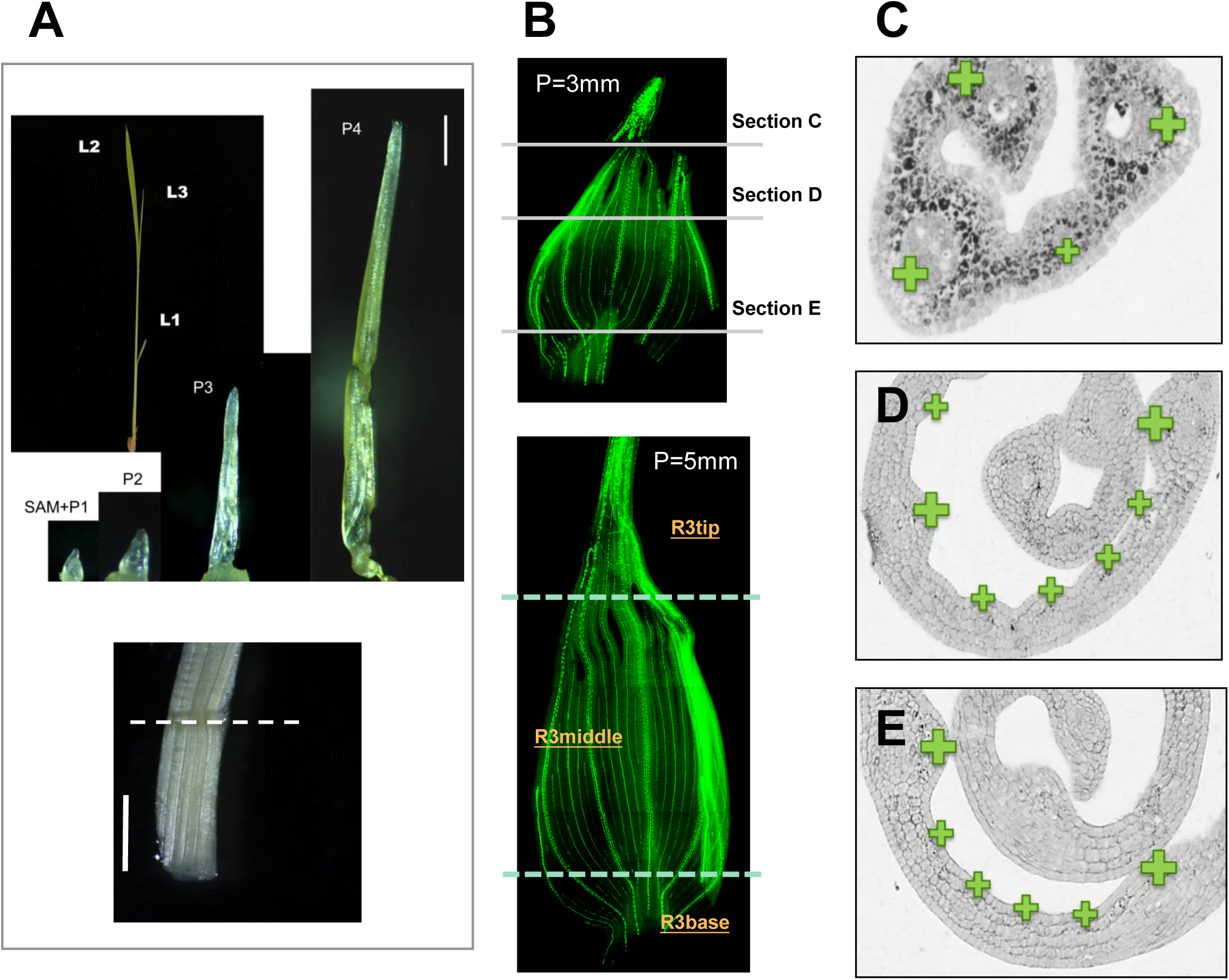
The growth pattern and vein development of rice leaf primordia. **A.** Upper: Images showing the visible leaves (L1-L3) and dissected leaf primordia (P1-P4) from rice seedlings. Note that P3 and P4 of rice are slimmer than those in maize. Scale bar: 500 μm. Lower: The leaf sheath of 1-2 mm, cut below the dashed line from P6. Scale bar: 1000 μm. Note: Dashed lines indicate the position where it was partitioned. **B.** The primodia P=3mm and P=5mm are excised from the base, unrolled, and flattented with the adaxial side facing up. Vein strands were visualized by DR5::VENUS. P=3mm means the length of primodium is 3 mm. Upper panel: Positions of cross sections made for images in C, D and E were indicated. Lower panel: Image indicating the rice tissues segmented for bulk RNA sequencing. The leaf primordium of about 5 mm were partitioned under dissection microscope into three parts: R3tip≈1.5 mm, R3middle≈2.5 mm, and R3base≈1 mm. Note: Dashed lines indicate the position where it was partitioned. **C-E.** Transverse sections showing vein patterning across the tip, upper middle, and lower middle of P =3mm rice leaf primordia. Larger and smaller green “+” mark larger and smaller lateral veins respectively, and indicate that the distribution of intermediate relative to lateral veins remained similar between middle and base sections.

**Supplemental Figure 3.**
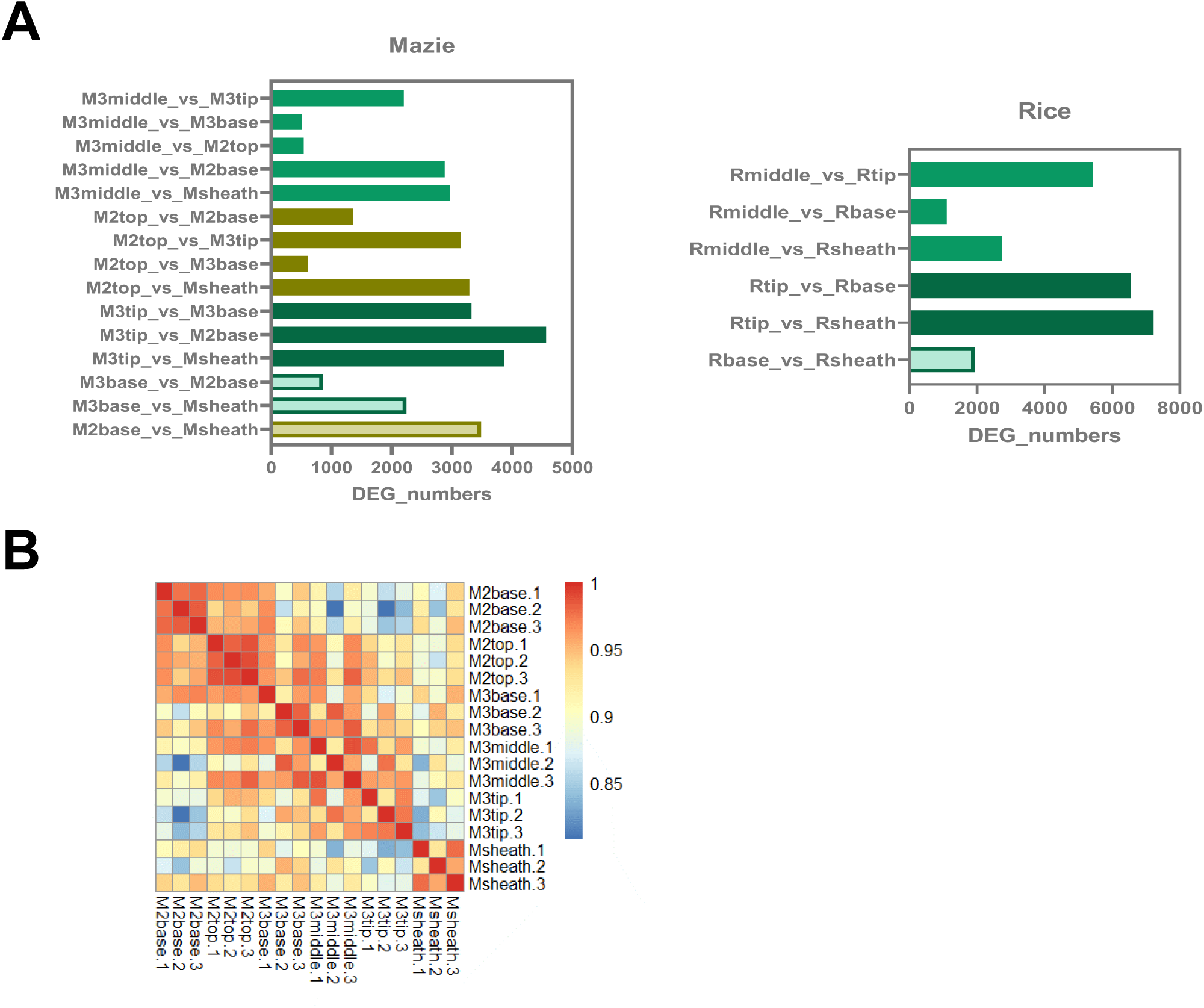
The DEGs of paired primordia samples from bulk RNAseq. **A.** The DEGs of paired samples across the 6 types of tissues in maize, and the 4 types of tissues in rice. Both up-regulated and down-regulated genes are included. **B.** Correlation matrix showing similarities among different maize samples.

**Supplemental Figure 4.**
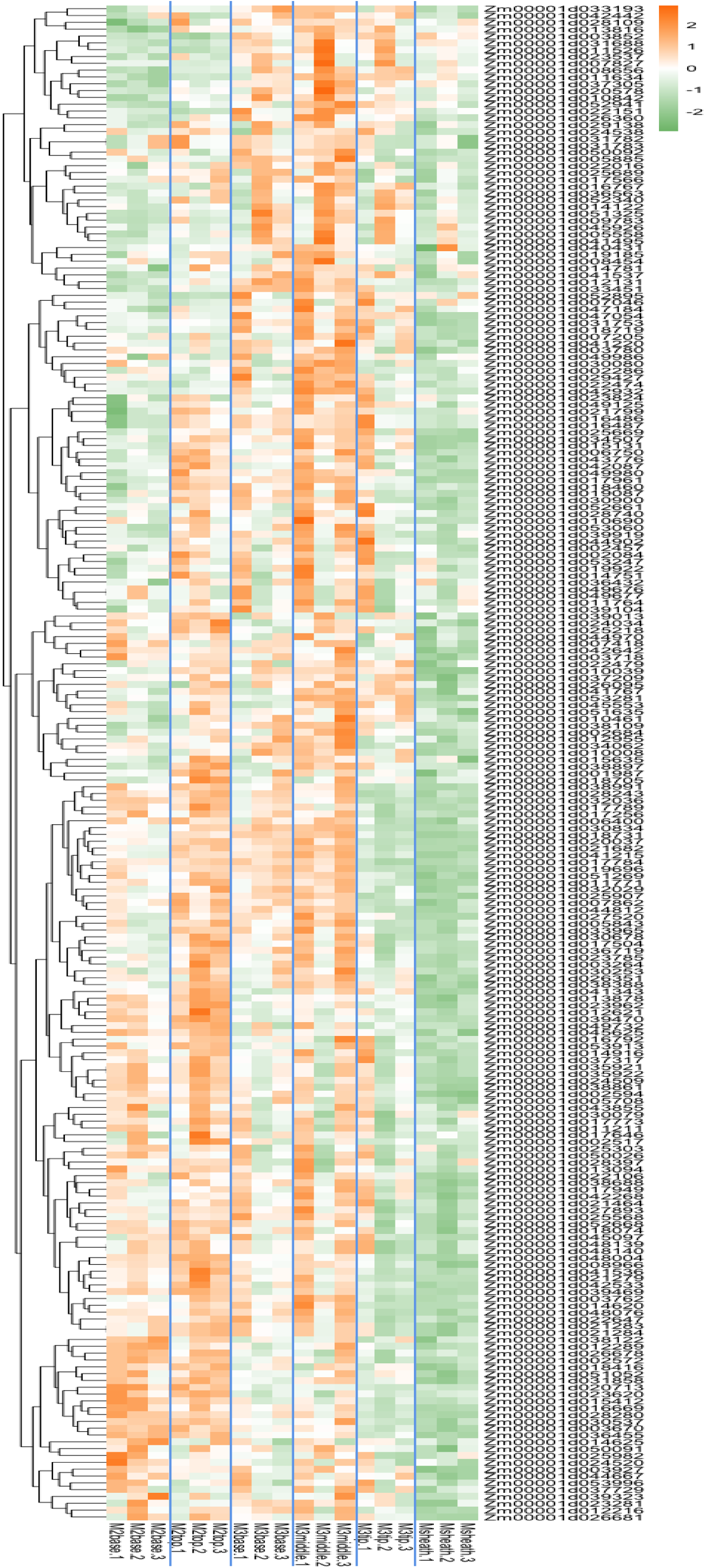
Heatmap showing the expression pattern of Kranz anatomy related genes obtained after multiple filtration steps. See Supplemental Dataset 9 for full size heatmap.

**Supplemental Figure 5.**
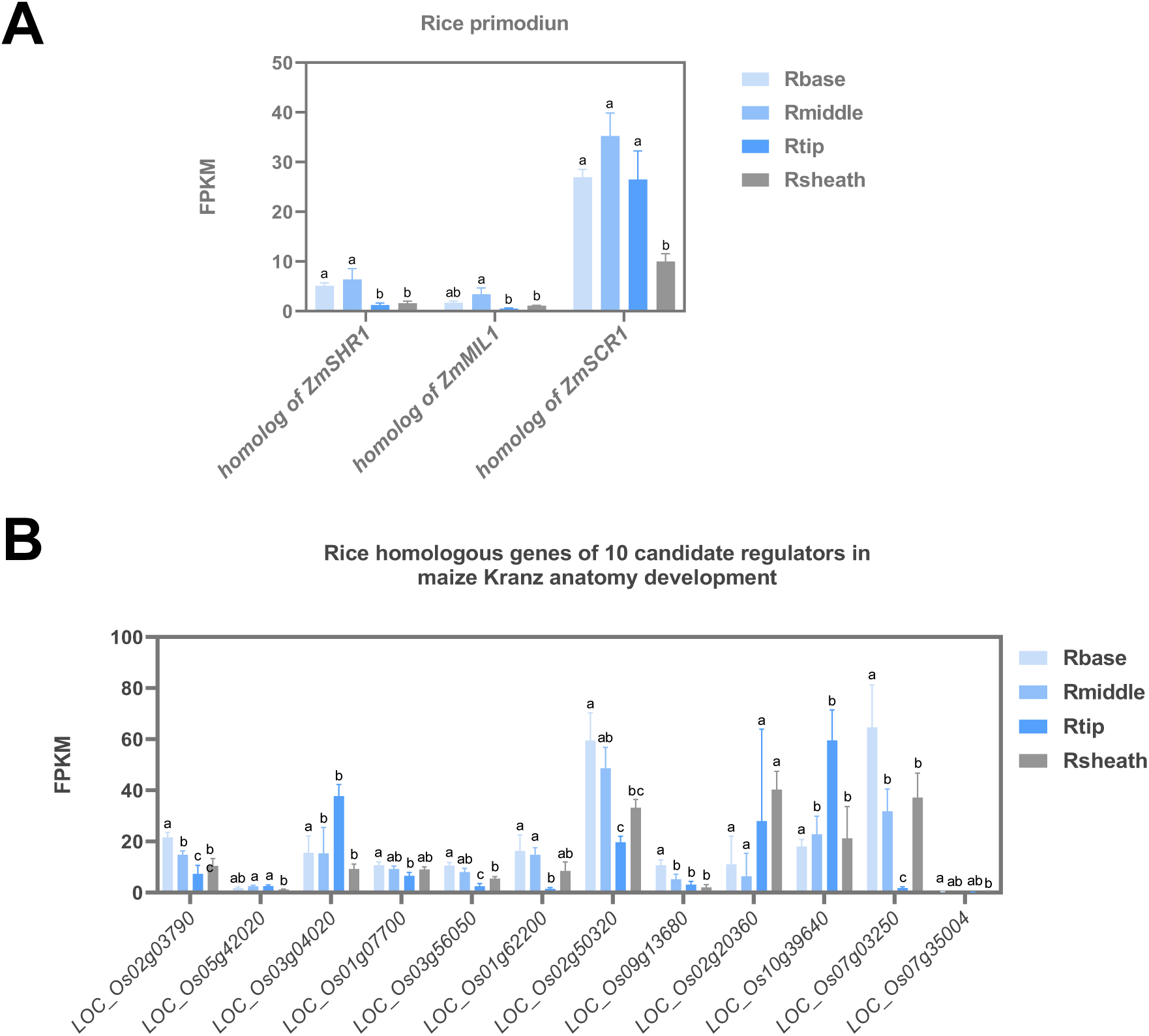
The expression pattern of rice homologous genes of interest in rice primodium. **A.** Bar graph illustrating the expression patterns of the homolog of *ZmSHR1*, *ZmSCR1*, and *ZmMIL1* across 4 rice primodium tissue types. **B.** Bar graph illustrating the expression profiles of the homologs of 10 putative Kranz anatomy regulators identified in this study. The gene list of B is given in Supplemental dataset 7. Error bars represent mean±SD (n=3). Statistical analysis was performed using one-way ANOVA with Tukey’s HSD test; *P* < 0.05, different letters on the bar graphs indicate statistically significant difference.

**Supplemental Figure 6.**
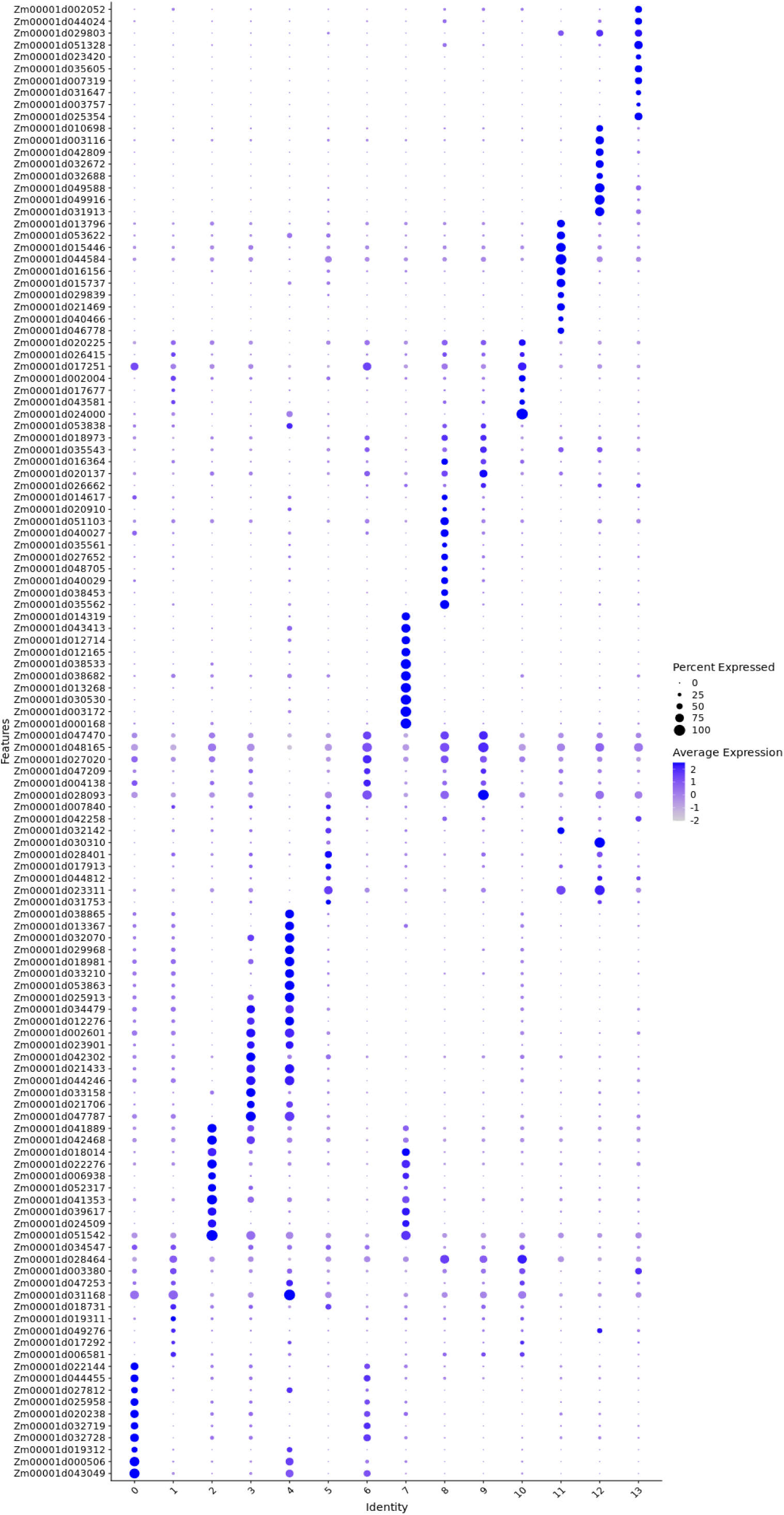
Dot map showing the top 10 marker genes from each of the different cell clusters.

**Supplemental Figure 7.**
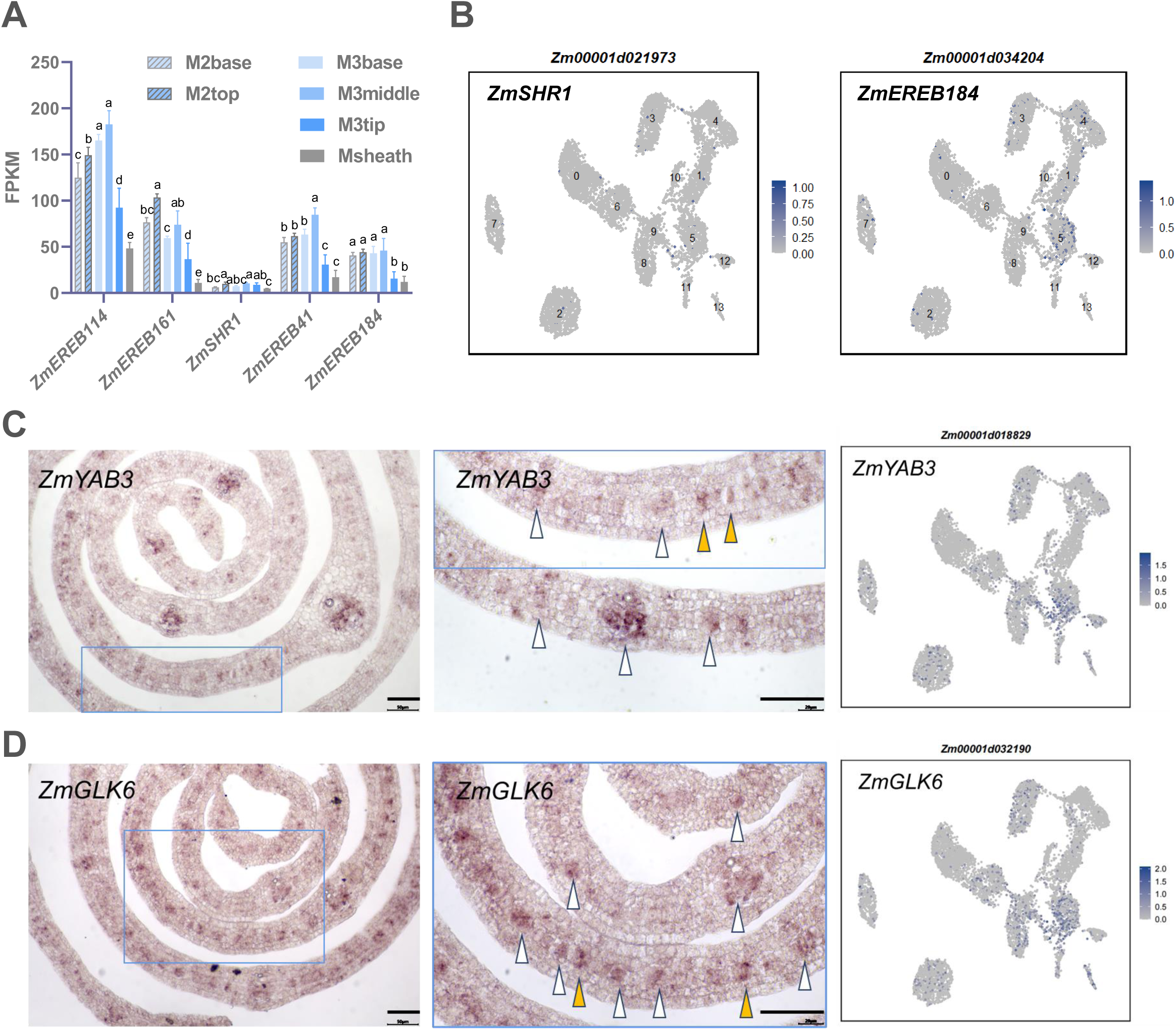
The expression patterns of *ZmSHR1*, *ZmEREB184*, *ZmYAB3* and *ZmGLK6*. **A and B.** The expression patterns of *ZmSHR1* and *ZmEREB184* across 6 maize tissue types and by UMAP plots. *EREB114*, *EREB161* and *EREB41* were presented in the column chart for comparison. Refer to Figures 6, 7 and 8 for *in situ* expression profiles. Error bars represent mean±SD (n=3). Statistical analysis in A was performed using one-way ANOVA with Tukey’s HSD test; *P* < 0.05, different letters on the bar graphs indicate statistically significant difference. **C and D.** Left column, *in situ* hybridization for the transcript localization of *ZmYAB3* and *ZmGLK6* on transverse sections of maize leaf primordia. Middle column, close-up images of the blue square framed regions from left column. Right column, expression pattern of selected genes by UMAP plots. Scale bar: 50 μm. White arrows indicate the differentiating procambium; orange arrows indicate the potential single procambial initial cell.

**Supplemental Figure 8.**
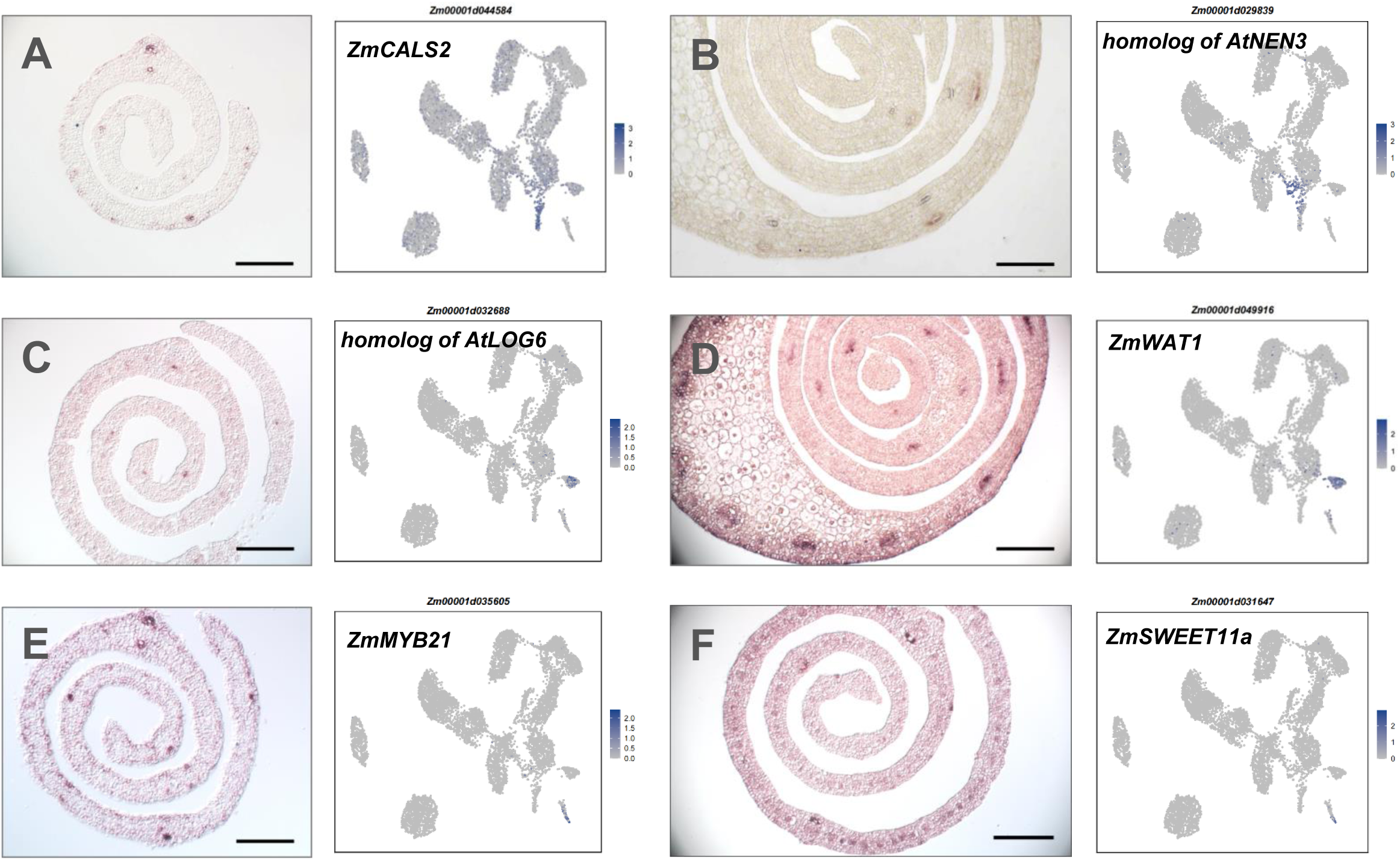
*In situ* expression patterns of representative marker genes from xylem and phloem related cell clusters. **A and B.** *In situ* hybridization and UMAP plots for the expression patterns of cluster 11 (phloem related)-enriched genes. **C and D.** *In situ* hybridization and UMAP plots for the expression patterns of cluster 12 (xylem related)-enriched genes. **E and F.** *In situ* hybridization and UMAP plots for the expression patterns of cluster 13-enriched genes. Names of the cluster-enriched genes are indicated above the figures. Scale bar: 100 μm.

**Supplemental Figure 9.**
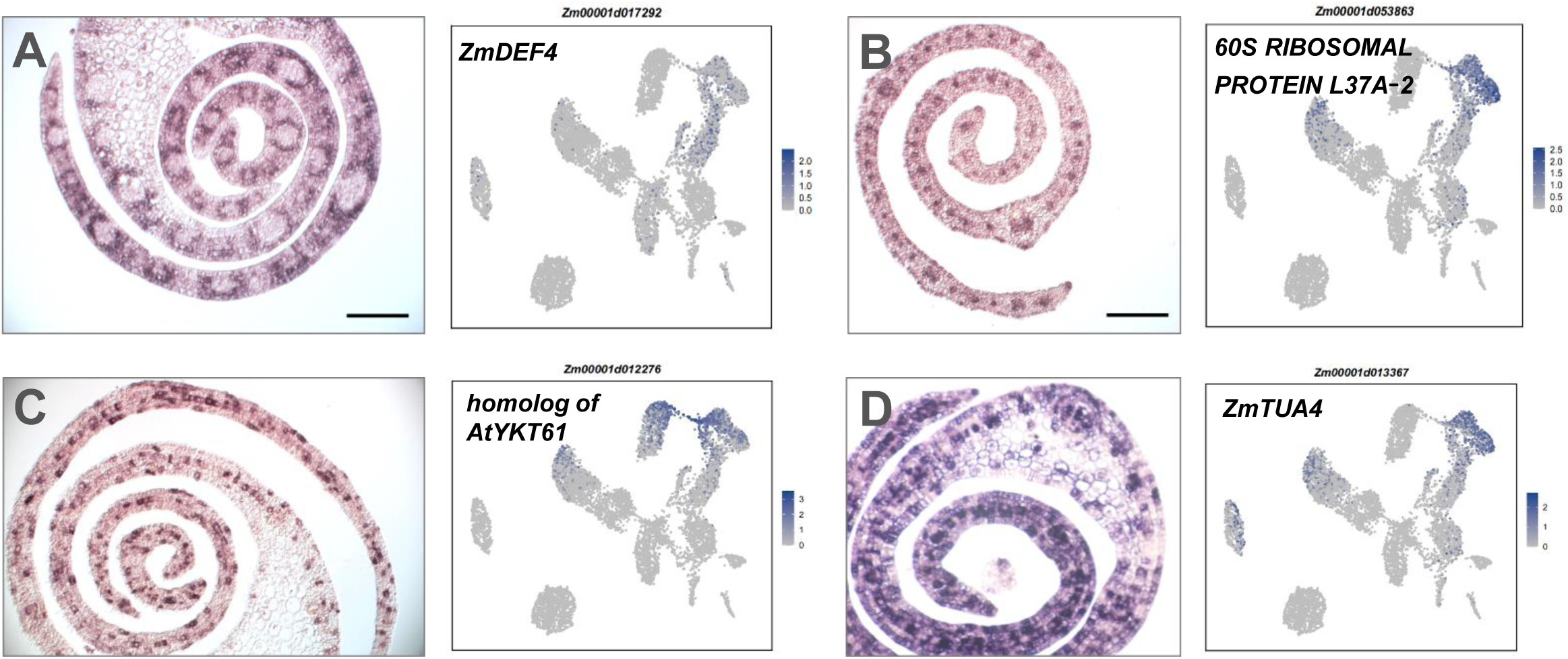
*In situ* expression patterns of representative marker genes related with clusters 1 and 4. A-D show *in situ* hybridization and UMAP plots for the expression patterns of clusters 0, 1, 3 and 4-enriched genes. Names of the cluster-enriched genes are indicated above the figures. Scale bar: 100 μm.

**Supplemental Figure 10.**
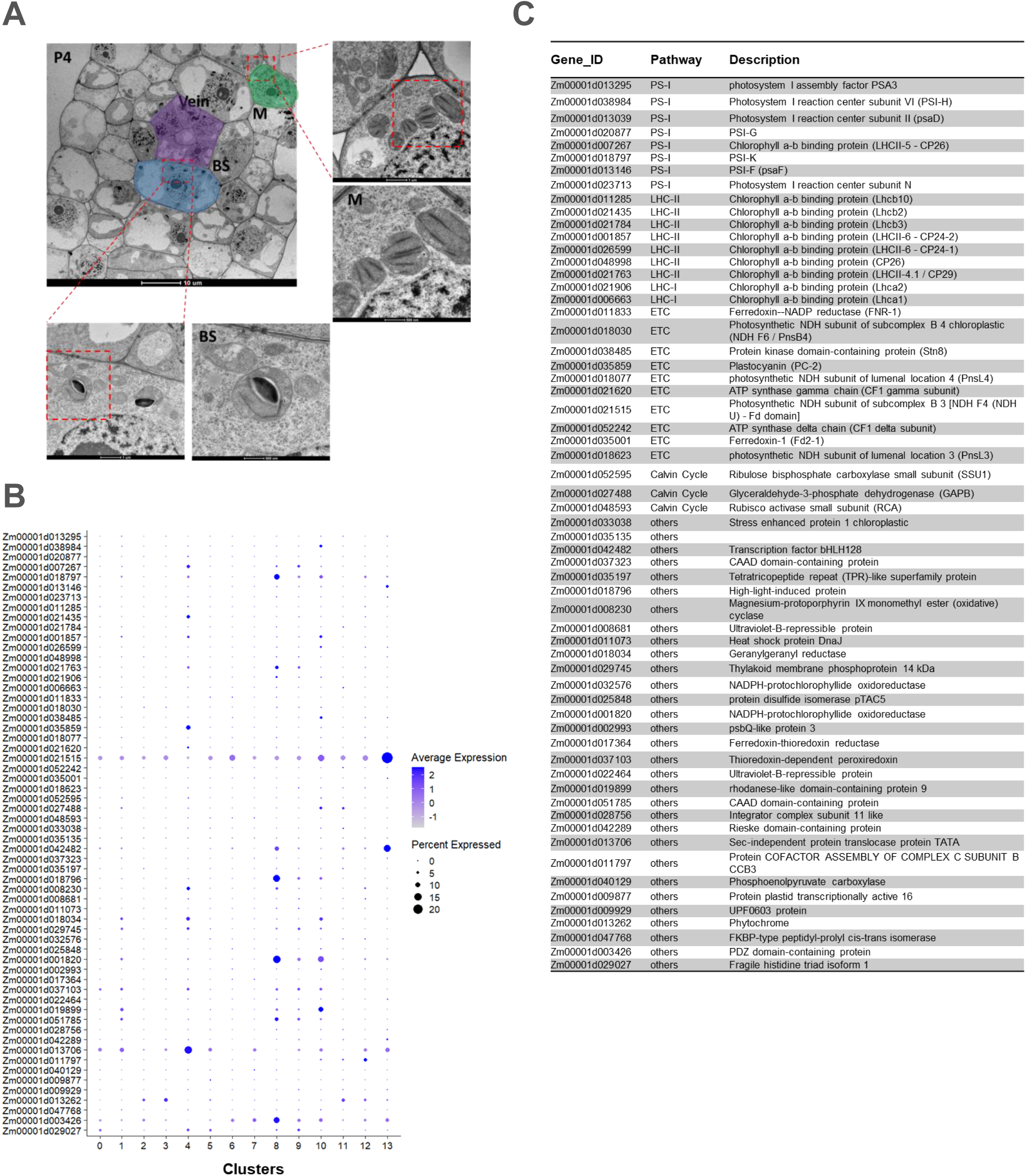
Transmission microscopy of maize leaf primordium and M3tip enrichment of photosynthesis-related genes. A. Transmission microscopy showing under-developed plastid in P4 leaf primordium. BS, bundle sheath cell; M, mesophyll cell; Scale bars: 10 μm, 1 μm, or 500 nm. **B.** Expression patterns of M3tip enriched photosynthetic genes. Dot colour, proportion of cluster cells expressing a given gene; Dot size, the average expression level. **C.** Annotation and description of genes listed in (B).

**Supplemental Figure 11.**
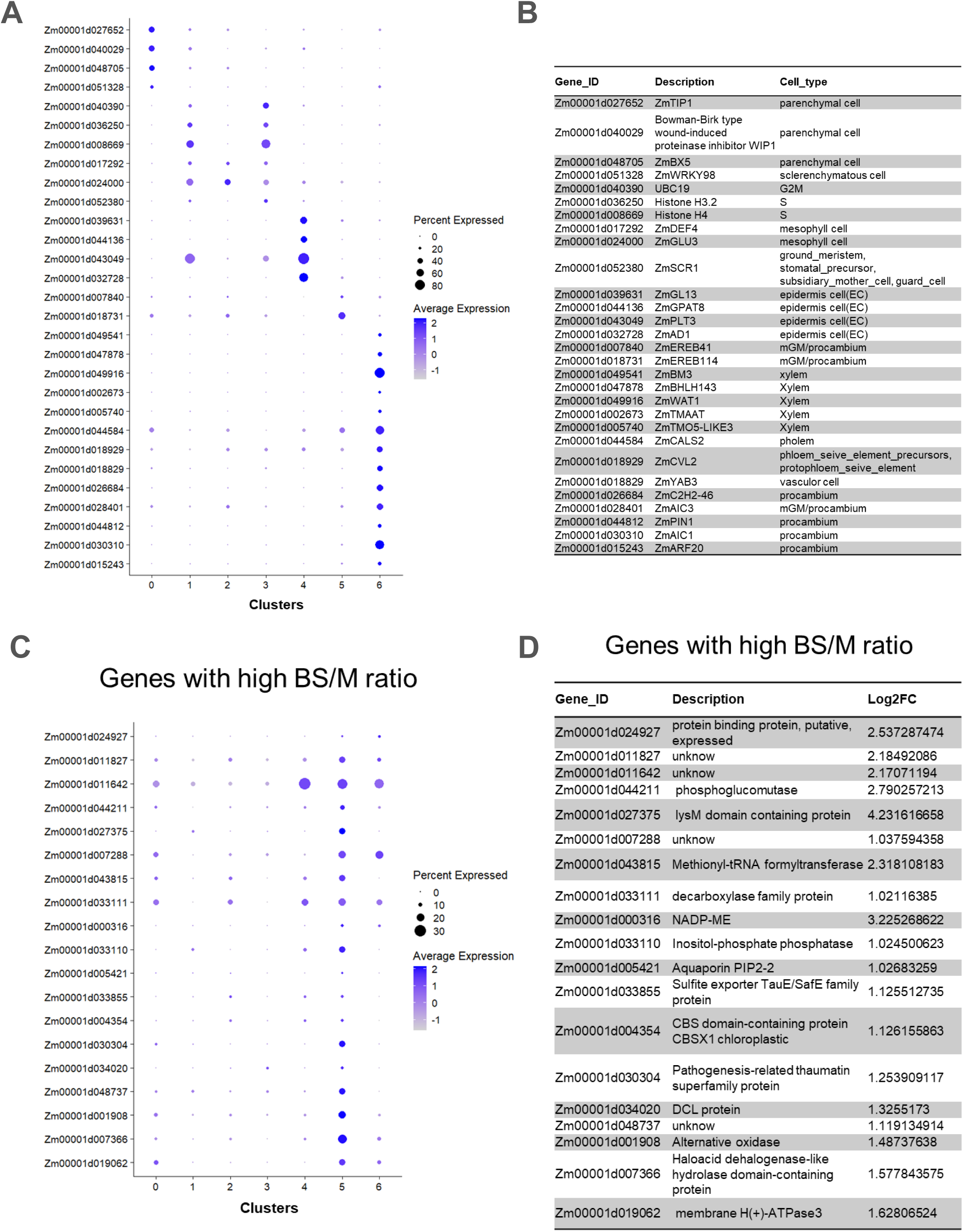
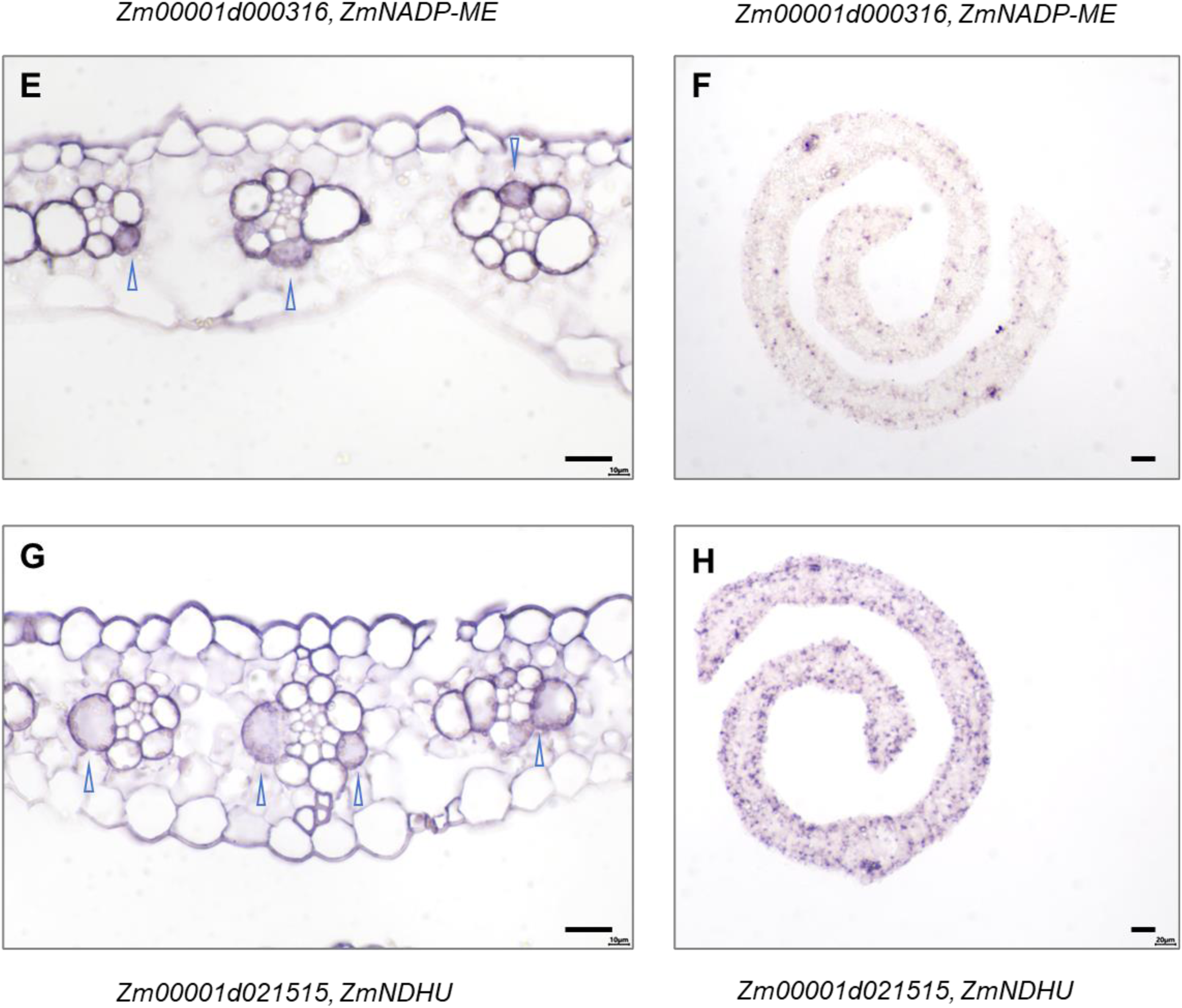
Cell heterogeneity and putative bundle sheath cell identity associated genes in the M3tip of maize leaf primordium. **A.** Expression patterns of representative cluster-specific marker genes. Dot colour, proportion of cluster cells expressing a given gene; Dot size, the average expression level. **B.** Annotation and description of genes listed in (A). **C.** Genes with high BS/M ratio of expression were potentially included in cluster 5 of **Figure 9A**. **D.** Annotation and description of genes listed in (C). **E and F.** *In situ* hybridization for the transcript localization of *ZmNADP-ME* on transverse sections of maize expended leaf (E) and leaf primordium (F). Scale bar: 20 μm. Blue arrows indicate bundle sheath cells with transcript enrichment. **G and H.** *In situ* hybridization for the transcript localization of *ZmNDHU* on transverse sections of maize expended leaf (G) and leaf primordium (H). Scale bar: 20 μm. Blue arrows indicate bundle sheath cells with transcript enrichment.

**Supplemental Figure 12.**
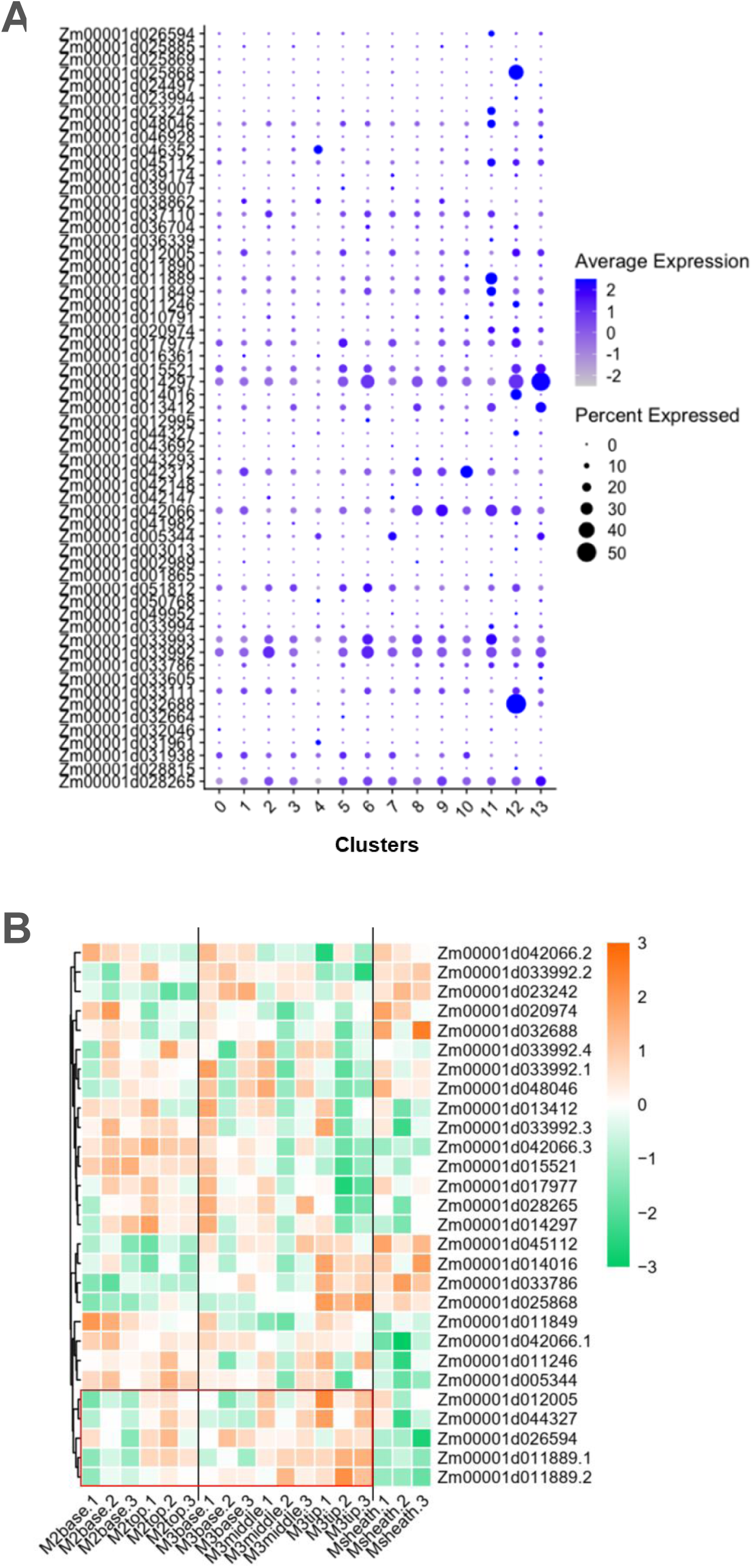
Expression of genes related to CK signaling pathway in maize leaf primordium. (A) and (B) show the distribution of gene expression in the total primordium cell clusters and tissue subsections respectively.

**Supplemental Figure 13.**
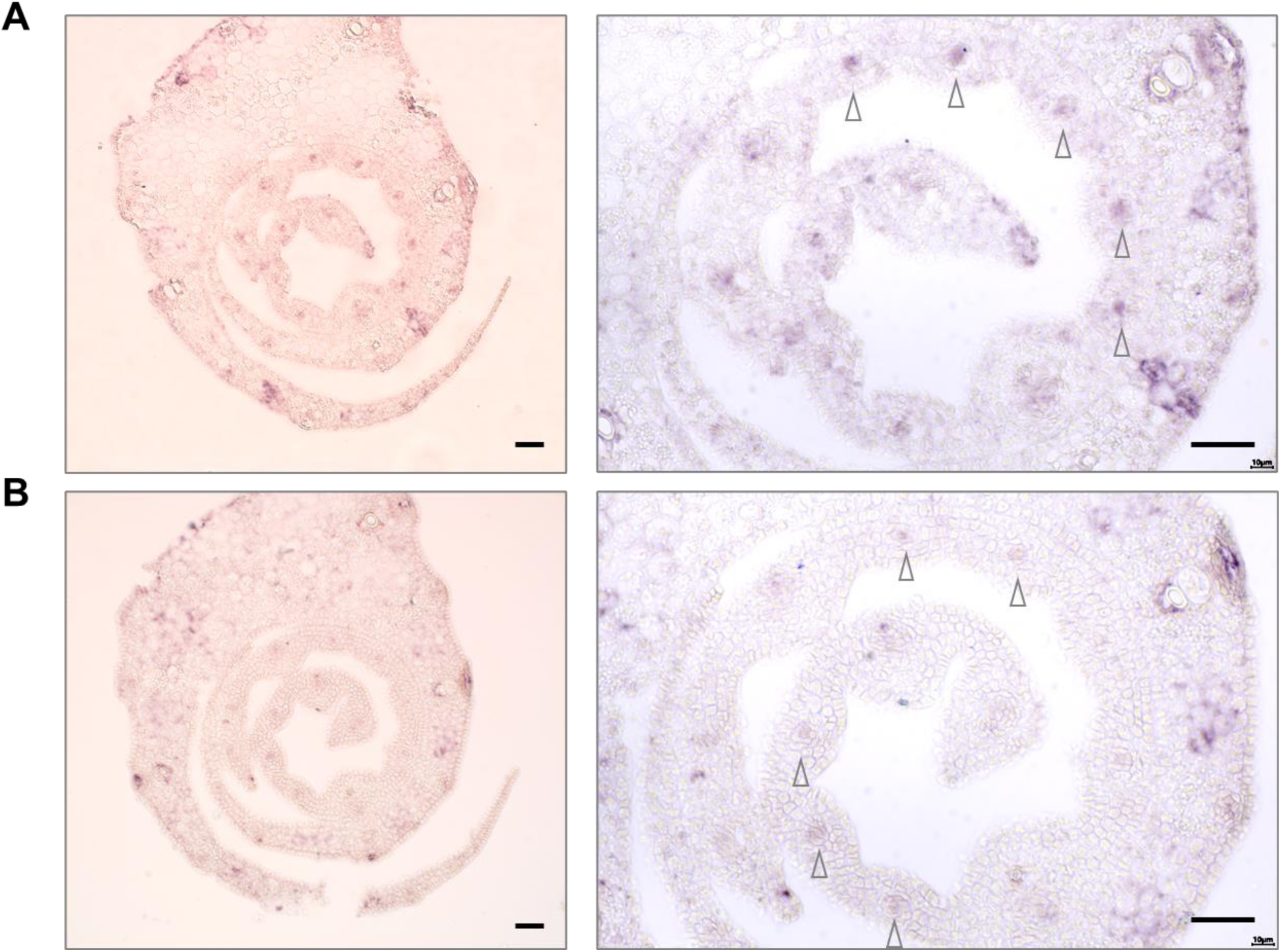
*In situ* expression of *OsEREB161* transcripts in rice leaf primordium. (A) and (B) are images from 2 different sections of 5mm rice primordium. Scale bar: 30 μm. Arrows indicate the widely spaced vascular procambia where the *OsEREB161* expression is restricted in.

**Supplemental Figure 14.**
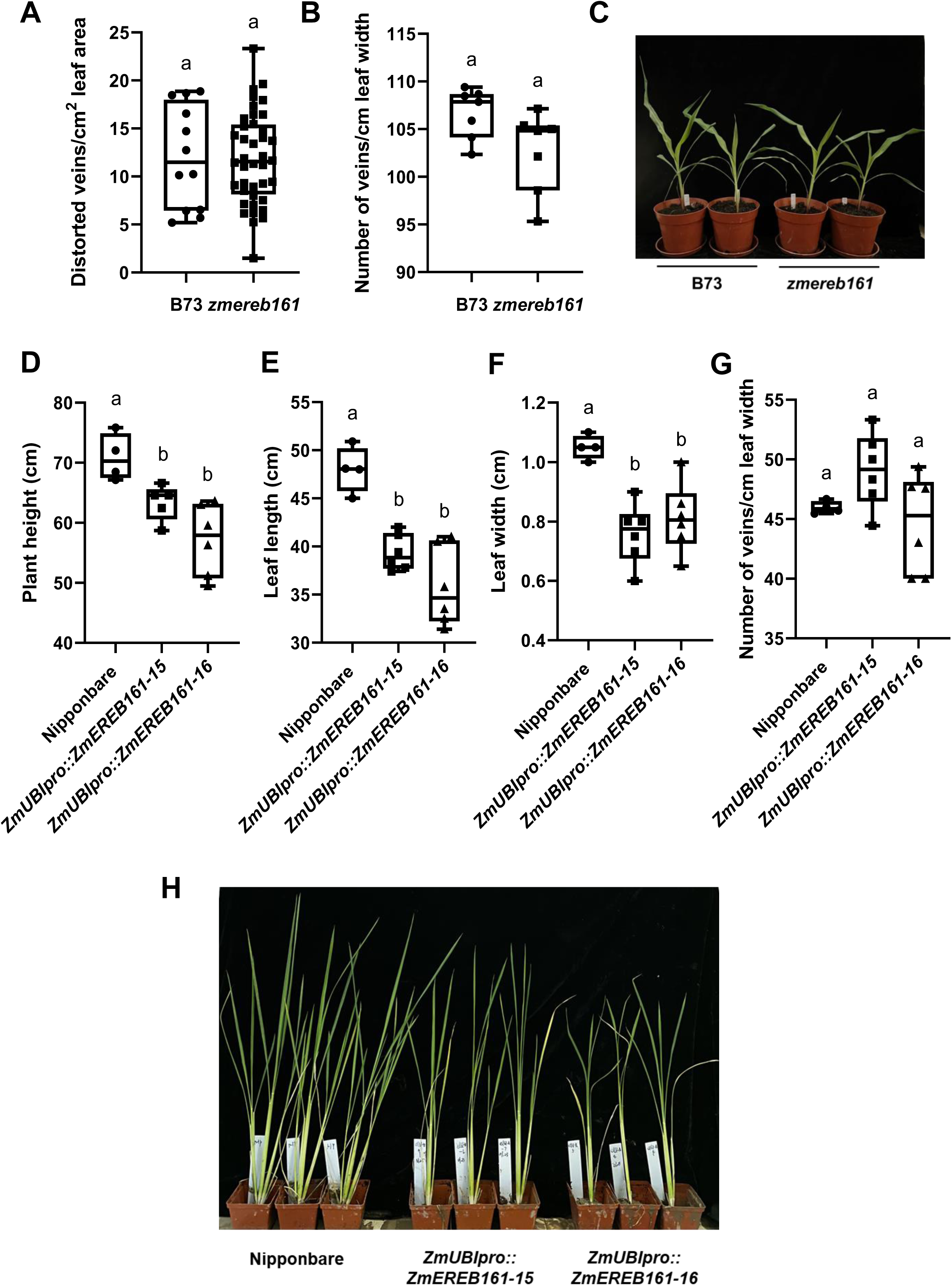
Characterization of maize *zmereb161* mutant and *ZmUBIpro::ZmEREB161* transgenic rice plants. **A and B.** Quantification of leaf vascular traits of maize wild-type (B73) and *ereb161* mutant. Boxplots show emergence rate of distorted veins (A) and vein density (B) in the middle sections of 2^nd^ fully expended leaves from the top of 4-weeks-old plants. **C.** The phenotype of maize wild-type and *zmereb161* mutant plants. **D-G.** Quantification of plant growth and leaf traits of rice wild-type (Nipponbare) and *ZmUBIpro::ZmEREB161* transgenic plants. Boxplots show plant height (D), leaf length (E), leaf width (F), and vein density (G). (F) and (G) were measured in the middle sections of the 2^nd^ fully expended leaves from the top of 6-weeks-old T1 plants. **H.** The phenotype of rice wild-type and *ZmUBIpro::ZmEREB161* plants. The box, black horizontal line, and whiskers indicate data within interquartile range (IQR, 25th-75th percentiles), the median, lowest and highest value within 1.5 times the IQR, respectively; the data represent means ± SD and *P* values are calculated using an unpaired *t* test (A and B, n >6) or 1-way ANOVA with Tukey’s HSD test (C-F, n=4 for wild-type, and n=6 for over-expression lines) ; different letters above the bars indicate significant differences (*P* < 0.05).

**Supplemental Figure 15.**
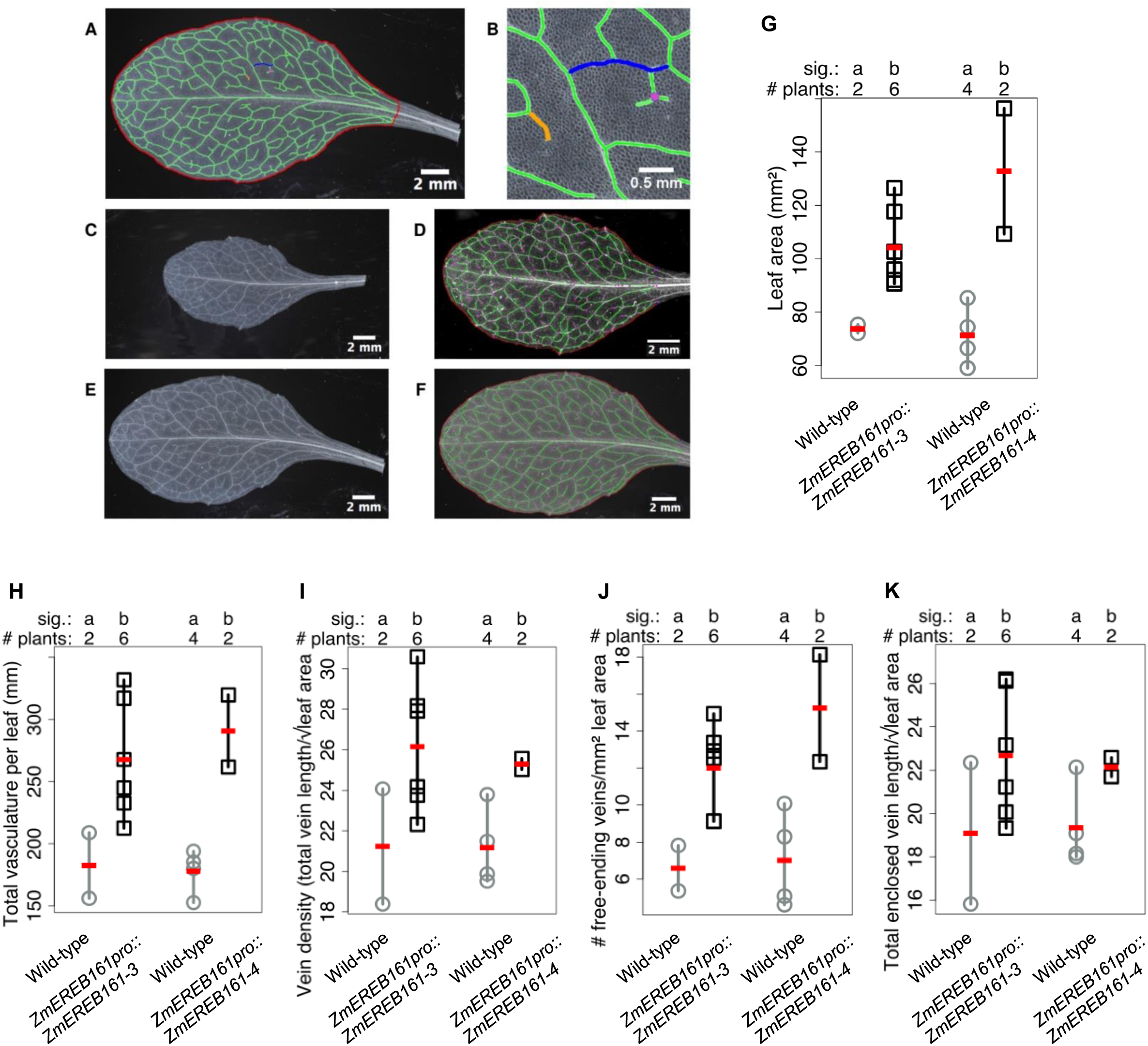
Z*m*EREB161pro*::ZmEREB161* transgenic arabidopsis lines have more vasculature and a higher vein density compared to wild-type. **A and B.** Dark field photographs of cleared arabidopsis leaf 5 overlaid with the leaf traits measured: leaf area (red), total vasculature (green), enclosed veins (blue), free-ending veinlets (orange) and vein branching points (purple)-in a whole leaf (A) and a magnified image (B). **C-F.** Darkfield photograph of a segregating *ZmEREB161pro::ZmEREB161* (wild-type) T2 sibling (C) overlaid with LIMANI output (D), and darkfield photograph of a *ZmEREB161pro::ZmEREB161* T2 sibling (E) overlaid with LIMANI output (F). **G-K.** Strip charts showing whole leaf - total leaf 5 area (mm^2^) (G), total vasculature (mm) (H), vein density (mm/mm^2^) (total vein length/leaf area) (I), free-ending veinlet number per mm^2^ leaf area (J) and enclosed vein density (total enclosed vein length/leaf area) (K). “Sig.” - different letters indicate significant differences in group means (*P*<0.05). # plants - number of individual plants used for analysis. Square/circle - individual data point, vertical line - range of data, red dash - mean.

**Supplemental Figure 16.**
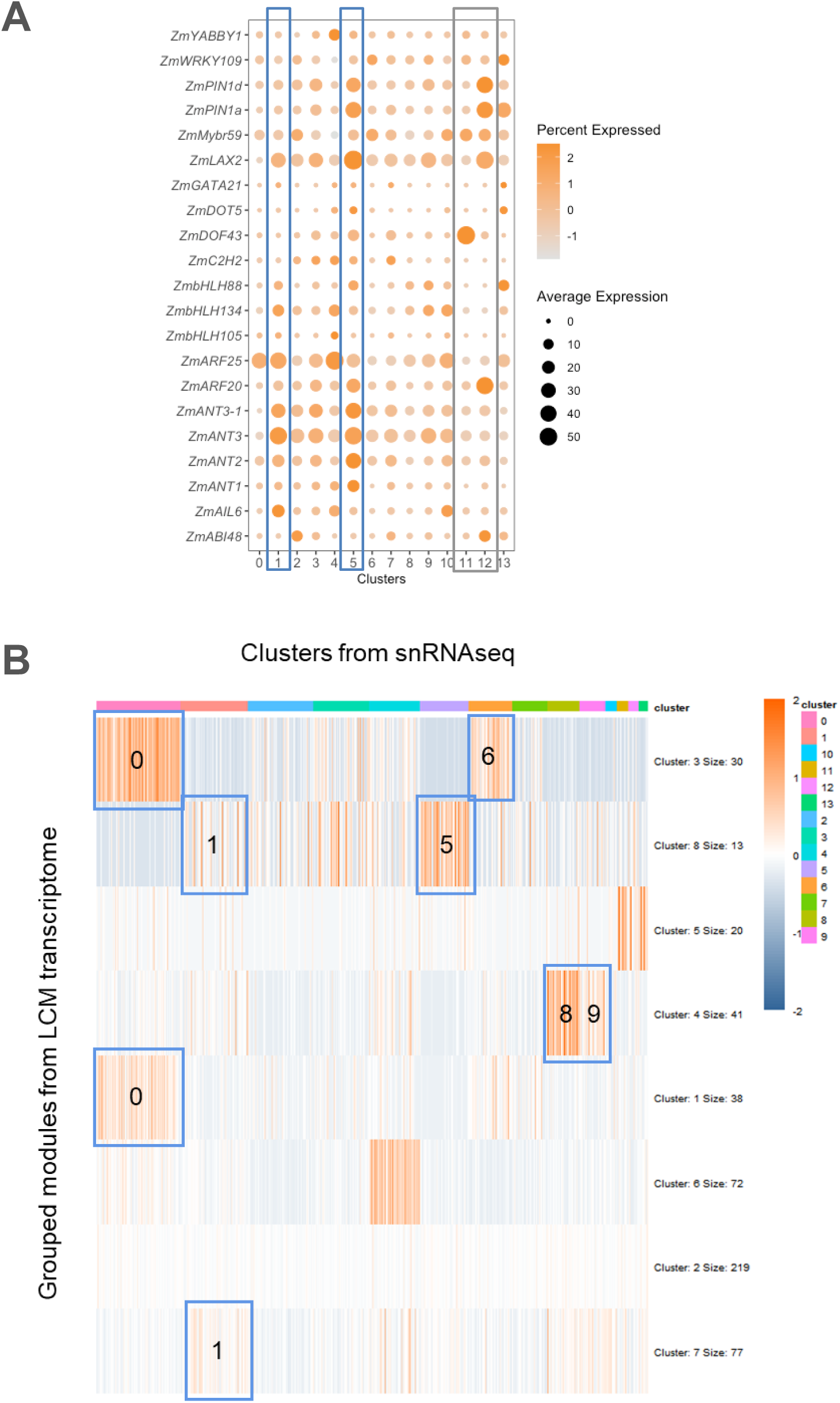
Comparison between current dataset and Liu et al. (2022). **A.** Cell cluster (of the 14 clusters from **Figure 4A**) distribution of genes that are highly expressed in median ground meristem (mGM) and three-contiguous cell (3C) from the LCM-transcriptomes (Fig. 7A of Liu et al., 2022). **B.** A comprehensive comparison of current snRNA-seq data with the LCM-transcriptomic data of Liu et al. (2022). Blue frames with numbers indicate genes in different clusters of current snRNA-seq data overlapping with grouped modules from the LCM-transcriptome data.

**Supplemental Figure 17.**
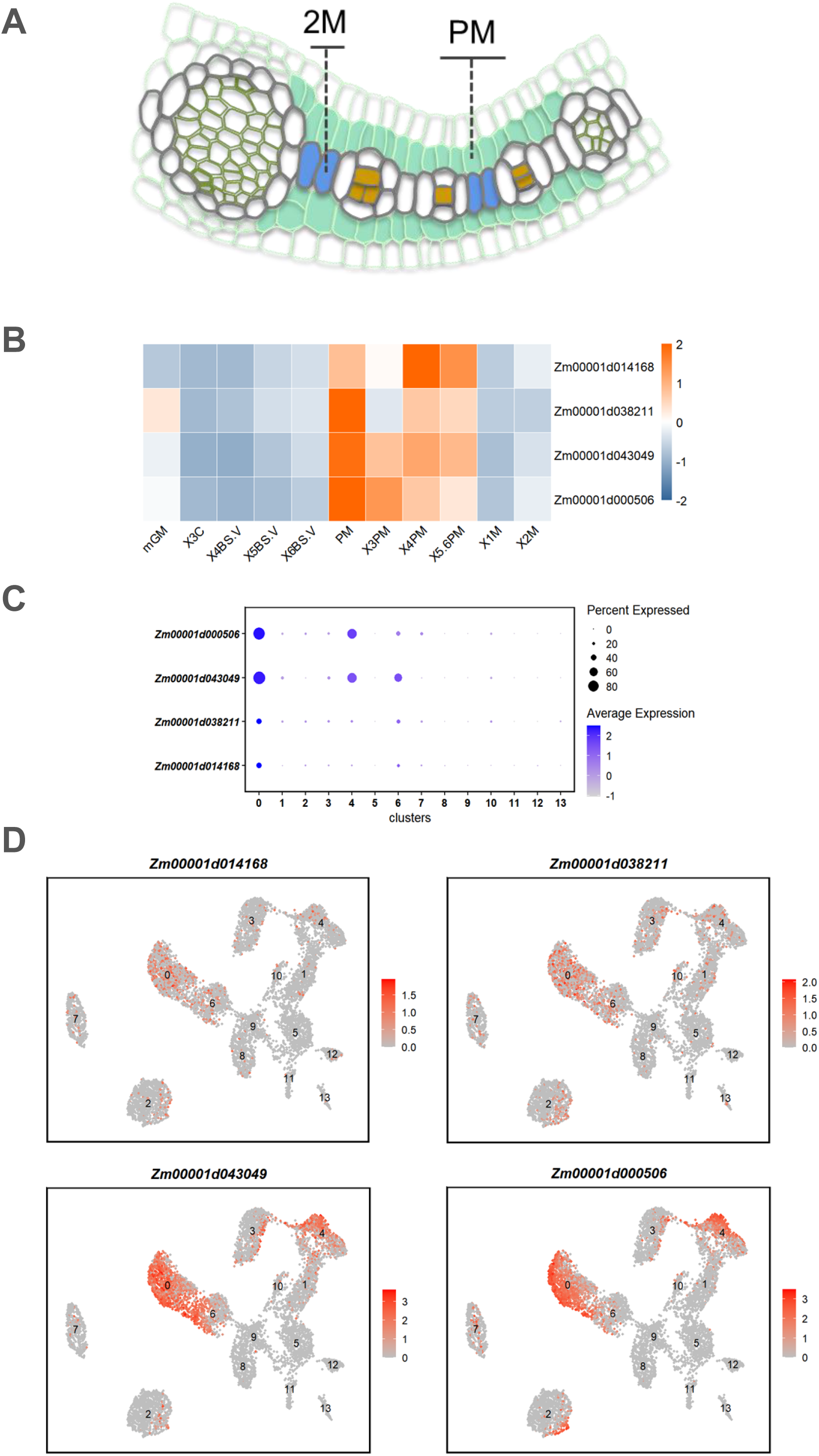
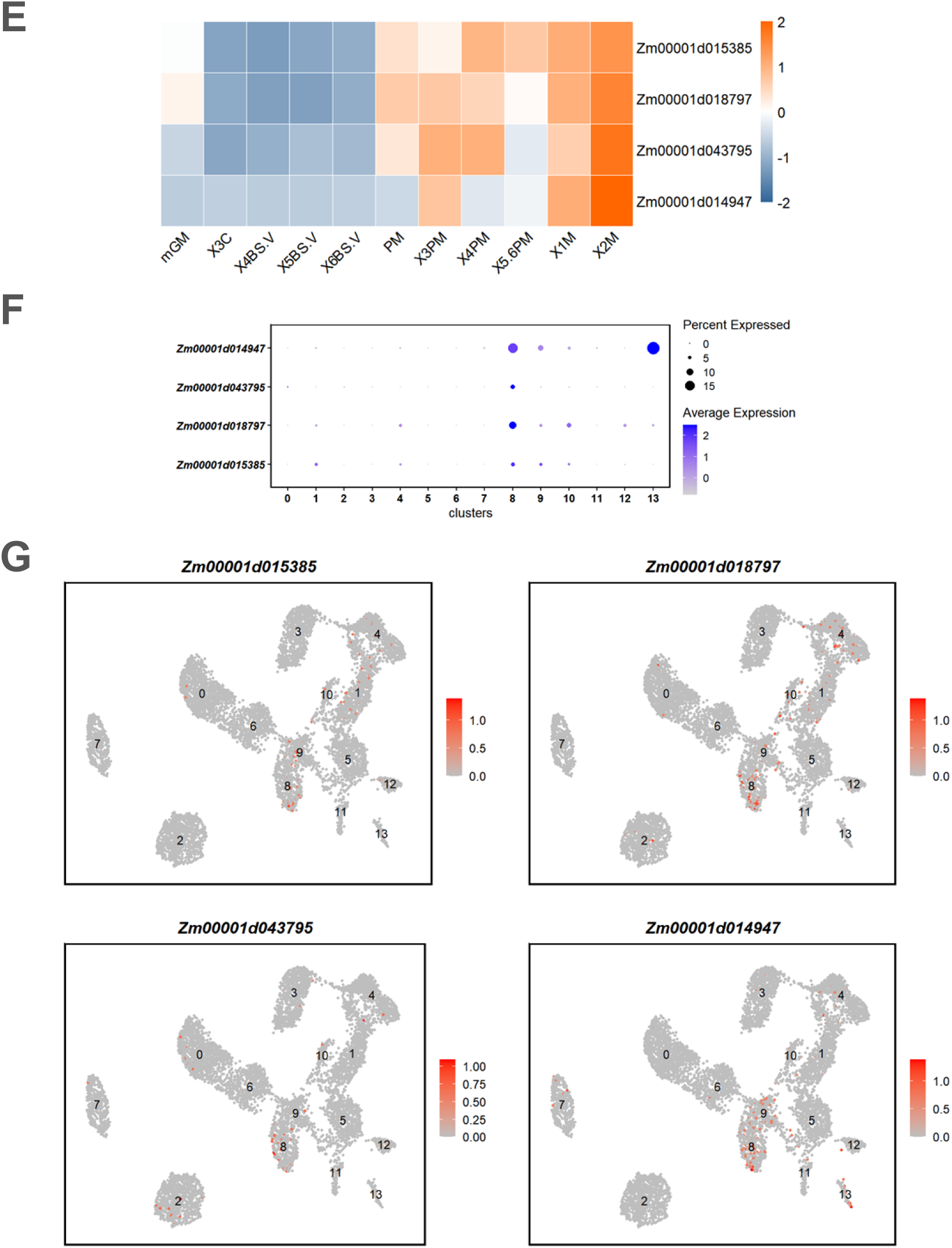
“PM” and “2M” samples from the LCM data assisted further annotation of mesophyll cell clusters for the snRNAseq data. **A.** Schematics of the anatomy and cell types representing the upper middle section of P4 leaf primordium. “2M” and “PM” cells described in Liu et al. (2022) were filled in bule and light green colors respectively. **B-D.** Heat maps showing the selected “PM” cell enriched genes (B); Dot plots (C) and UMAP plots (D) showing the “cluster 0” enrichment of the selected genes. **E-G.** Heat maps showing the selected “2M” cell enriched genes (E); Dot plots (F) and UMAP plots (G) showing the “cluster 8” enrichment of the selected genes. Dot colour, proportion of cluster cells expressing a given gene; Dot size, the average expression level.

## References

Aubry S, Kelly S, Kümpers BM, Smith-Unna RD, Hibberd JM. 2014. Deep evolutionary comparison of gene expression identifies parallel recruitment of trans-factors in two independent origins of C4 photosynthesis. PLoS Genet 10: e1004365.

Arrivault S, Alexandre Moraes T, Obata T, Medeiros DB, Fernie AR, Boulouis A, Ludwig M, Lunn JE, Borghi GL, Schlereth A, Guenther M, Stitt M. 2019. Metabolite profiles reveal interspecific variation in operation of the Calvin-Benson cycle in both C4 and C3 plants. J Exp Bot. 70(6):1843–1858.

Bassham JA. 2003. Mapping the carbon reduction cycle, a personal retrospective. Photosynthesis Reseach 76: 35–52.

Bezrutczyk M, Zöllner NR, Kruse CPS, Hartwig T, Lautwein T, Köhrer K, Frommer WB, Kim JY. 2021. Evidence for phloem loading via the abaxial bundle sheath cells in maize leaves. Plant Cell 33(3): 531–547.

Bräutigam A, Kajala K, Wullenweber J, Sommer M, Gagneul D, Weber KL, Carr KM, Gowik U, Mass J, Lercher MJ, Westhoff P, Hibberd JM, Weber AP. 2011. An mRNA blueprint for C4 photosynthesis derived from comparative transcriptomics of closely related C3 and C4 species. Plant Physiol. 155(1):142–56.

Bräutigam A, Schliesky S, Külahoglu C, Osborne CP, Weber AP. 2014. Towards an integrative model of C4 photosynthetic subtypes: insights from comparative transcriptome analysis of NAD-ME, NADP-ME, and PEP-CK C4 species. J Exp Bot. 65(13):3579–93.

Billakurthi K, Wrobel TJ, Bräutigam A, Weber AP, Westhoff P, Gowik U. 2018. Transcriptome dynamics in developing leaves from C3 and C4 Flaveria species reveal determinants of Kranz anatomy. bioRxiv doi: 10.1101/473181

Bosch M, Mayer CD, Cookson A, Donnison IS. 2011. Identification of genes involved in cell wall biogenesis in grasses by differential gene expression profiling of elongating and non-elongating maize internodes. J Exp Bot 62(10): 3545–3561.

Brown WV. 1975. Variations in anatomy, associations, and origins of Kranz tissue. American Journal of Botany 62: 395–402.

Burgess SJ, Granero-Moya I, Grangé-Guermente MJ, Boursnell C, Terry MJ, Hibberd JM. 2016. Ancestral light and chloroplast regulation form the foundations for C4 gene expression. Nature Plants 2(11): 16161.

Bakken TE, Hodge RD, Miller JA, Yao Z, Nguyen TN, Aevermann B, Barkan E, Bertagnolli D, Casper T, Dee N, Garren E, Goldy J, Graybuck LT, Kroll M, Lasken RS, Lathia K, Parry S, Rimorin C, Scheuermann RH, Schork NJ, Shehata SI, Tieu M, Phillips JW, Bernard A, Smith KA, Zeng H, Lein ES, Tasic B. 2018. Single-nucleus and single-cell transcriptomes compared in matched cortical cell types. PLoS One. 13(12):e0209648.

Chang YM, Liu WY, Shih ACC, Shen MN, Lu CH, Lu MYJ, Yang HW, Wang TY, Chen SCC, Chen SM, Li WH, Ku MSB. 2012. Characterizing regulatory and functional differentiation between maize mesophyll and bundle sheath cells by transcriptomic analysis. Plant Physiol 160: 165–177.

Christin PA, Boxall SF, Gregory R, Edwards EJ, Hartwell J, Osborne CP. 2013. Parallel recruitment of multiple genes into c4 photosynthesis. Genome Biol Evol. 5(11):2174–87.

Covshoff S, Szecowka M, Hughes TE, Smith-Unna R, Kelly S, Bailey KJ, Sage TL, Pachebat JA, Leegood R, Hibberd JM. 2016. C4 Photosynthesis in the Rice Paddy: Insights from the Noxious Weed Echinochloa glabrescens. Plant Physiol. 170(1):57–73.

Conde D, Triozzi PM, Balmant KM, Doty AL, Miranda M, Boullosa A, Schmidt HW, Pereira WJ, Dervinis C, Kirst M. 2021. A robust method of nuclei isolation for single-cell RNA sequencing of solid tissues from the plant genus Populus. PLoS One 16(5): e0251149.

Conklin PA, Strable J, Li S, Scanlon MJ. 2019. On the mechanisms of development in monocot and eudicot leaves. New Phytol 221(2): 706–724.

Ding Z, Weissmann S, Wang M, Du B, Huang L, Wang L, Tu X, Zhong S, Myers C, Brutnell TP, Sun Q, Li P. 2015. Identification of Photosynthesis-Associated C4 Candidate Genes through Comparative Leaf Gradient Transcriptome in Multiple Lineages of C3 and C4 Species. PLoS One. 10(10):e0140629.

Dhondt S, Van Haerenborgh D, Van Cauwenbergh C, Merks RMH, Philips W, Beemster GTS & Inze D. 2012. Quantitative analysis of venation patterns of Arabidopsis leaves by supervised image analysis. The Plant Journal, 69, 553–563.

Dong WT, Chang TG, Dai HL, Yang WB, Su Y, Chao DY, Zhu XG, Wang P, Yu N, Wang ET. 2023. Creating a C4-like vein pattern in rice by manipulating SHORT ROOT and auxin levels. Science Bulletin 68(24):3133–3136. doi: 10.1016/j.scib.2023.10.005.

Edwards EJ, Still CJ. 2008. Climate, phylogeny and the ecological distribution of C4 grasses. Ecol Lett 11(3): 266–276.

Fouracre JP, Ando S, Langdale JA. 2014. Cracking the Kranz enigma with systems biology. Journal of Experimental Botany 65(13): 3327–3339.

Furbank RT. 2017. Walking the C4 pathway: past, present, and future. J Exp Bot 68(2): 4057–4066.

Furumoto T, Yamaguchi T, Ohshima-Ichie Y, Nakamura M, Tsuchida-Iwata Y, Shimamura M, Ohnishi J, Hata S, Gowik U, Westhoff P, Bräutigam A, Weber AP, Izui K. 2011. A plastidial sodium-dependent pyruvate transporter. Nature. 476(7361):472–5.

Frey M, Chomet P, Glawischnig E, Stettner C, Grün S, Winklmair A, Eisenreich W, Bacher A, Meeley RB, Briggs SP, Simcox K, Gierl A. 1997. Analysis of a chemical plant defense mechanism in grasses. Science 277(5326): 696–699.

Farmer A, Thibivilliers S, Ryu KH, Schiefelbein J, Libault M. 2021. Single-nucleus RNA and ATAC sequencing reveals the impact of chromatin accessibility on gene expression in Arabidopsis roots at the single-cell level. Mol Plant. 14(3):372–383.

Ghannoum, O., Evans, J.R., von Caemmerer, S. 2010. Chapter 8 Nitrogen and Water Use Efficiency of C4 Plants. In: Raghavendra, A., Sage, R. (eds) C4 Photosynthesis and Related CO2 Concentrating Mechanisms. Advances in Photosynthesis and Respiration, vol 32. Springer, Dordrecht.

Gowik U, Bräutigam A, Weber KL, Weber AP, Westhoff P. 2011. Evolution of C4 photosynthesis in the genus Flaveria: how many and which genes does it take to make C4? Plant Cell. 23(6):2087–105.

Germain PL, Lun A, Garcia Meixide C, Macnair W, Robinson MD. 2021. Doublet identification in single-cell sequencing data using scDblFinder. F1000Res 10:979.

Hao Y, Hao S, Andersen-Nissen E, et al. 2021. Integrated analysis of multimodal single-cell data. Cell 184(13): 3573–3587.

Hatch MD, Slack CR. 1998. C4 photosynthesis, discovery, resolution, recognition and significance. In: Kung S, Yang S, eds. Discoveries in plant biology, Vol 1. Singapore: World Scientific Publishing, 175–196.

Hendron RW, Kelly S. 2020. Subdivision of Light Signaling Networks Contributes to Partitioning of C4 Photosynthesis. Plant Physiol. 182(3):1297–1309.

Huang P, Brutnell TP. 2016. A synthesis of transcriptomic surveys to dissect the genetic basis of C4 photosynthesis. Current Opinion in Plant Biology 31: 91–99.

Hughes TE, Sedelnikova O, Thomas M, Langdale JA. 2023. Mutations in NAKED-ENDOSPERM IDD genes reveal functional interactions with SCARECROW during leaf patterning in C4 grasses. PLoS Genet 19(4): e1010715.

Han M, Park Y, Kim I, Kim EH, Yu TK, Rhee S, Suh JY. 2014. Structural basis for the auxin-induced transcriptional regulation by Aux/IAA17. Proc Natl Acad Sci U S A. 111(52):18613–8.

Jin J, Tian F, Yang DC, et al. 2017. PlantTFDB 4.0: toward a central hub for transcription factors and regulatory interactions in plants. Nucleic Acids Res 45(D1): D1040–D1045. doi:10.1093/nar/gkw982

John CR, Smith-Unna RD, Woodfield H, Covshoff S, Hibberd JM. 2014. Evolutionary convergence of cell-specific gene expression in independent lineages of C4 grasses. Plant Physiol 165: 6275.

Kitomi Y, Ito H, Hobo T, Aya K, Kitano H, Inukai Y. 2011. The auxin responsive AP2/ERF transcription factor CROWN ROOTLESS5 is involved in crown root initiation in rice through the induction of OsRR1, a type-A response regulator of cytokinin signaling. Plant J. 67(3):472–84.

Külahoglu C, Denton AK, Sommer M, Maß J, Schliesky S, Wrobel TJ, Berckmans B, Gongora-Castillo E, Buell CR, Simon R, De Veylder L, Bräutigam A, Weber AP. 2014. Comparative transcriptome atlases reveal altered gene expression modules between two Cleomaceae C3 and C4 plant species. Plant Cell. 26(8):3243–60.

Kurihara D, Mizuta Y, Sato Y. & Higashiyama T. 2015. ClearSee: a rapid optical clearing reagent for whole-plant fuorescence imaging. Development 142, 4168–4179.

Langdale JA. 1994. In situ Hybridization. In: Freeling, M., Walbot, V. (eds) The Maize Handbook. Springer Lab Manuals. Springer, New York, NY. 10.1007/978-1-4612-2694-9_18.

Langdale JA, Zelitch I, Miller E & Nelson T. 1988. Cell position and light influence C4 versus C3 patterns of photosynthetic gene expression in maize. EMBO J. 7: 3643–3651.

Li P, Ponnala L, Gandotra N, Wang L, Si Y, Tausta SL, Kebrom TH, Provart N, Patel R, Myers CR, Reidel EJ, Turgeon R, Liu P, Sun Q, Nelson T, Brutnell TP. 2010. The developmental dynamics of the maize leaf transcriptome. Nature Genetics 42: 1060–1067.

Liu Q, Teng S, Deng C, Wu S, Li H, Wang Y, Wu J, Cui X, Zhang Z, Quick WP, Brutnell TP, Sun X, Lu T. 2023. SHORT ROOT and INDETERMINATE DOMAIN family members govern PIN-FORMED expression to regulate minor vein differentiation in rice. Plant Cell doi: 10.1093/plcell/koad125.

Liu WY, Chang YM, Chen SC, Lu CH, Wu YH, Lu MY, Chen DR, Shih AC, Sheue CR, Huang HC, Yu CP, Lin HH, Shiu SH, Ku MS, Li WH. 2013. Anatomical and transcriptional dynamics of maize embryonic leaves during seed germination. Proc Natl Acad Sci U S A. 110(10):3979–84.

Liu WY, Lin HH, Yu CP, Chang CK, Chen HJ, Lin JJ, Lu MJ, Tu SL, Shiu SH, Wu SH, Ku MSB, Li WH. 2020. Maize ANT1 modulates vascular development, chloroplast development, photosynthesis, and plant growth. Proc Natl Acad Sci U S A. 117(35):21747–21756.

Liu WY, Yu CP, Chang CK, Chen HJ, Li MY, Chen YH, Shiu SH, Ku MSB, Tu SL, Lu MJ, Li WH. 2022. Regulators of early maize leaf development inferred from transcriptomes of laser capture microdissection (LCM)-isolated embryonic leaf cells. Proc Natl Acad Sci U S A 119(35): e2208795119.

Love MI, Huber W & Anders S. 2014. Moderated estimation of fold change and dispersion for RNA-seq data with DESeq2. Genome Biol 15, 550.

Lee RD, Munro SA, Knutson TP, LaRue RS, Heltemes-Harris LM, Farrar MA. Single-cell analysis identifies dynamic gene expression networks that govern B cell development and transformation. Nat Commun. 2021 Nov 25;12(1):6843.

Mallmann J, Heckmann D, Bräutigam A, Lercher MJ, Weber AP, Westhoff P, Gowik U. 2014. The role of photorespiration during the evolution of C4 photosynthesis in the genus Flaveria. Elife. 3:e02478.

Majeran W, van Wijk KJ. 2009. Cell-type-specific differentiation of chloroplasts in C4 plants. Trends in Plant Science 14(2): 100–109.

Nazipova A, Gorshkov O, Eneyskaya E, Petrova N, Kulminskaya A, Gorshkova T, Kozlova L. 2022. Forgotten Actors: Glycoside Hydrolases During Elongation Growth of Maize Primary Root. Front Plant Sci 12: 802424.

Ortiz-Ramírez C, Guillotin B, Xu X, Rahni R, Zhang S, Yan Z, Coqueiro Dias Araujo P, Demesa-Arevalo E, Lee L, Van Eck J, Gingeras TR, Jackson D, Gallagher KL, Birnbaum KD. 2021. Ground tissue circuitry regulates organ complexity in maize and Setaria. Science 374(6572): 1247–1252.

Perico C, Tan S, Langdale JA. 2022. Developmental regulation of leaf venation patterns: monocot versus eudicots and the role of auxin. New Phytologist 234:783–803.

Perico C, Zaidem M, Sedelnikova O, Bhattacharya S, Korfhage C, Langdale JA. 2024. Spatial transcriptomics reveals distinct lineage identities for major and minor vein initiation during maize leaf development. bioRxiv 2024.02.05.578898; doi: 10.1101/2024.02.05.578898

Pick TR, Brä utigam A, Schlüter U, Denton AK, Colmsee C, Scholz U, Fahnenstich H, Pieruschka R, Rascher U, Sonnewald U, Weber AP. 2011. Systems analysis of a maize leaf developmental gradient redefines the current C4 model and provides candidates for regulation. Plant Cell 23: 4208–4220.

Qiu X, Mao Q, Tang Y, et al. 2017. Reversed graph embedding resolves complex single-cell trajectories. Nat Methods 14(10): 979–982. doi:10.1038/nmeth.4402

Robil JM, McSteen P. 2023. Hormonal control of medial-lateral growth and vein formation in the maize leaf. New Phytol. 238(1):125–141.

Ran X, Zhao F, Wang Y, et al. 2020. Plant Regulomics: a data-driven interface for retrieving upstream regulators from plant multi-omics data. Plant J 101(1): 237–248.

Rohrmeier T, Lehle L. 1993. WIP1, a wound-inducible gene from maize with homology to Bowman-Birk proteinase inhibitors. Plant Mol Biol 22(5): 783–792.

Sharman BC. 1942. Developmental Anatomy of the Shoot of Zea mays L. Annals of Botany 6(2): 245–282.

Singh P, Stevenson SR, Reyna-Llorens I, Reeves G, Schreier TB, Hibberd JM. 2020. Upregulation and cell specificity of C4 genes are derived from ancestral C3 gene regulatory networks. bioRxiv 2020.07.03.186395.

Satterlee JW, Strable J, Scanlon MJ. 2020. Plant stem-cell organization and differentiation at single-cell resolution. Proc Natl Acad Sci U S A 117(52): 33689–33699.

Seyfferth C, Renema J, Wendrich JR, Eekhout T, Seurinck R, Vandamme N, Blob B, Saeys Y, Helariutta Y, Birnbaum KD, De Rybel B. 2021 Advances and Opportunities in Single-Cell Transcriptomics for Plant Research. Annu Rev Plant Biol. 72:847–866.

Tausta SL, Li P, Si Y, Gandotra N, Liu P, Sun Q, Brutnell TP, Nelson T. 2014. Developmental dynamics of Kranz cell transcriptional specificity in maize leaf reveals early onset of C4-related processes. J Exp Bot 65: 3543–3555.

Thibivilliers S, Anderson D, Libault M. 2020. Isolation of Plant Root Nuclei for Single Cell RNA Sequencing. Curr Protoc Plant Biol 5(4): e20120.

von Caemmerer S, Ghannoum O, Furbank RT. 2017. C4 photosynthesis: 50 years of discovery and innovation. J Exp Bot 68(2): 97–102.

van Campen JC, Yaapar MN, Narawatthana S, Lehmeier C, Wanchana S, Thakur V, Chater C, Kelly S, Rolfe SA, Quick WP, Fleming AJ. 2016. Combined Chlorophyll Fluorescence and Transcriptomic Analysis Identifies the P3/P4 Transition as a Key Stage in Rice Leaf Photosynthetic Development. Plant Physiol 170(3):1655–1674.

Vlad D, Zaidem M, Perico C, Sedelnikova O, Bhattacharya S, Langdale JA. 2023. The WIP6 transcription factor *TOO MANY LATERALS* specifies vein type in C4 and C3 grass leaves. bioRxiv 2023.12.20.572592; doi: 10.1101/2023.12.20.572592

Wang L, Czedik-Eysenberg A, Mertz RA, Si Y, Tohge T, Nunes-Nesi A, Arrivault S, Dedow LK, Bryant DW, Zhou W, Xu J, Weissmann S, Studer A, Li P, Zhang C, LaRue T, Shao Y, Ding Z, Sun Q, Patel RV, Turgeon R, Zhu X, Provart NJ, Mockler TC, Fernie AR, Stitt M, Liu P, Brutnell TP. 2014. Comparative analyses of C4 and C3 photosynthesis in developing leaves of maize and rice. Nature Biotechnology 32: 1158–1165.

Wang P, Kelly S, Fouracre JP, Langdale JA. 2013. Genome-wide transcript analysis of early maize leaf development reveals gene cohorts associated with the differentiation of C4 Kranz anatomy. Plant Journal 75(4): 656–670.

Wang P, Vlad D, Langdale JA. 2016. Finding the genes to build C4 rice. Current Opinion in Plant Biology 31: 44–50.

Weigel, D. A. G. J. 2022. Arabidopsis: A Laboratory Manual (Cold Spring Harbor Laboratory Press).

Young MD, Behjati S. 2020. SoupX removes ambient RNA contamination from droplet-based single-cell RNA sequencing data. Gigascience 9(12):giaa151. doi:10.1093/gigascience/giaa151

Xu X, Crow M, Rice BR, Li F, Harris B, Liu L, Demesa-Arevalo E, Lu Z, Wang L, Fox N, Wang X, Drenkow J, Luo A, Char SN, Yang B, Sylvester AW, Gingeras TR, Schmitz RJ, Ware D, Lipka AE, Gillis J, Jackson D. 2021. Single-cell RNA sequencing of developing maize ears facilitates functional analysis and trait candidate gene discovery. Dev Cell 56(4): 557–568.e6.

Xia K, Sun HX, Li J, Li J, Zhao Y, Chen L, Qin C, Chen R, Chen Z, Liu G, Yin R, Mu B, Wang X, Xu M, Li X, Yuan P, Qiao Y, Hao S, Wang J, Xie Q, Xu J, Liu S, Li Y, Chen A, Liu L, Yin Y, Yang H, Wang J, Gu Y, Xu X. 2022. The single-cell stereo-seq reveals region-specific cell subtypes and transcriptome profiling in Arabidopsis leaves. Dev Cell. 57(10):1299–1310.e4.

Yang F, Bui HT, Pautler M, Llaca V, Johnston R, Lee BH, Kolbe A, Sakai H, Jackson D. 2015. A maize glutaredoxin gene, abphyl2, regulates shoot meristem size and phyllotaxy. Plant Cell 27(1): 121–131.

Yu NI, Lee SA, Lee MH, Heo JO, Chang KS & Lim J. 2010. Characterization of SHORT-ROOT function in the Arabidopsis root vascular system. Molecules and cells 30, 113–119.

Zeng J, Li X, Ge Q. et al. 2021. Endogenous stress-related signal directs shoot stem cell fate in Arabidopsis thaliana. Nat Plants 7, 1276–1287.

Zhong W, Zheng C, Dong L et al. 2023. The maize callose synthase SLM1 is critical for a normal growth by controlling the vascular development. Mol Breeding 43, 2.

