## Supplementary figures and images for "Single-cell resolved differentiation of pre-Kranz anatomy in maize leaf primordia"

### Supplemental Dataset 9

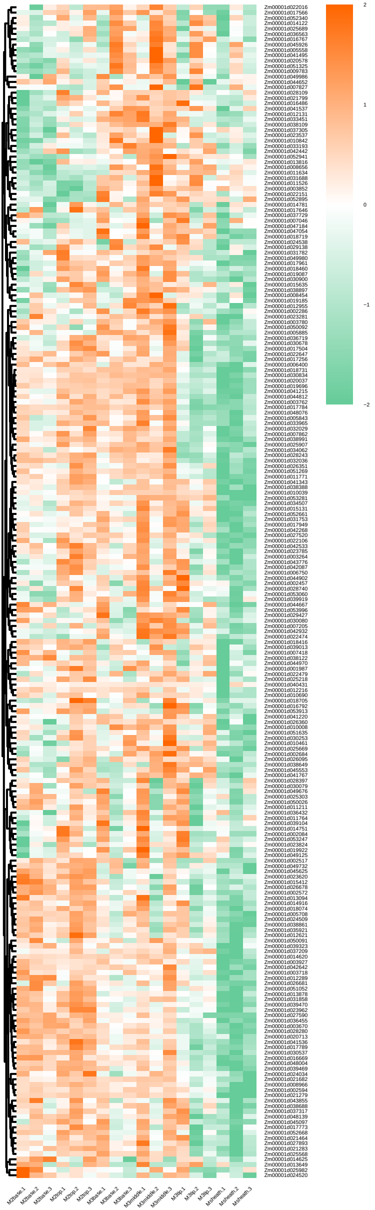
